# Syntheses of Pyrimidine-Modified Seleno-DNAs as Stable Antisense Molecules

**DOI:** 10.1101/2023.05.02.539140

**Authors:** Ziyuan Fang, Yuliya Dantsu, Cen Chen, Wen Zhang, Zhen Huang

## Abstract

Chemically modified antisense oligonucleotides (ASO) currently in pre-clinical and clinical experiments mainly focus on the 2′-position derivatizations to enhance stability and targeting affinity. Considering the possible incompatibility of 2′-modifications with RNase H stimulation and activity, we have hypothesized that the atom specific modifications on nucleobases can retain the complex structure and RNase H activity, while enhancing ASO’s binding affinity, specificity, and stability against nucleases. Herein we report a novel strategy to explore our hypothesis by synthesizing the deoxynucleoside phosphoramidite building block with the seleno-modification at 5-position of thymidine, as well as its Se-oligonucleotides. Via X-ray crystal structural study, we found that the Se-modification was located in the major groove of nucleic acid duplex and didn’t cause the thermal and structural perturbations. Surprisingly, our nucleobase-modified Se-DNAs were exceptionally resistant to nuclease digestion, while compatible with RNase H activity. This affords a novel avenue for potential antisense modification in the form of Se-antisense oligonucleotides (Se-ASO).

## INTRODUCTION

The first idea of using antisense oligonucleotides (ASO) technology to control expression of disease genes occurred in 1970s^1^. In the past decades, there have been steady progress in modifying the ASO platform as optimal therapeutics, including development of nucleic acid chemistry, delivery straegy of ASOs and more advanced safety and distribution studies^2-5^. There are several approved and marketed ASO drugs^6-8^, and numerous candidates are in a broad pipeline in development. The ASO technology is conceptually simple: using a short DNA oligonucleotide (13-25 nucleotides) to hybridize to a specific mRNA sequence, which blocks the regular flowing of genetic information from DNA to protein^9^. However, although ASOs have been commonly in use in clinic and laboratory, many uncertain questions cannot be belied concerning the activity of ASOs, like difficult delivery, off-target effect, target accessibility and susceptibility to nucleases in vivo^10,11^.

Due to the presence of nucleases (DNases and RNases) and their rapid attacking ASOs in biological fluid^12^, stability of potential oligonucleotides against endogenous nucleases is a prerequisite property for ASO therapeutics^13^. To increase the enzymatic stability, ASOs are regularly heavily chemically modified to prolong the circulation time in human body, enhance the targeting affinity, and reduce the non-specific cytotoxicity^14^. The most canonical modifications are focused in altering the structures of sugar and backbone^15,16^, including phosphorthioates^17,18^, 2′-O-methyl ASOs^19^, 2′-O-methoxy-ethyl ASOs^20^, borano-phosphates^21,22^, peptide nucleic acids (PNAs^23,24^), locked nucleic acids (LNAs^25^), morpholino nucleic acids^26^, and many others. The development of these novel chemical modifications in ASOs has provided significant breakthrough in this filed and paved the road for the potential therapeutic applications.

Generally, ASO technology includes two strategies: the RNase H-dependent approach and the RNase H-independent approach (RNA arresting)^27-29^. In RNase H-independent strategy, the modified antisense DNA binds to target mRNA through Watson-Crick base pairs. Formation of the RNA/DNA duplex exerts a direct steric effect to block the assembly of ribosomal subunits to initiate the translation, resulting in protein expression termination. In this way, the translation was arrested, and the target gene could be downregulated. Furthermore, antisense molecules can help prevent undesired mRNA splicing^30^. In the RNase H-dependent approach, the hybridized ASO/mRNA duplex is recognized by RNase H and mRNA is cleaved, leading to gene expression inhibition and release of ASO to further circulate^31^. In theory, RNase H-dependent ASO can lead to more profound gene silencing effect, considering the ubiquitous existence of RNase H in cytoplasm and nucleus^32^. Therefore, it is ideal to retain the compatibility to RNase H when designing ASO therapies.

RNase H enzymes uniquely recognize RNA/DNA hybrid geometry via numerous contacts with functional groups on both RNA and ASO^33^. Many modifications to the ASO therapies, especially to the 2′-position of deoxyribose, can distort the DNA geometry and prevent cleavage of the complementary RNA strand^34^. The classic phosphorothioate (PS)-modification can resist the nuclease degradation while maintaining the 2’-endo conformation in ASO, however, the thiol-modifications can cause non-specific binding of ASO to cellular proteins^35^ and introduce the thermal instability to the DNA/RNA duplex^36^. Therefore, it is important to keep exploring novel chemical modifications for ASO therapies, which contribute to three aspects of ASO features: (1) enhanced enzymatic stability in vitro and in vivo; (2) optimal binding affinity to target RNA via sequence complementarity; and (3) preserved conformations of deoxyribose and DNA/RNA hybrid, as well as the RNase H activity.

Atom-specific modifications on non-Watson-Crick edge of nucleobases are promising candidates, because the significant structure perturbation can be avoided by rotating the glycosidic bond to accommodate the modifications in the major or minor grooves^37,38^. Plus, the base pairs are maintained to form the DNA/RNA duplex as substrate of RNase H. A few instances of nucleobase-modified ASO have been reported^39-41^, including 2-thiouridine, 5-methyl-cytidine and 2′-deoxyl-7-deaza-guanosine, which have higher duplex stability by enhancing the base stacking interactions^42^. Particularly, various substituents have been tethered at the 5-position of pyrimidine^43-45^, and these modified DNA/DNA or DNA/RNA duplexes present enhanced thermal stability than their natural counterparts^46^.

Selenium element is in the same family of oxygen in the periodic table, with the enlarged atomic radius. It has been revealed that the replacement of O with Se on the nucleobases provides nucleic acid molecules innovative features, like improving base-paring specificity, duplex stability, complementary recognition and fidelity in enzymatic replication^47^. This modification leads to Se-mediated hydrogen bond and stacking interactions, and the Se-derivatization can be tolerated by DNA duplex without structural perturbation^48,49^. In addition, the selenium derivatization in nucleic acid and nucleic acid-protein complex can facilitate X-ray crystal structure studies via multiwavelength anomalous dispersion (MAD) phasing^50,51^. Therefore, the selenium substitution on nucleobase likely provides a unique opportunity in design of novel ASO therapy.

The previous results have demonstrated that, when DNA molecule containing 6-Se-dG modification hybridizes to RNA target, it can engender the RNA cleavage by RNase H, with an enhanced activity^52^. The facilitation is probably due to the local unwinding of the substrate RNA/DNA duplex. However, the replacement of oxygen with selenium on 6-position of Guanosine is quickly oxidized in physiological condition, making us keep searching for new selenol-modification stable and effective enough in ASO therapeutics. Based on our previous research, 5-methylseleno-thymidine is the most stable Se-atom-specific modification on nucleobase^53^. The selenium atom located in the major groove and did not perturb the base pairing recognition or sugar pucker conformation. Thus, we hypothesize that, the 5-Se-pyrimidine DNA strands hold promise as a novel method for antisense modification, presenting a potential avenue for nucleic acid drug development.

## RESULTS and DISCUSSION

### Chemical Synthesis

The 5-Methylselenol-thymidine nucleoside and DNA have been successfully synthesized via the Mn(OAc)_3_ assisted electrophilic addition of CH_3_SeSeCH_3_^53^. However, the reaction yield was moderate (∼56%), and the reaction had to be carried out in acetic acid at 90°C, which potentially caused the nucleoside damage. Therefore, we tried to optimize the reaction conditions, and report here an optimized method to incorporate methylselenol modification (and other chalcogen modifications) to the 5-position of thymidine in higher yield (Scheme 1).

**Scheme 1.**
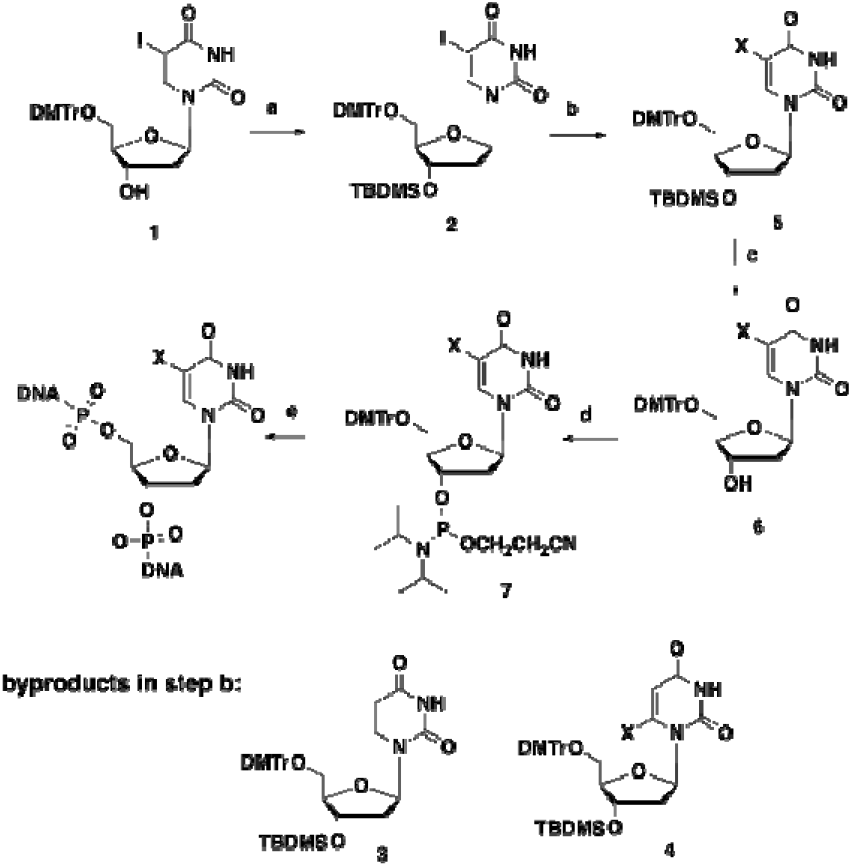
Synthesis of 5-chalcogen-T DNAs. Reagents and conditions: a) TBDMS-Cl, Im., DMF, 4 h, rt, 92%; b) i) NaH, THF, 15 min, r.t., ii) lithiating agent (table 1), THF, 30 min., −78 °C, iii) X_2_, 1 h, −78 °C; c) TBAF, THF, 4 h, rt, 92%; d) i-Pr_2_ NP(Cl)CH_2_ CH_2_ CN, DIEA, CH_2_ Cl_2_, 1 h, rt, 82%; e) solid phase synthesis. X=CH_3_Se, CH_3_S or PhSe.

The synthesis started from 5-iodo-5′-DMTr-thymidine (Scheme 1, 1). After the protection of 3′-hydroxyl group by TBDMS, methylselenol functionality was introduced into 5-position by replacing the iodide substitution. To introduce non-carbon substituents at the 5-position of pyrimidine, there are reported approaches including electrophilic addition-elimination to the C5-C6 double bond^54^, palladium catalyzed nucleophilic substitution of 5-mercururio derivatives^46^, and lithiation of 5-halo derivatives in the presence of the electrophilic species^55^. The milder reaction condition of the later procedure led us to examine its versatility for the proposed 5-methylselenyl-thymidine derivative.

Lithiation of suitably protected uridine^56,57^ and thymidine^39,55,58^ in the presence of electrophile has been reported for the synthesis of variety of 5-substituted pyrimidine nucleosides. However, two issues remained to be concerned, including the formation of the reduction byproduct, the uridine/ thymidine derivatives, and the regioselectivity between C-5 and C-6 positions of the pyrimidine. Generally, when 2′-OH is present as in uridine, a high C-5/C-6 regioselectivity is observed due in part to the steric effects of 2′-position^57^. However, when the nucleoside is lacking 2′-OH group, lithium/halogen exchange produces a relatively low C-5 regioselectivity^58^. In that case, the N^3^-imido moiety can be protected to eliminate the competition for the lithiation agent between N^3^-H and C-5, yielding higher C5 lithiation yield^55,59^ .Therefore, we envision that applying a transient N^3^-imido moiety protection of 5-iodo-thymidine derivative (2) followed by lithium-halogen exchange and quenching with (CH_3_)_2_Se_2_ would provide an efficient access to the proposed nucleoside (5). As the first attempt, treatment of compound 2 with NaH (1.5 equiv), n-BuLi (2.2 equiv) and excess of (CH_3_)_2_Se_2_ resulted in an inseparable mixture of 5-methylselenyl derivative (5) and the corresponding 6-methylselenyl derivative (4) in 7:1 ratio (entry 1, Table 1), along with the reduced by-product derivative (3).

**Table 1.** Transient N^3^ protection and lithium/halogen exchange of compound 2 in the presence of dimethyldiselenide.^a^

| entry | NaH<sup>b</sup><br>(eq.) | lithiating<br>agent<sup>c</sup>(eq.) | concentration (M) | ratio<sup>d</sup><br>5:4 | yield<sup>e</sup> %<br>5+4 | 3 |
| --- | --- | --- | --- | --- | --- | --- |
| 1 | 1.5 | *n*-BuLi (2.2) | 0.1 | 7:1 | 71 | 20 |
| 2 | 1.5 | *sec*-BuLi (2.2) | 0.1 | 11:1 | 68 | 23 |
| 3 | 1.5 | *tert*-BuLi (2.2) | 0.1 | 10:1 | 65 | 21 |
| 4 | 1.5 | LiHMDS (2.2) | 0.1 | -f | - | - |
| 5 | 1.5 | *n*-BuLi (2.2) | 0.15 | 35:1 | 75 | 20 |
| 6 | 1.5 | *n*-BuLi (2.2) | 0.2 | only 5 | 71 | 19 |
| 7 | 1.05 | *n*-BuLi (2.2) | 0.2 | only 5 | 85 | 5 |

a) All reactions were performed in the presence of 4 equivalents of (CH<sub>3</sub>)<sub>2</sub>Se<sub>2</sub>; b) Deprotonation of the N<sup>3</sup>-H was performed at ambient temperature for 30 min; c) Lithium/halogen exchange was performed at -78 °C for 30 min. d) The ratio was determined by the integration of the H-1' of the <sup>1</sup>H-NMR of the mixture. e) Isolated yields after silica gel chromatography. f) The starting material was recovered.

We further screened various conditions to optimize the C5/C6 regioselectivity and reaction yield. Lithiation using sec-BuLi or tert-BuLi gave similar results to n-BuLi in terms of C-5/C-6 regioselectivity (entries 2 and 3, Table 1). We also applied a bulkier lithiating agent, LiHMDS, in attempt to improve the C5/C6 regioselectivity, however, no corresponding products were observed, probably due to steric hindrance (entry 4). We next examined the effect of the reaction concentration on the C-5/C-6 regioselectivity. At higher concentration (0.15 M and 0.2 M) with n-BuLi as the lithiating agent, the methylselenylation occurred exclusively at the C-5 position, yet the reduced byproduct 3 was isolated in about 20% yield (entries 5 and 6). Apparently, at higher concentration, the rate of lithium/halogen exchange at the 5-position is faster, discouraging the formation of the 6-lithio intermediate and thus results in the observed high regioselectivity. We also found that decreasing the molar equivalence of NaH (1.05 equiv) substantially eliminates the formation of the reduced product 3 (entry 7). In the following steps, deprotection of the 3’-TBDMS group and the phosphitylation reaction were carried out under standard conditions, yielding the building block 7 for solid phase synthesis.

To make the comprehensive comparison, we also produced the thymidine derivatives containing different chalcogen modifications at 5-position, including Methylthiol-, Phenylselenol- and Methoxy-moieties. When synthesizing the 5-CH_3_S-T and 5-PhSe-T nucleosides, the same reaction conditions were followed as 5-MeSe-T synthesis. 5-CH_3_O-T derivative was synthesized using different strategy (Scheme S2). 5-Hydroxy-thymidine was methylated at room temperature by methyliodide, yielding the corresponding product. Four different thymidine phosphoramidites containing 5-chalcogen-modifications were successfully synthesized and incorporated into oligonucleotides by solid-phase synthesis. The detailed operations and results are discussed in experimental section and SI.

### Synthesis of 5-chalcogen-T modified DNAs

We then attempted the synthesis of these 5-chalcogen-T modified DNAs using regular phosphoramidites and 5-(benzylmercapto)-^1^H-tetrazole as the coupling catalyst. The coupling of modified phosphoramidites was performed for 25s, and the coupling yields of our synthetic phosphoramidites were over 95%. After solid-phase synthesis, ammonium (conc.) and 3% trichloroacetic acid were used to deprotect the DNAs and remove the 5’-DMTr groups. The modified oligonucleotides were purified twice with DMTr-on and DMTr-off. The HPLC spectra and mass spectroscopy analyses indicated the purity and integrated chalcogen modifications in DNAs (SI and Table S1). In addition, it was found that all the chalcogen functionalities were compatible to conditions used for regular DNA synthesis, including strong acid treatment, strong base treatment at elevated temperature, and I_2_ oxidation.

### Thermo-stability of 5-chalcogen-modified DNAs

The binding affinity to the complementary oligonucleotide is one critical feature of ASO in clinical application. In consequence, we performed the UV-melting temperature assay to assess whether the 5-chalcogen modifications impact the thermodynamic stability of the nucleic acid duplexes. 5-MeSe-modification in thymidine was reported to be thermodynamically stable in DNA duplex^53^. We expect that, similarly, the other chalcogen modifications don’t cause significant lost in duplex thermostability.

Melting experiments in three different DNA duplexes were tested. The melting temperatures are summarized in Table 2. It indicates that CH_3_Se-, CH_3_S- and CH_3_O-derivatized DNAs have almost the same thermostability as native DNAs. That is likely because the 5-chalcogen-modifications are tolerated by the major grooves of duplexes, and the electron delocalization ability of chalcogen elements contributes partially to the stacking interactions between base pairs. However, the melting temperatures of 5-PhSe-DNAs are roughly 4-6 °C lower than native DNAs, which is probably caused by the steric effect of bulky phenyl group, which produces structural disorder to the duplex. Principally, when DNAs are modified by CH_3_Se-, CH_3_S- and CH_3_O-groups, the binding affinities are acceptable as potential ASO drugs.

**Table 2.** UV melting temperatures of the 5-chalcogen-T DNAs.

| entry | Duplex System | X base | Melting Temp (°C) |
| --- | --- | --- | --- |
| *a* | 5'-ATGGXGCTC-3'<br>3'-TACCACGAG-5' | T | 40.3 ± 0.6 |
| *b* |  | 5-MeSe-T | 39.3 ± 0.8 |
| *c* |  | 5-MeS-T | 39.0 ± 0.9 |
| *d* |  | 5-MeO-T | 37.8 ± 0.6 |
| *e* |  | 5-PhSe-T | 34.4 ± 1.2 |
| *f* | 5'-CTCCCA<sup>X</sup>CC-3'<br>3'-GAGGGTAGG-5' | T | 36.5 ± 0.7 |
| *g* |  | 5-MeSe-T | 36.3 ± 0.7 |
| *h* |  | 5-MeS-T | 36.8 ± 1.0 |
| *i* |  | 5-MeO-T | 36.9 ± 0.6 |
| *j* |  | 5-PhSe-T | 32.3 ± 1.0 |
| *k* | 5'-CTTCT<sup>X</sup>GTCCG-3'<br>3'-GAAGAACAGGC-5' | T | 44.9 ± 0.8 |
| *l* |  | 5-MeSe-T | 43.9 ± 0.7 |
| *m* |  | 5-MeS-T | 43.6 ± 0.8 |
| *n* |  | 5-MeO-T | 43.5 ± 0.9 |
| *o* |  | 5-PhSe-T | 39.6 ± 1.1 |

### X-ray crystal structures of 5-Chalcogen-T modified DNAs

We also carried out the structural study of the self-complementary 5-chalcogen-T-DNAs (5′-G-dU_2_’-Se-GXACAC-3′), facilitated by 2′-MeSe-dU to crystallize. The oligonucleotides with 5-CH_3_O-, 5-CH_3_S- and 5-CH_3_Se-modifications were crystallized and the structures were determined successfully. Our structural information indicated these modified DNAs had almost the same crystal structures as native one, and the modified thymidines could form Watson-Crick base pairs with adenosines^60^. Unfortunately, the 5-PhSe-T DNA didn’t crystallize as expected. This result is probably due to the perturbation PhSe functionality brought in, and matches the results of thermostability studies. The 5-chalcogen-modifications don’t result in observable structural instability to DNA duplexes. The crystal structures of modified DNAs are shown in Figure 1.

**Fig. 1.**
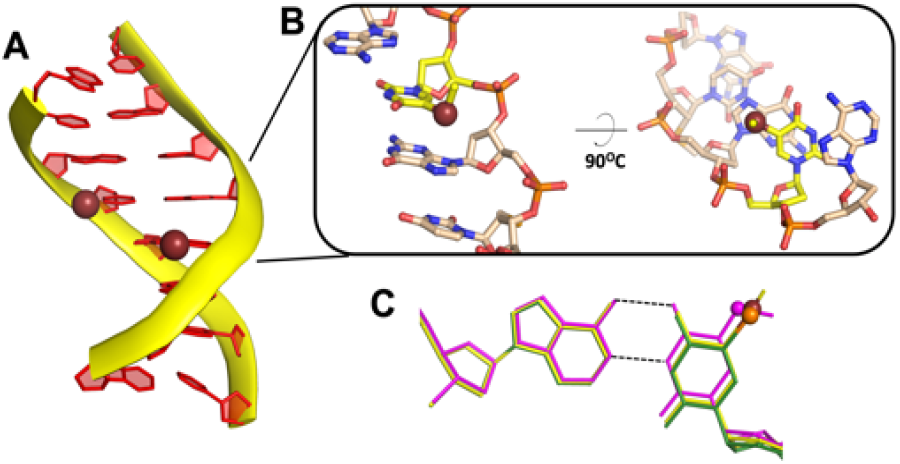
Crystal structures of 5-chalcogen-modified DNAs. (A) Structure of DNA 8mer containing 5-MeSe-modifications. Ruby spheres: Se atoms. (B) Local structures of 5-MeSe-thymidine and the stacking nucleobases. (C) Superimposed structures of base pairs. Purple: 5-MeO-T:A; green: 5-MeS-T:A; yellow: 5-MeSe-T:A. Ruby sphere: Se; Orange sphere: sulfur; purple sphere: oxygen.

### Enzymatic stability assay of 5-Chalcogen-T modified DNAs

In ASO application, DNA molecules are expected to avoid the rapid digestion by intracellular enzymes-mediated hydrolysis, such as endo- and exonucleases. Accordingly, we decide to incubate the 5-Chalcogen-T modified DNAs in endo- and exonucleases solutions, and evaluate their stability against hydrolytic metabolism. Currently, there have been over 3000 types of restriction endonucleases isolated from bacteria and studied in laboratory^61^. The recognition site for each restriction endonuclease is variable, usually 4-8 nucleotides long, and mostly palindromic. Here we choose two kinds of type II restriction endonucleases, AseI and SalI, to study how 5-Chalcogen-T residue affects the cleavage efficiency. Both endonucleases have the cleavage sites close to thymidine, and they require magnesium as cofactor in the hydrolysis^62^. The designed ASO sequences containing different modifications are listed in Table 3. The DNAs we tested are 24mer oligonucleotides containing different modified residues, used to test enzymatic stability against endonucleases, exonuclease and serum. The complementary DNA is also synthesized containing all native residues. The native and modified oligonucleotides were 5′-^32^P labeled using T4 polynucleotide kinase (New England Biolabs) following manufactures recommendations, and annealed with the complementary strand. The DNA duplexes were treated with different types of nucleases, and the reaction products were analyzed by PAGE gel electrophoresis after quenched by loading dye. The gels were dried and exposed to film for analysis. Imaging and quantification of the reaction substrate and products were performed with an Amersham Biosciences Typhoon 9410 Phosphorimager, and the ImageJ software^63^. Gels depicted in Figures are representative of triplicate experiments.

**Table 3.**
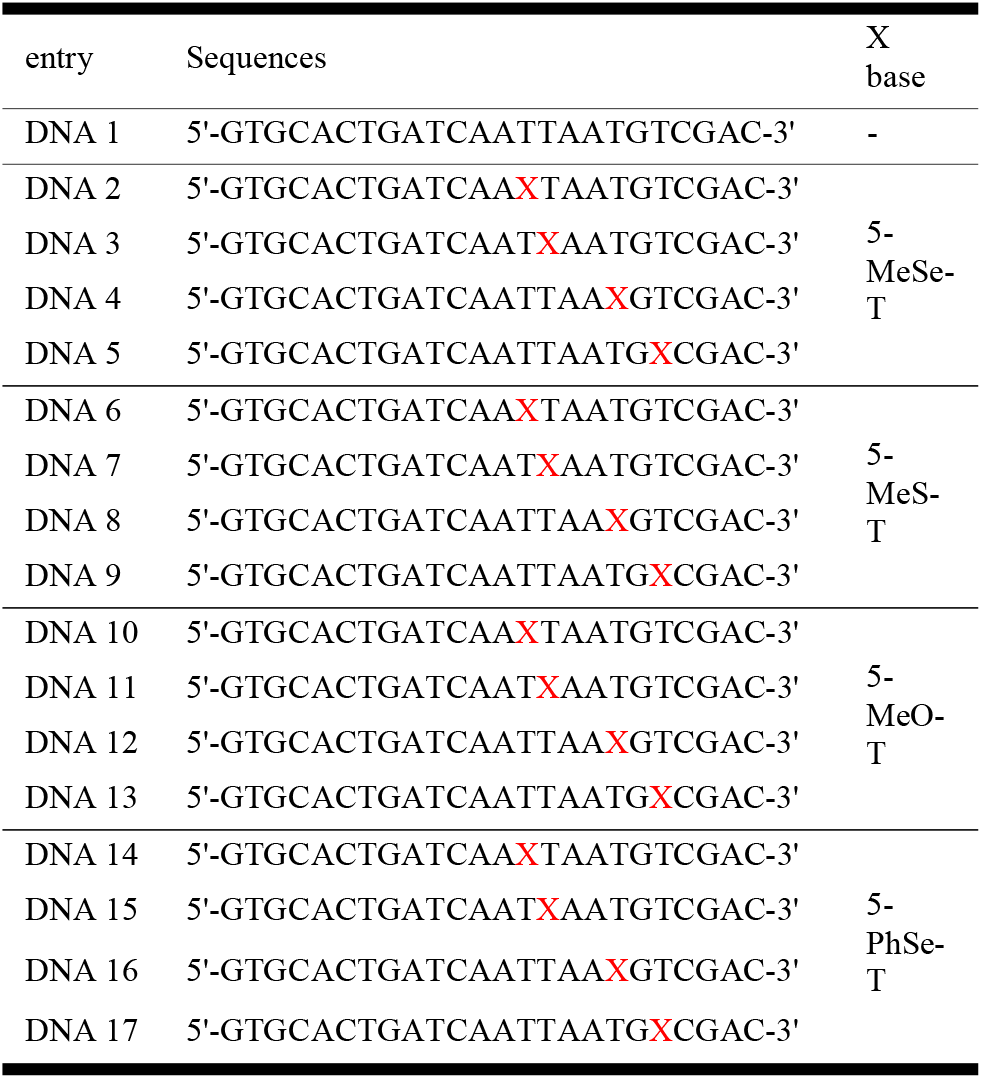
Sequences of 5-chalcogen-T DNAs as ASOs.

We first examine the capacity of 5-MeSe-modification to resist AseI, and its relationship with the location of modification (Figure 2). Originated from Aquaspirillum serpens, AseI specially recognizes the sequence 5′-ATTAAT-3′, and the cleavage site is located between the two consecutive thymidines (Figure 2A). DNA 1-5 were incubated with AseI, aliquots were taken at different time points, and the reaction was quenched for hydrolysis analysis. We observe that, without any modification, the wild-type DNA 1 was completely digested within 16 hours. Interestingly, when DNAs were modified by 5-MeSe-groups inside the recognition sequence (DNA 2-4), there was dramatic resistance to AseI hydrolysis (Figure 2B-E). While when the modification is distant from the cleavage site, like in DNA 5, the similar hydrolysis efficiency was observed as native DNA (Figure 2F). The time-dependent plot was made to compare the hydrolysis rates of different DNAs against AseI in 16 h reaction time (Figure 2G). It reveals that the 5-MeSe-T residue has inhibitory effect on endonuclease digestion, especially when the modification is close to the cleavage site. The possible reason could be that the bulkiness and rich electron of selenium atom discourage the binding of DNA to the enzyme, or Se causes the distortion of conformations of catalytic residues in AseI, leading to the loss of hydrolysis function.

**Fig. 2.**
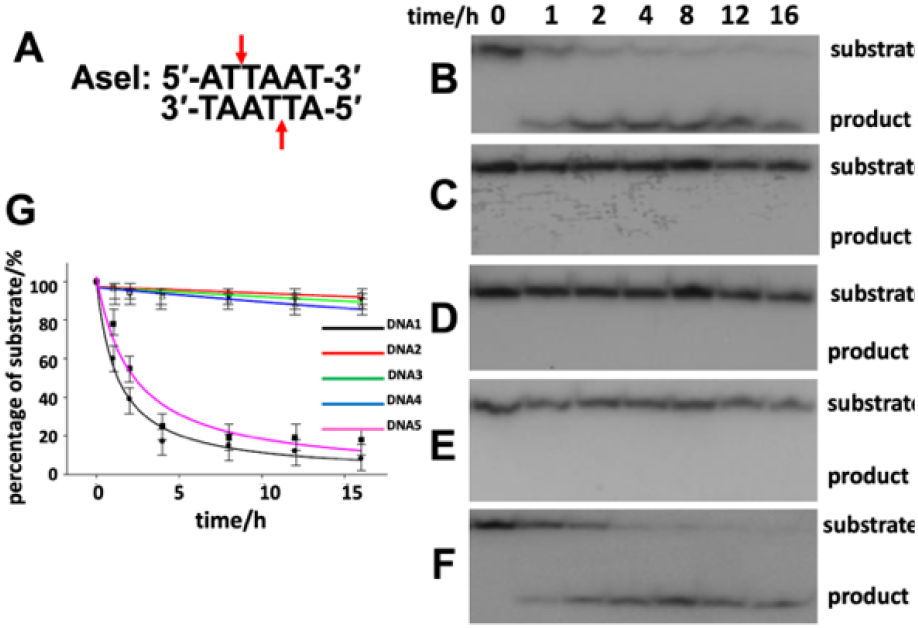
Nuclease resistance and stabilities of 5-CH_3_Se-T DNAs to endonuclease AseI. (A) Recognition and cleavage site of Ase I; (B-F) Time-dependent cleavage of DNA1-5 over 16 hours; (G) Plot of the ratio of DNA unhydrolyzed over time.

We then compared the different chalcogen modifications, on their resistant impacts against AseI. Different ASO strands (DNA 1-17) were incubated with AseI enzyme for 16 hours, and the reactions were quenched for gel analysis (Figure 3). As demonstrated, native DNA 1 was almost completely digested in each individual incubation, and 5-CH_3_Se-modification led to striking suppression of the digestion, especially when the selenium element is close to the cleavage site (Figure 3A). Nonetheless, when other chalcogen modifications are present, the inhibitory effects become diverse accordingly. In the case of 5-MeS-modification, only DNA 6 and 7 exhibited resistance to AseI, and DNA 8 and 9 could be digested to a great extent (Figure 3B). When the modification became the smaller O atom, only DNA 10 presented a weaker resistance, while DNA 11, 12 and 13 behaved almost identically to native DNA (Figure 3C). These results indicated that, the inhibitory effect caused by chalcogen modifications in DNA are related to the electronic and steric properties of the elements, and CH_3_Se-modification generates more optimal enzymatic stability than others. Additionally, we surprisingly discovered that, the 5-PhSe modification, though it had the selenium element and bulky phenyl group, had weaker stability than 5-CH_3_Se group (Figure 3D). When the 5-PhSe modifications are beside the cleavage site in DNA 14 and 15, we could observe the significant resistant effects. But DNA 16 and 17 showed very weak resistance, which makes the overall effect of PhSe group almost identical to CH_3_S group. Considering that PhSe group generated instability to the DNA thermo- and structural features as discussed above, we speculated the bulky phenyl group could locally destabilize and unwind the DNA duplex, which somehow facilitates the formation of the catalytic center of AseI. Meanwhile, selenium atom has the unusual suppressive effect to the nuclease by resisting the binding, and the combinatorial result was that PhSe group induced inhibition, but not as strong as smaller CH_3_Se group.

**Figure 3.**
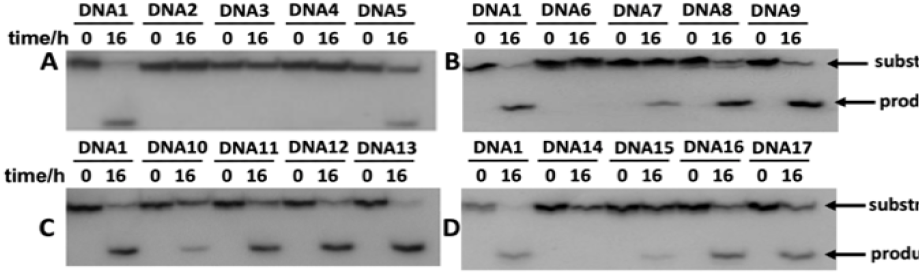
Nuclease resistance and stabilities of 5-chalcogen-T DNAs to endonuclease AseI. (A) X=5-CH_3_Se-T; (B) X=5-CH_3_S-T; (C) X=5-CH_3_O-T; (D) X=5-PhSe-T.

The resistant capacities of 5-Chalcogen-modified DNAs are confirmed by the stability assays using other nucleases, including SalI endonuclease, exonuclease III and snake venom phosphodiesterase I (PDE I). In brief, chalcogen modifications were able to enhance the stability of chalcogen-ASOs, and CH_3_Se group has the most potent effect against the digestions. Similar to our observation in AseI treatment, the efficiencies of resistance were related to the positions of modification. The results proved the potential prospective of 5-CH_3_Se modification in antisense drug design. The experimental results are listed in supporting information.

### Stabilities of 5-methylselenyl-T DNAs against serum

Serum is the important component added in cell culture to facilitate cell growing and maintaining. Serum is a complex mix of enzymes, growth factors and growth inhibitors, that is closer to the physiological conditions in human body. Therefore, we decide to appraise whether the selenium-modified DNAs, as the potential ASO therapeutics, have extended survival time when treated with serum. We incubated DNA 1-5 with 4% fetal bovine serum (FBS, life technology), and analyzed the degradation products by gel electrophoresis. The results indicate that, DNA molecule has stronger resistance against serum after modified by 5-methylselenyl group. After incubation for 60 min, ∼60% of DNA 5 remained intact, in contrast to that only 20% of native DNA 1 persisted. Among the 5-methylselenyl-DNAs, DNA 5 had the strongest resistance, while DNA 2 behaved in the same manner as native DNA. Apparently, the inhibitory effect is related to the position of modification. The Se-modification close to the 3′-end could effectively protect the DNA from enzymatic degradation, likely caused by the 3′-exonucleases in serum. The experimental results are listed in supporting information.

### Impact of 5-CH_3_Se-T on RNase H cleavage

RNase H is able to destroy the specific RNA when complementary DNA is hybridized, therefore, the RNase H mediated degradation of target mRNA is the effective mechanism of ASO therapies^64^. After demonstrating that 5-CH_3_Se modification significantly improves the stability of ASO in enzymatic environment, we then attempt to explore whether it influences the necessary RNase H digestion.

We designed and synthesized RNA c (5’-GUCGACAUUAA U-3’), which formed DNA/RNA duplexes with the synthetic DNAs. Since 5-CH_3_Se-T had the strongest resistance to nuclease digestion, we selected different 5-CH_3_Se-T-DNAs (DNA 1-5) to anneal with RNA c. The Se-modifications are all included in the region of hybridized duplex. After incubating with RNase H, the 5’-^32^P labeled RNA fragments were analyzed by gel. The result indicates that, when hybridizing to native DNA 1, the RNA molecule was completely fragmented within 15 min by RNase H (Figure 4A). Delightfully, we discovered that, even when the target RNA bound to the complementary Se-DNAs, there was no reduction on the activity of RNase H. The degradations of RNAs in all DNA/RNA hybrids were equally efficient (Figure 4B-E). The 5-CH_3_Se functionality can not only offer DNA excellent stability by resulting to better resistance to nucleases, but also retain adequate activity of RNase H to destruct target RNA. It displays great potential as promising ASO drugs.

**Fig. 4.**
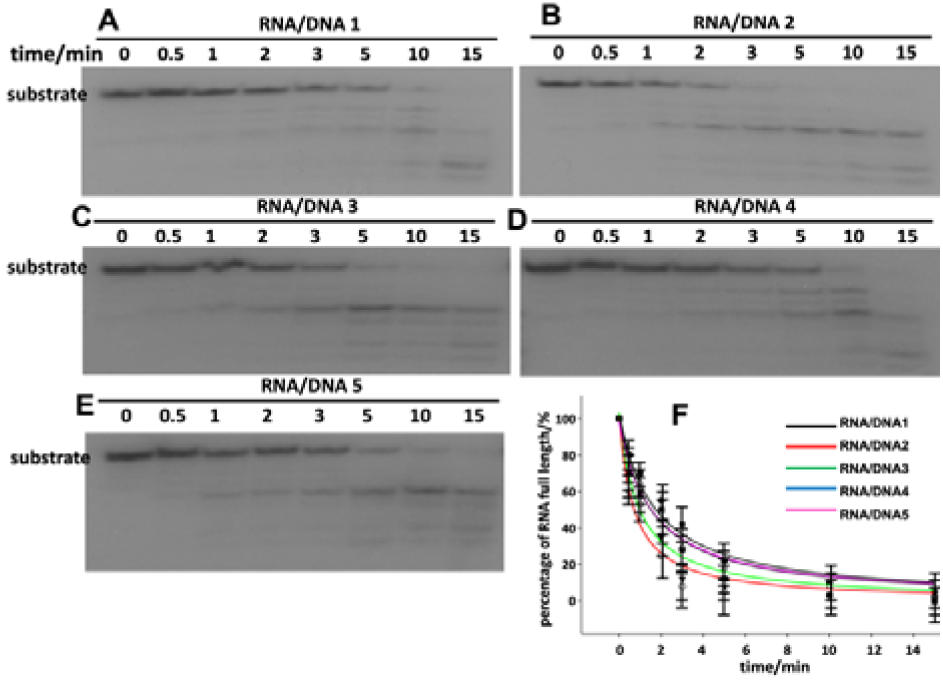
RNase H digestion of RNA after binding to different DNAs. The complementary DNAs are (A) native DNA 1; (B) DNA 2; (C) DNA 3; (D) DNA 4; (E) DNA5; (F) Plot of the ratio of RNA unhydrolyzed over time.

### X-ray crystal structure of Se-DNA/RNA/RNase H complex

As the innovative ASO candidate, the synthetic Se-DNAs are compatible to the activity of RNase H. We then attempt to investigate the structural insights into how RNase H recognizes Se-DNA/RNA hybrid and whether selenium modification brings in different mechanism of protein recognition. The RNase H enzyme from Bacillus halodurans (Bh)^52,65^ was successfully expressed and crystallized, complexed with both native DNA/RNA and Se-DNA/RNA hybrids. The Bh RNase H (196 AA) contains a short linker (10 aa) between the RNA/DNA hybrid binding domain and RNase H domain. The binding nucleic acid sequences in crystals are designed as DNA (5′-AXGXCG-3′) and RNA (5′-UCGACA-3′), where 5-methylselenyl-T or native T is incorporated to replace X accordingly. The DNA and RNA molecules form duplex with one nucleotide overhang, which base pairs with the nucleotide overhang of the neighboring complex to facilitate crystallization. We crystallized the C-terminal Bh-RNase H domain (RNase HC, 59-196 aa), and the structures were determined with molecular replacement using 2G8U as searching model. The refined model consists of amino acid residues 62-193 (Figure 5A). Crystallization conditions and crystallographic statistics are listed in SI.

**Fig. 5.**
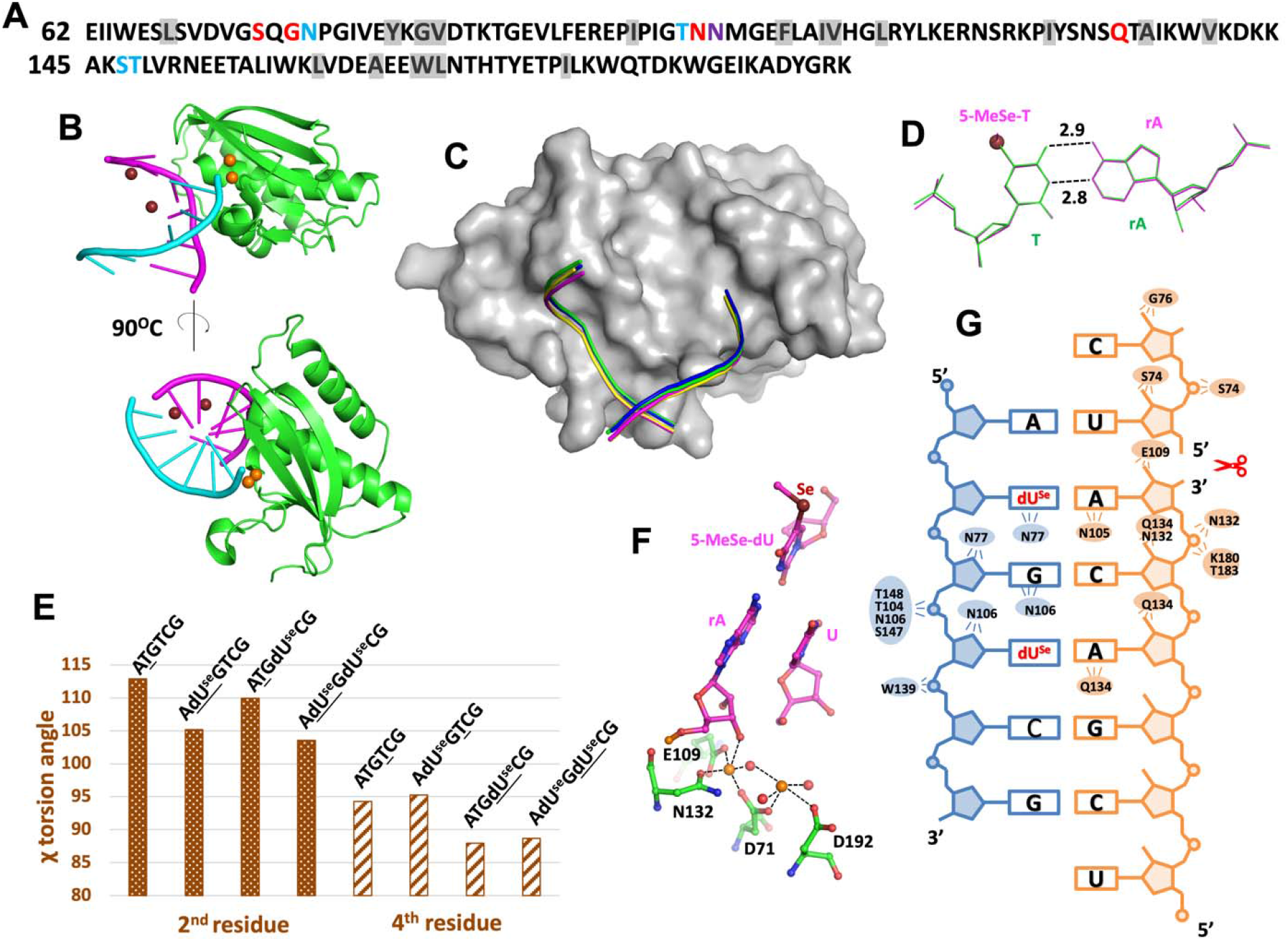
Structure of Bh RNase H. (A) Sequence of RNase H. Residues contacting the RNA strand are labeled in red, contacting the DNA strand are labeled in blue, contacting both DNA and RNA is labeled in purple. Conserved hydrophobic-core residues are highlighted in gray. (B) Overall structure of RNase H binding with Se-DNA/RNA hybrid. Se atoms are represented by ruby spheres in major groove, and magnesium atoms are represented by orange spheres at the interface of Protein and nucleic acid. (C) Stereoview of RNase H-substrate complex. The molecular surface of RNase H is shown in gray. Substrates of DNA/RNA hybrids are shown as rods with different colors to represent different DNAs: green: 5′-ATGTCG-3′; yellow: 5′-A^Se^TGTCG-3′, blue: 5′-ATG^Se^TCG-3′; purple: 5′-A^Se^TG^Se^TCG-3′. (D) Superimposed structures of T:rA and ^Se^T:rA base pairs at the active site. (E) Plot of chi (χ) torsion angles of the 2nd and 4th residues (T or ^Se^T) in the DNA strands (underlined residues). (F) Local structure of the active site of RNase H. Ruby sphere: Se; orange spheres: magnesium; red spheres: water. (G) Interactions between RNase H and Se-DNA/RNA hybrid. The contacting protein residues are shown.

We determined 4 different structures totally, which share the same RNase HC and RNA molecules. The DNA strand is modified accordingly (5’-ATGTCG-3′, 5′-A^Se^TGTCG-3′, 5′-ATG^Se^TCG-3′ and 5′-A^Se^TG^Se^TCG), to study the possible structural variation. The structures of RNase HC, with substrate of DNA/RNA or Se-DNA/RNA, are superimposable, with root-mean-square deviations (rmsd) of 0.122 Å (for native and double Se-duplex). The tertiary structure consists of a mixed β sheet with three antiparallel and two parallel strands, as well as three α helices (Figure 5B). In each asymmetric unit, one protein molecule binds one DNA/RNA hybrid, and binding pattern remains identical to the reported RNase H structures. As expected, the two selenium atoms on the 5-positions of thymidines of DNA strand are located in the major groove of the hybrid, which doesn’t directly contact the RNase H to interfere the substrate recognition. Moreover, since the major groove completely tolerates the Se-modifications, we don’t observe any structural perturbation in the Se-containing complexes. In the structures, the DNA strands are highly superimposable, and they follow the identical conformational features, including binding to DNA binding groove of RNase HC and contacting to several amino acid residues (Figure 5C). At the active site, the Se-T:A and T:A base pairs are highly superimposable (Figure 5D).

RNase H binds to DNA/RNA hybrid containing the mixed A and B sugar pucker conformations, while discriminating the dsRNA with 3’-endo conformation^66^. One important structural feature is, when binding to RNase H, the DNA strand adopts 2’-endo or 1’-exo sugar conformations. In our structures, we delightedly find, even after Selenium-modification, the thymidine residues still follow the 2’-endo sugar pucker. Compared to the native thymidine residues, selenium modifications result in mild local structural variation. For example, the T and 5-MeSe-T nucleotides present different chi (χ) torsion angles of the glycosidic bonds. The χ angles of Se-residues are generally ∼8oC less than Thymidines (Figure 5E). This is probably caused by the steric effect of Se and hydrophobicity of methyl group. However, the double helical geometry of the hybrid fully tolerates the derivatization, and the RNase H remains the identical conformational features to recognize the DNA strand, even in presence of selenium modifications. At the active site, we observed the two-metal-ion catalytic mechanism, in the same manner as in native RNase H/nucleic acid complex^65-67^. The four significant residues, D71, E109, N132 and D192, are observed to be directly involved in the substrate binding, by coordinating the two magnesium ions (Figure 5F). In the crystal structure, the catalytic carboxylate of D132 is mutated to N to inactivate the RNase H activity^66^, somehow, we still observe two Mg^2+^ ions and their coordinating waters to be positioned for the pseudo-catalysis. The 5-MeSe-T nucleobase on the complementary strand base pairs to the Adenosine at the active site, and the MeSe-modification doesn’t directly contact the amino acid residues of RNase HC. The interactions between the Se-DNA/RNA hybrid and RNase HC are shown in Figure 5G.

The minor groove of DNA is commonly characterized by its shallowness, rendering it a readily accessible target for a number of nucleic acid binding proteins^68^. In addition, certain chemical modifications to the DNA backbone, such as phosphorthioates, can intensify the protein binding, potentially leading to augmented cellular uptake as a drug^69^. However, these modifications on 2’-position of minor groove and phosphorthioates can easily alter the B-form geometry of antisense DNA, or loose the hybridization affinity to the target mRNA^70^. Our structures prove that, the selenium modification is situated in the deep major groove of DNA, preserving the sugar pucker conformation of DNA while potentially reducing protein binding and unintended toxicity. Therefore, the 5-MeSe-T modified DNA exhibits notable binding affinity to the complementary strands, while retaining RNase H activity, thus holding promise for gene silencing.

### Use Se-ASO to silence EGFP expression in cells

The molecular experimental results have revealed the stability in solution and compatibility to RNase H of Se-modified antisense oligonucleotides (Se-ASO). We then decided to investigate the gene silencing effect of Se-ASO in HeLa cells to suppress the EGFP expression.

EGFP was inserted into the plasmid in pcDNA 3.1+ vector, and the 18 nucleotides antisense is designed to target to the position 1496. Compared to the wild-type ASO, the Se-ASO has three thymidine residues replaced by 5-MeSe-T (Figure 6A). Both EGFP plasmid and ASO agent were delivered into HeLa cells using lipofectamine 2000 and incubated for 24h. The cells were then washed and visualized under Axiovert 200 fluorescence microscope excited by blue light. Without ASO co-transfected with EGFP plasmid, green fluorescence protein is actively expressed and strong fluorescent signal is observed (Figure 6B). When cells are transfected with ASO, in the groups of both native and Se-oligonucleotides, we observe suppressed EGFP signal. The result indicates the effective gene silencing of our synthetic DNA molecules. We also performed the EGFP expression assay by Western Blot. As shown in Figure 6C, both native and selenium modified ASOs result in significant decrease of EGFP expression, compared to the vehicle group and the group treated by the scrambled ASO. All the results strongly support the conclusion that Se-ASO has the potential as effective DNA therapy. In the cellular experiment, we haven’t observed more potent gene silencing effect of Se-ASO, compared to native antisense DNA. The reason could be the limited number of Se-modifications introduced to ASO, so the influence on stability is not significant enough. Moving forward, our aim is to utilize our Se-ASO as antisense agents to target other essential genes for interfering cell growth and proliferation, such as genes encoding bcl-2 family. Extending the treatment duration will be considered, to assess how the enhanced stability of Se-ASO can affect the gene silencing. Further optimization and more systematic cellular experiments will be needed in the future.

**Fig. 6.**
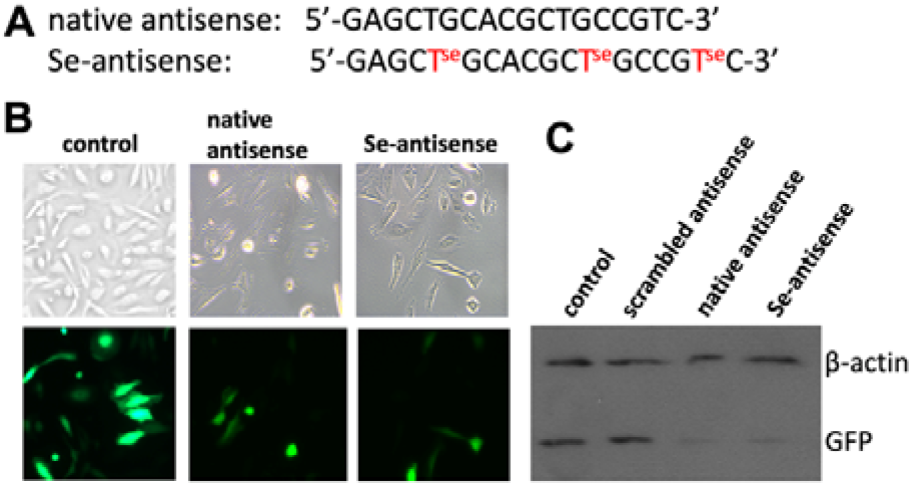
Gene silencing effect of Se-ASO. (A) Sequences of native ASO and Se-ASO to target EGFP expression. (B) Representative fluorescent microscopic images to show the reduction of EGFP expression by vehicle, native ASO and Se-ASO. (C) Western blot analysis of suppressing GFP protein expression with different antisense molecules.

## CONCLUSION

After the waning interest in antisense technology for a long period, there has been regained attention in this field because of the major improvement of chemical modifications that can provide high binding affinity, enhanced stability and low toxicity. The derivatization of DNA with chalcogen elements, especially selenium, offers an innovative possibility as antisense therapeutic application. The 5-methylselenyl-modified thymidine nucleoside and its oligonucleotides have been successfully synthesized quantitatively. Our thermostability assay and X-ray structures demonstrate the acceptable stability of Se-DNAs, likely due to the structural tolerance of DNA duplex and the enhanced stacking interaction. Moreover, we have observed the exceptional enzymatic stability of Se-DNAs, when treated with nucleases and serum. A reasonable hypothesis is that the selenium atom at 5-position has steric hindrance to destruct the molecular contacting between nuclease and major groove of DNA, leading to diminished binding and activity. Interestingly, even though the bulky selenium atom weakens nuclease catalysis, it causes no observable impact on RNase H activity when binding to complementary RNA as substrate. Our crystal structures of RNase H/nucleic acid complexes reveal the enzymatic binding and catalytic mechanism, which is completely compatible to Se-modification in the major groove because RNase H binds to the minor groove of DNA/RNA hybrids. The cellular experiment demonstrates the potential of Se-ASO to silence gene expression. As a novel antisense candidate, Se-ASO presents promise with its stability and activity to move forward to future in vitro and in vivo assessment. In the future, the investigation on the pharmacokinetics and subcellular distribution of Se-ASO is critical, since it is known that changing the chemistry of antisense oligonucleotides causes the change of organ distribution of drugs, independent of DNA sequences^71^. We hope Se-ASO could provide innovative strategy in this filed to modify antisense therapies, eventually making it a potential therapeutic candidate in clinic.

## MATERIAL AND METHODS

### Synthesis of 5-chalcogen-thymidine phosphoramidite (7)

Anhydrous and air-sensitive solvents and reagents were used and stored in between uses in a Vacuum Atmospheres Company (VAC) M040-2 glove box that was pressurized with nitrogen boil-off gas from a liquid nitrogen tank or in a VAC CS-40 glove box freezer at -20 °C. All starting materials for anhydrous reactions were dried prior to use on a vacuum line (1-4 ×10^−4^ torr). Reactions were monitored with glass-backed TLC plates pre-coated with silica gel 60 F_254_ (EMD Chemicals). Flash column chromatography was carried out using Fluka silica gel (60 Å pore, 230-400 mesh) that was packed in glass columns and pressurized with nitrogen. NMR Spectra were recorded on a Varian Unity +300 or Brucker Avance 400 spectrometer. Chemical shifts for ^1^H NMR were referenced relative to tetramethylsilane (0.00 ppm), CDCl_3_ (7.24 ppm) or DMSO (2.50 ppm). Chemical shifts for ^13^C NMR were referenced relative to CDCl_3_ (77.23 ppm) or DMSO (39.50 ppm). ^13^C NMR signals were assigned using ^13^C-APT technique. High resolution (HR) MS were either obtained with electrospray ionization (ESI) on a Q-TOFTM Waters Micromass at Georgia State University. The methods and characterization data for synthesis of 5-MeSe-T phosphoramidite and DNAs are listed here. The method to synthesize and analyze other modified DNAs is the same as 5-MeSe-T DNA. The other data for synthesis of 5-MeO-T, 5-MeS-T and 5-PhSe-T phosphoramidites and DNAs is provided in supporting information.

### Synthesis of modified DNAs

All the chalcogen-modified oligonucleotides were synthesized chemically on a 1.0 μmol scale using an ABI392 DNA/RNA Synthesizer. The concentration of the modified nucleoside phosphoramidites was identical to that of the conventional phosphoramidites (0.1 M in acetonitrile). Coupling was carried out using a 5-(benzylmercapto)-^1^H-tetrazole solution in acetonitrile. The coupling time was 25s. Syntheses were performed on control pore glass (CPG-500) immobilized with the appropriate nucleoside through a succinate linker (Glen Research). All the oligonucleotides were prepared with DMTr-on. After synthesis, the DNA oligonucleotides were cleaved from the solid support and fully deprotected by aqueous ammonia (conc.) treatment for 14 h at 55 °C. The 5’-DMTr deprotection of DNA oligonucleotides was performed in a 3% trichloroacetic acid solution (from 10% w/w, 0.9 M in water) for 1.5 min, followed by neutralization to pH 7.0 with a freshly made aqueous solution of triethylamine (1.1M) and petroleum ether extraction to remove DMTr-OH.

### HPLC purification and MS & HPLC analyses

The DNA oligonucleotides were analyzed and purified by reverse-phase high performance liquid chromatography (RP-HPLC) both DMTr-on and DMTr-off. Purification was carried out using a 21.2 × 250 mm Zorbax, RX-C18 column at a flow rate of 6 ml/min. Buffer A consisted of 20 mM triethylammonium acetate (TEAAc, pH 7.4, RNase-free water), while buffer B contained 50% aqueous acetonitrile and 50 mM TEAAc, pH 7.1. Similarly, analysis was performed on a Zorbax SB-C18 column (4.6 × 250 mm) at a flow rate of 1.0 ml/min using the same buffer system. The DMTr-on oligonucleotides were eluted with up to 90% buffer B in 25 min in a linear gradient, while the DMTr-off oligonucleotides were eluted with up to 40% of buffer B in a linear gradient in the same period of time. The collected fractions were lyophilized; the purified compounds were redissolved in RNase-free water. The pH was adjusted to 7.0 after the final purification of the oligonucleotides without the DMTr group. The typical HPLC profiles and MS spectra are listed in supporting information.

### Thermodenaturization of the modified duplex DNAs

We measured the melting temperatures of the Se-DNA duplexes along with those of the native duplexes. Prior to acquisition of the melting curves, duplexes were annealed by heating to 80 °C for two minutes, followed by slowly cooling to 5 °C and keeping at the temperature for three hours. Denaturation curves were acquired at 260 nm and 1 cm path length at heating or cooling rates of 0.5 °C/min, using a UV-Vis spectrophotometer equipped with a six-sample thermo-stated cell block and a temperature controller. The experiments were performed using the samples (DNA duplexes, 1.0 μM) dissolved in the buffer of 50 mM NaCl, 10 mM NaH_2_PO_4_-Na_2_HPO_4_ (pH 6.5), 0.1 mM EDTA and 10 mM MgCl_2_. The original melting curves are listed in supporting information.

### Enzymatic stability assay of modified DNAs

Both native and modified oligonucleotides were 5’-^32^P labeled by [γ-^32^P]-ATP using T4 polynucleotide kinase (New England Biolabs) following manufactures recommendations. Then γ-^32^P ATP was removed by size-exclusion column. The DNAs were redissolved in RNase-free water to a concentration 50 μM. The single strand DNA oligonucleotides were heated to 80 °C for 2 minutes with the complementary sequences, followed by slow cooling to room temperature to form the duplex. The digestion solutions were prepared by treating the DNA oligonucleotides with endonuclease or exonuclease in NEBuffer. The endonuclease or exonuclease enzymes were added into the solution containing DNA duplexes (2.5 μL), reaction buffer (0.5 μL) and water (1.5 μL). For AseI, the enzyme final concentration was 2.2 u/μL, and reaction was incubated for 16 h at 37 °C. For SalI, the enzyme final concentration was 4.0 u/μL, and reaction was incubated for 16 h at 37 °C. For Exonuclease III, the enzyme final concentration was 3.0 u/μL, and reaction was incubated for 15 min at 37 °C. For phosphordiesterase I, the enzyme final concentration was 0.5 u/μL and reaction was incubated for 30 min at 37 °C. The reactions were quenched by the addition of the gel loading dye solution (5 μL, containing 100 mM EDTA, saturated urea, 0.05% bromophenol blue, 0.05% xylene cyanol) and the analysis was performed on 15% polyacrylamide gel electrophoresis (PAGE), which was supplemented by 7 M urea. The gels were run for 2 h at 500 V, dried and exposed.

### RNase H activity assay of modified DNAs

The native RNA oligo was labeled with γ-^32^P using same strategy as above. After free γ-^32^P ATP was removed, RNA strand was annealed with native and modified DNAs by heating and slow cooling. RNase H enzyme was added to solutions containing NEBuffer to initiate the reaction. Aliquots were taken at different time points (0, 0.5, 1, 2, 3, 5, 10, 15 min), and the reaction was quenched by adding quenching buffer. The reaction products were analyzed by electrophoresis, following the same protocol as above.

### RNase H expression and purification for crystallization

RNase H mutant (D132N) construct was a gift from Dr. Wei Yang lab at NIH. Protein expression was carried out in BL21 (DE3) pLysS E. coli cells at 37°C in LB medium. Transformation was accomplished by heat shock method. A single colony was picked up to LB-ampicillin broth (20 mL). The culture was shaken (220 rpm) overnight at 37 °C. Two liters LB-ampicillin-chloramphenicol broth was prepared and the overnight culture was added to inoculate it. When the OD600 of inoculated broth reached at 0.7 OD, protein expression was then induced by adding IPTG (final concentration 1 mM) and the culture was shaken (220 rpm) overnight at 20 °C. Cells were harvested by centrifugation and then lysed by sonication.

RNase H enzyme was purified on a Ni column and eluted in 20 mM Tris-HCl (pH 7.5), 300 mM NaCl, 5% glycerol, 1.4 mM β-mercaptoethanol, and 300 mM imidazole. The buffer was exchanged to 20 mM Tris-HCl, 100 mM NaCl, 5% glycerol, 1.4 mM β-mercaptoethanol, and 0.5 mM EDTA for thrombin digestion. The proteins were further purified on a Phenyl Superose column with a 2 to 0.3 M gradient of ammonium sulfate, concentrated to 8–10 mg/ml, and stored in 20 mM HEPES (pH 7.0), 75 mM NaCl, 5% glycerol, 0.5mM EDTA, 2 mM DTT at 4°C or −20°C.

### Crystallization of DNA/RNA/RNase H complex

Prior to co-crystallization with RNase H, the purified Se-DNA or native DNA was annealed with the complementary RNA at 1:1 molar ratio, by first heating the mixture to 90 °C for 1 min, and then allowing it to cool slowly down to 25°C. The resulting Se-DNA/RNA duplex was mixed with the protein (final concentration: 6 mg/mL) at 1:1 molar ratio in the presence of 2 mM MgCl_2_. Co-crystallization of Se-DNA/RNA hybrid with RNase H was achieved by screening with the QIAGEN Classics Suite Kit (www.qiagen.com) by sitting-drop vapor diffusion method at 25°C. The optimal crystallization conditions are shown in SI.

### Data collection and structure determination

Crystal diffraction data of the Se-DNA/RNA/RNase H complex were collected at beamline 5.0.1 or 8.2.2 in the Advanced Light Source (ALS). 25% glycerol was used as cryoprotectant while X-ray data were collected under the liquid nitrogen stream at 100 K. The data were processed and scaled using HKL2000^72^, and the structures were solved by molecular replacement using 2G8U as search model. The resulted model was refined using Refmac5 within CCP4i^73^. The modified DNA was modeled into the structure using Coot^74^. The data collection and refinement statistics are shown in SI.

### Gene silencing by ASO in cells

HeLa cells were cultured in DMEM supplemented with 10% fetal bovine serum, 100 unit/mL penicillin and 100 μg/mL streptomycin, and grown at 37 °C with 5% CO_2_. For gene knockdown assay, cells were plated onto 6 well plates at a density of 1 × 105 cells/well in 2 mL of media. Before the gene knockdown assay, EGFP plasmid and plasmid/ASO transfection were carried out with Lipofectamine2000 (Thermo Fisher #11668030) following the manufacturer protocol. In the groups of antisense treatment, 5 nmole of different oligonucleotides were used per well. Fluorescent cell images were captured using a Zeiss Axiovert 200 inverted fluorescence microscope, with excitation wavelength of 457 - 487 nm, emission wavelength of 502 - 538 nm.

For Western blot analysis, cells were collected after transfection, and whole cell lysates were prepared by lysing the cells in RIPA buffer. Lysates were subjected to 12% polyacrylamide gel electrophoresis, and the proteins were transferred to nitrocellulose membrane for 1h. Membranes were blocked in 3% nonfat milk in TBS supplemented with 0.1% Triton X-100 and 0.02% sodium azide. EGFP was detected with a rabbit anti-GFP polyclonal antibody (1:3000), using chemiluminescent detection of ChemiDoc XRS (Bio-Rad). β-Actin was used as the loading control.

## SUPPORTING INFORMATION

The supporting information is available free of charge via the Internet at http://pubs.acs.org.

▪ NMR and MS spectra of nucleosides (Figure S1-S26)
▪ HPLC and MALDI-TOF MS spectra of oligonucleotides (Table S1 and Figure S27-S51)
▪ UV-melting temperature experiment (Figure S52-S54)
▪ Nuclease stability experiment (Figure S55-S81)
▪ MS spectrum of Se-antisense to suppress EGFP gene expression in HeLa cells (Figure S82)
▪ X-ray Crystallography statistics (Table S2 and S3)

## AUTHOR INFORMATION

### Author Contributions

‡These authors contributed equally. Z. Fang and W. Zhang wrote the manuscript with input from the other authors. All authors have given approval to the final version of the manuscript.

### Notes

Atomic coordinates and structure factors for the reported crystal structures have been deposited with the Protein Data bank under accession number 5usa, 5use, 5usg, 5wjr, 3ijk, 3hg8, 3ltu.

## ACKNOWLEDGMENT

We thank Drs. S. Wang and D. Walthall for their help on MS analysis. This research used resources of the Advanced Light Source (ALS), which is a DOE Office of Science User Facility under contract no. DE-AC02-05CH11231. We thank Drs. K. Royal and A. Rozales for their kindly help on X-ray diffraction data collection at the Advanced Light Source SIBYLS beamlines 501, 821 and 822. Plasmids expressing RNase H proteins were kindly given as gifts by Dr W. Yang at NIH. We thank all members from labs of Drs. W. Zhang and Z. Huang for the helpful discussion and insightful commentary of the manuscript.

