## Supporting Information for "Syntheses of Pyrimidine-Modified Seleno-DNAs as Stable Antisense Molecules"

### Supporting Information
for

### Syntheses of Pyrimidine-Modified Seleno-DNAs as Stable Antisense Molecules

Ziyuan Fang[1#], Yuliya Dantsu[2#], Cen Chen[3], Wen Zhang[2,4,*] and Zhen Huang[5,6,*]

[1] Department of Chemistry, University of Chicago, Chicago, IL 60637, USA
[2] Department of Biochemistry and Molecular Biology, Indiana University School of Medicine, Indianapolis, IN 46202, USA
[3] Firebird Biomolecular Sciences LLC, Alachua, FL 32615, USA
[4] Melvin and Bren Simon Cancer Center, 535 Barnhill Drive, Indianapolis, IN 46202, USA
[5] Key Laboratory of Bio-Resource and Eco-environment of Ministry of Education, College of Life Sciences, Sichuan University, Chengdu, Sichuan, 610064, P. R. China
[6] SeNA Research Institute and Szostak-CDHT Large Nucleic Acids Institute, Chengdu, Sichuan, 618000, P. R. China

[#] Contribute equally

#### Table of contents

Materials and general synthetic methods S2

$^{1}$H NMR, $^{13}$C NMR (APT), $^{31}$P NMR and MS spectra of nucleosides S2

HPLC and MALDI-TOF MS spectra of oligonucleotides S21

UV-melting temperature experiment S36

Nuclease stability experiment S38

MS spectrum of Se-antisense to knockdown EGFP expression in cells S50

X-ray crystallography statistics S51

**1. Materials and general procedures.** Anhydrous and air-sensitive solvents and reagents were used and stored in between uses in a Vacuum Atmospheres Company (VAC) M040-2 glove box that was pressurized with nitrogen boil-off gas from a liquid nitrogen tank or in a VAC CS-40 glove box freezer at -20 °C. Solvents were dried and redistilled using standard methods. Distilled solvents and reagents were transferred under nitrogen gas to the glove box immediately after distillation using an evacuated Schlenk tube or flask containing activated molecular sieves. All starting materials for anhydrous reactions were dried prior to use on a vacuum line (1 - 4 × $10^{-4}$ torr). Reactions were monitored with glass-backed TLC plates pre-coated with silica gel 60 $F_{254}$ (EMD Chemicals). Flash column chromatography was carried out using Fluka silica gel (60 Å pore, 230-400 mesh) that was packed in glass columns and pressurized with nitrogen. NMR Spectra were recorded on a Varian Unity 300 or Brucker Avance 400 spectrometer. Chemical shifts for $^1$H NMR were referenced relative to tetramethylsilane (0.00 ppm), $CDCl_3$ (7.24 ppm) or DMSO (2.50 ppm). Chemical shifts for $^{13}$C NMR were referenced relative to $CDCl_3$ (77.23 ppm) or DMSO (39.50 ppm). $^{13}$C NMR signals were assigned using $^{13}$C-APT technique. High resolution (HR) MS were either obtained with electrospray ionization (ESI) on a Q-TOFTM Waters Micromass at Georgia State University or Indiana University.

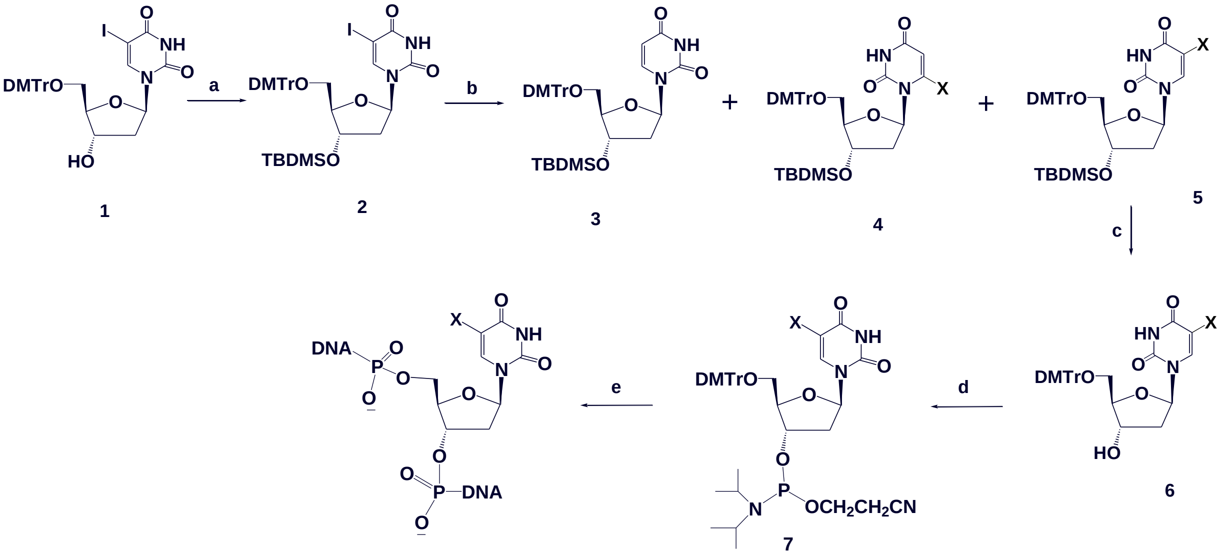


Scheme S1: Synthesis of 5-chalcogen-T DNAs. Reagents and conditions: a) TBDMS-Cl, Im., DMF, 4 h, rt, 92 %; b) i) NaH, THF, 15 min, r.t., ii) lithiating agent, THF, 30 min., -78 °C, iii) $X_2$, 1 h, -78 °C; c) TBAF, THF, 4 h , rt, 92%; d) i-$Pr_2NP(Cl)CH_2CH_2CN$, DIEA, $CH_2Cl_2$, 1 h, rt, 82%; e) solid phase synthesis. X=$CH_3Se$, $CH_3S$ or PhSe.

**2. Spectra of synthetic nucleosides.**

**3'-O-*tert*-Butyldimethylsilyl-5'-O-(4,4-dimethoxytrityl-2'-deoxy-5-methylselenyl-thymidine (5, X=$CH_3$Se).** NaH 95% (35.5 mg, 1.4 mmol) was added portion wise to a solution of **2** (1.03 g, 1.33 mmol) in dry THF (7 mL) at room temperature in dry glove box. The mixture was stirred for 30 min until complete cease of hydrogen gas evolution, then cooled down to -78 °C and treated with 1.4 M solution of *n*-BuLi in hexanes (2.1 mL, 2.93 mmol) was added dropwise over 10 min. The mixture was stirred for 30 min then treated with $(CH3)_2Se_2$ (0.5 mL, 5.32 mmol) and the mixture was further stirred for 1 h at the same temperature. Saturated solution of $NH_4Cl$ (5 mL) was added and the mixture was warmed to room temperature.

Ethylacetate was added to the mixture and the whole was washed with $H_2O$, brine, dried over $MgSO_4$, and evaporated under reduce pressure. The residue was purified by flash silica gel chromatography (eluate: 20% EtOAc in hexanes) gave **5** (0.83 g, 85%) as a colorless foam. Elution with 25% EtOAc in hexanes gave **3** (45 mg, 5%) as a colorless foam. Spectral Data for 5: $^1$H-NMR ($CDCl_3$): 8.93 (1H, br s, NH, exchanged with $D_2O$), 7.99 (1H, s, H-6), 7.47-7.22 (9 H, m, Ar), 6.87-6.84 (4H, m, Ar), 6.31 (1H, dd, H-1', J = 5.9, J = 7.6 Hz), 4.46 (1H, m, H-3'), 4.02 (1H, m, H-4'), 3.81 (6H, s, $CH_3O$), 3.41 (1H, dd, H-5'a, J = 3.2, J = 10.7 Hz), 3.31 (1H, dd, H-5'b, J = 3.4, J = 10.7 Hz), 2.39 (1H, ddd, H-2'a, J = 2.6, J = 5.8, J = 13.2 Hz), 2.19 (1H, m, H-2'b), 2.05 (9 H, s, *tert*-Butyl), 0.05 (3H, s, CH3), 0.01 (3H, s, $CH_3$); $^{13}$C-NMR ($CDCl_3$): 161.62 (C4), 158.67 (Ar), 150.09 (C2), 144.42 (Ar), 141.57 (C-6), 135.60 (Ar), 135.54 (Ar), 130.11 (Ar), 128.12 (Ar), 127.95 (Ar), 127.00 (Ar), 113.28 (Ar), 103.56 (C-5), 87.13 (Ar), 86.86 (C4'), 85.55 (C-1'), 72.54 (C-3'), 63.13 (C-5'), 55.25 (OMe), 41.72 (C2'), 25.74 ($CMe_3$),17.95 ($CMe_3$), 7.30 ($SeCH_3$), -4.69 ($SiMe_2$), -4.86 ($SiMe_2$); HRMS (ESI-TOF): Molecular formula $C_{37}H_{45}N_2O_7SeSi$ $[M-H]^+$: 737.2147 (calc.737.2161).

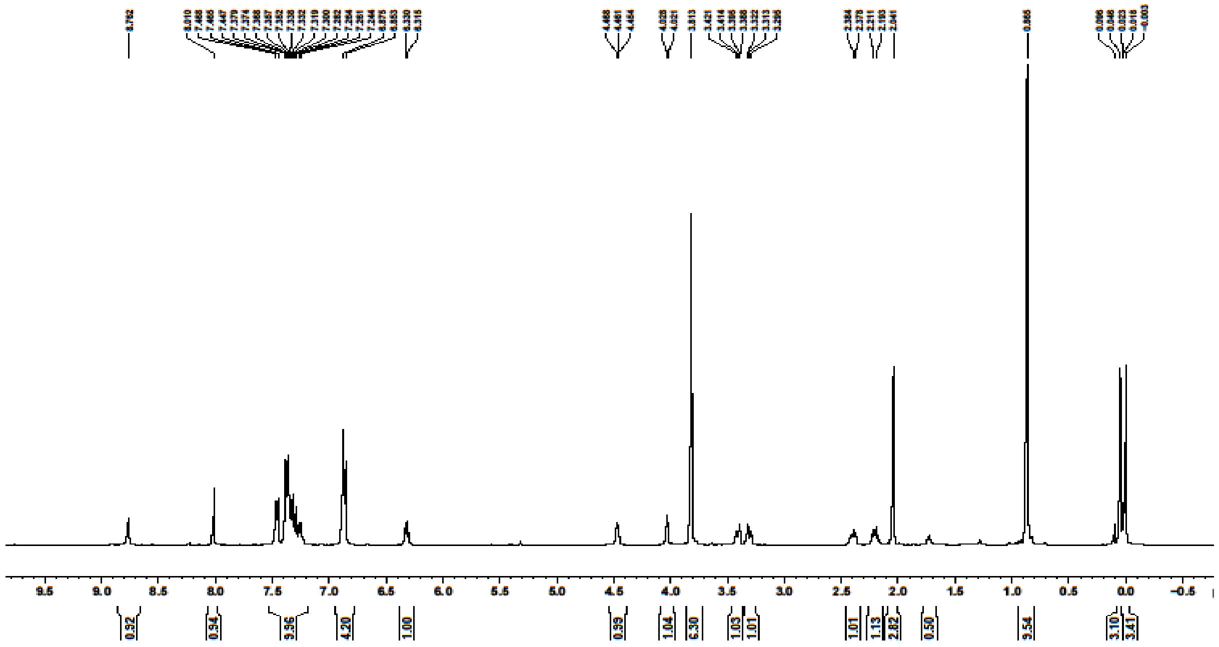

Figure S1. 1H NMR of compound **5** (X=$CH_3$Se).

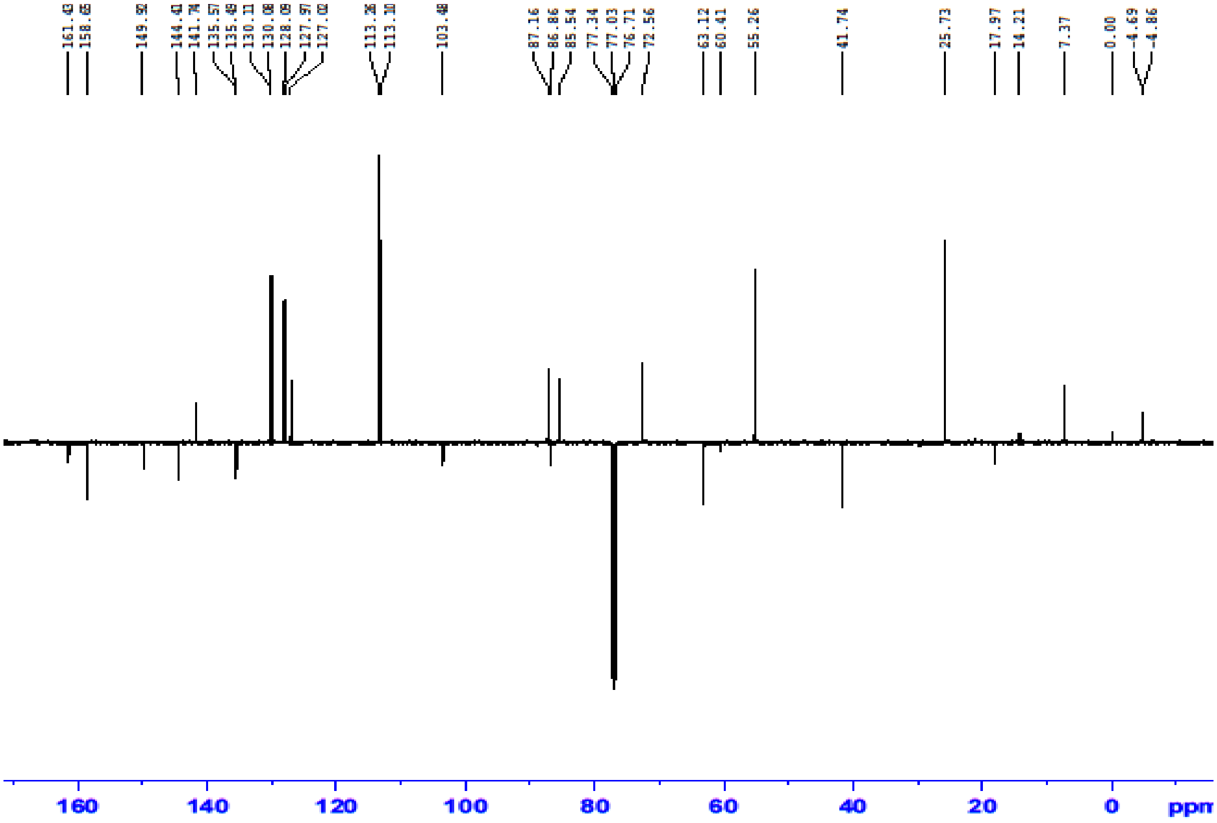


Figure S2. [13]C NMR of compound **5** (X=$CH_3Se$).

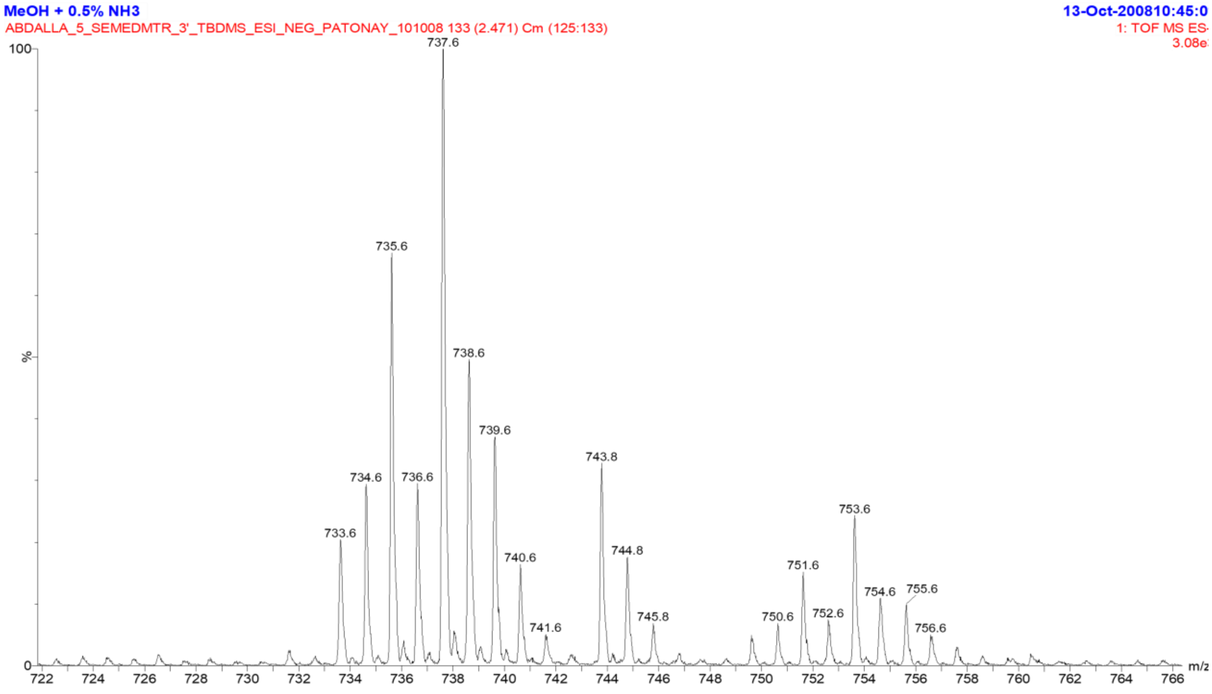


Figure S3. MS spectrum of compound **5** (X=$CH_3Se$).

**5'-O-(4,4-dimethoxytrityl-2'-deoxy-5-methylselenylthymidine (6, X=$CH_3$Se).** A 1 M solution of TBAF in THF (1.55 mL) was added to a solution of **5** (0.75 g, 1.02 mmol) in THF (15 mL) at 0 ºC. The mixture was stirred for 6 h at room temperature. The solvent was evaporated and the residue partitioned between EtOAc and $H_2O$. The organic phase was dried over $MgSO_4$ and evaporated the residue was purified by silica gel column chromatography (the silica gel was pre-equalized with 1% $Et_3N$ in $CH_2Cl_2$, eluate 4% MeOH in $CH_2Cl_2$) to give (0.58 mg, 92%) of **6** as pale yellow foam: $^1$H-NMR ($CD_2Cl_2$) 9.39 (1H, s, NH, exchanged with $D_2O$), 7.90 (1H, s, H-6), 7.46-7.21 (9H, m, Ar), 6.87-6.84(4H, m, Ar), 6.29 (1H, dd, H-1’, J = 6.4, J = 7.6 Hz), 4.43 (1H, m, H-3'), 4.05 (1H, m, H-4'), 3.68 (6H, 2 s, OMe), 3.30 (1H, dd, H5'a, J = 3.8, J = 10.5 Hz), 3.24 (1H, dd, H5'b, J = 3.6, J = 10.5 Hz), 2.58 (1H, d, 3'-OH), 2.42 (1H, ddd, H-2'a, J = 3.8, J = 7.7, J = 10.8 Hz), 2.36 (1H, m, H-2'b), 1.90 (3H, s, $SeCH_3$); $^{13}$C-NMR ($CD_2Cl_2$) 162.48 (C4), 159.29 (Ar), 150.94 (C2), 145.27 (Ar), 141.81(C-6), 136.21 (Ar), 136.06 (Ar), 130.64 (Ar), 130.62 (Ar), 128.57 (Ar), 128.50 (Ar), 127.47 (Ar), 113.76 (Ar), 104.21 (C-5), 87.33 (Ar), 86.82 (C4'), 85.89 (C-1'), 72.82 (C-3'), 64.24 (C-5'), 41.55 (C2'), 7.44 ($SeCH_3$); HRMS (ESI-TOF): Molecular formula $C_{31}H_{32}N_2O_7Se$ $[M-H]^-$: 623.1735 (calc. 623.1260).

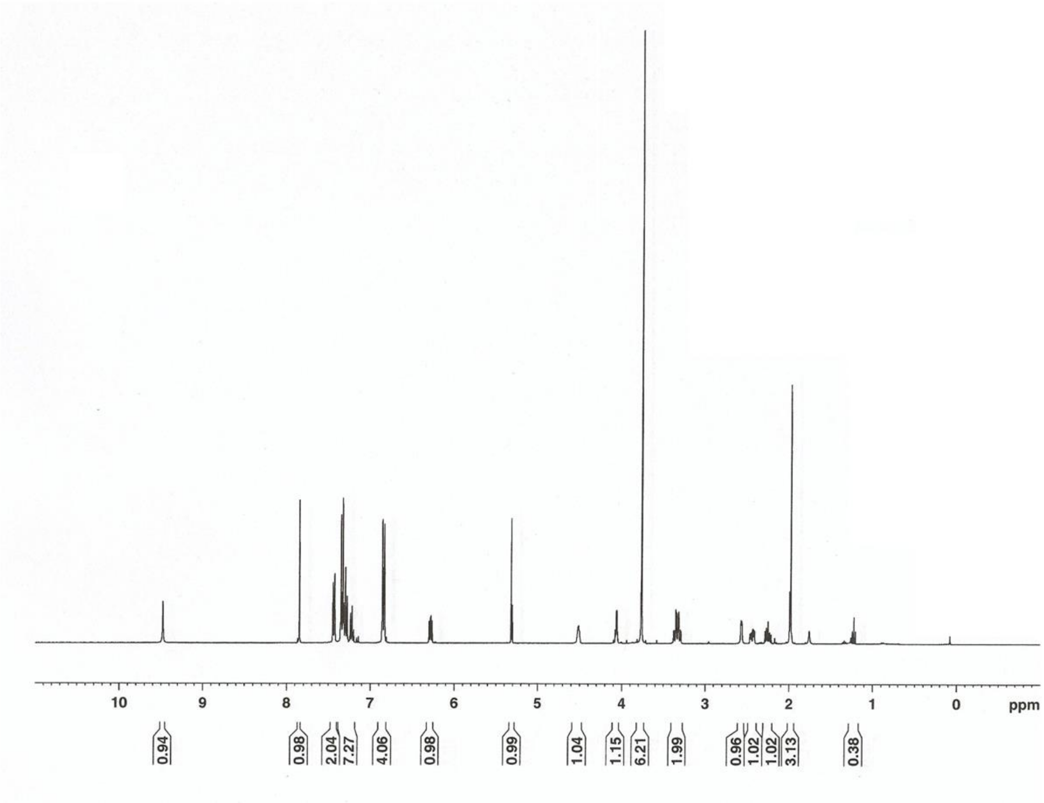


Figure S4. 1H NMR of compound **6** (X=$CH_3$Se).

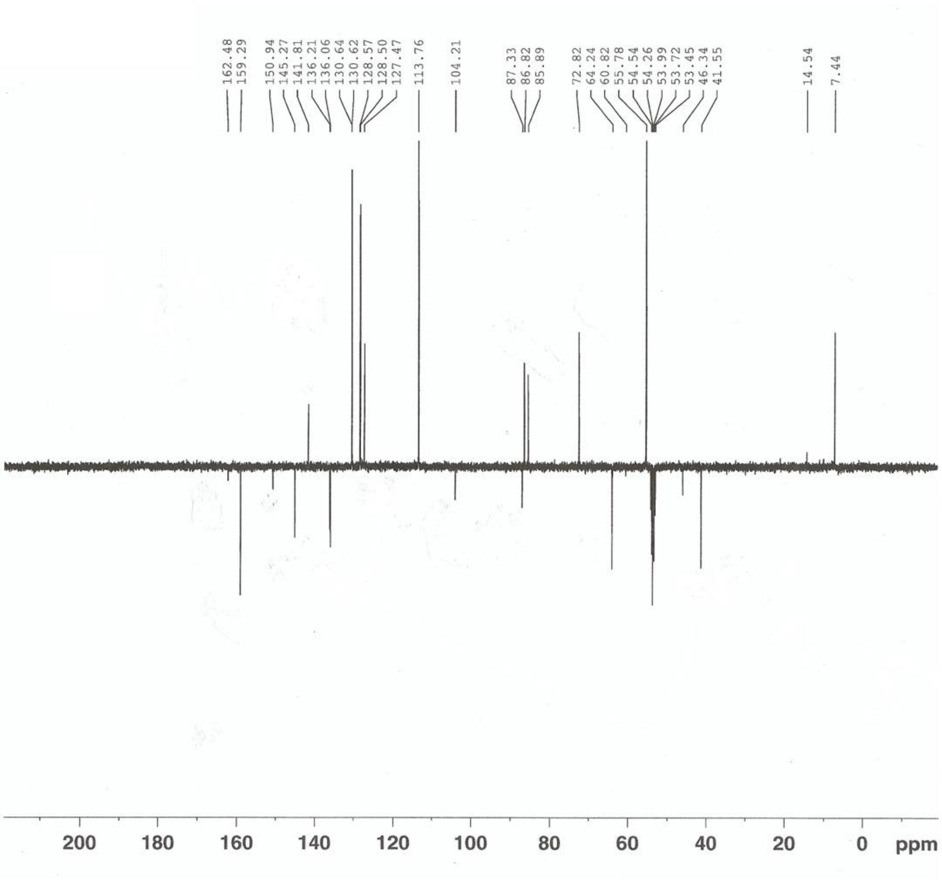


Figure S5. $^{13}C$ NMR of compound **6** ($X=CH_3Se$).

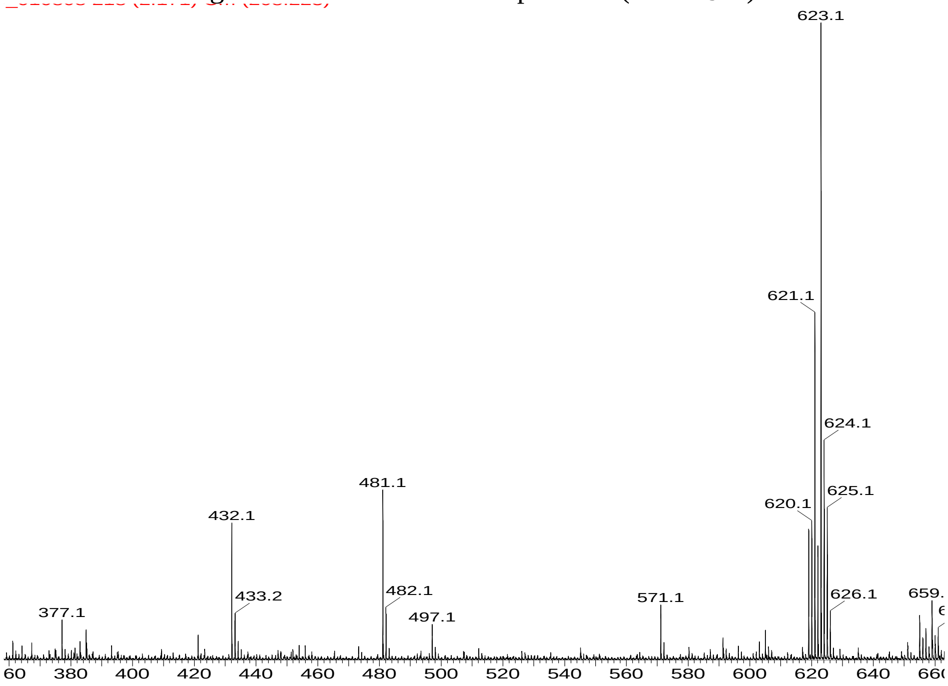


Figure S6. MS spectrum of compound **6** ($X=CH_3Se$).

**1-[2'-deoxy-3'-O-(2-cyanoethyl-*N,N*-diisopropylamino)-phosphoramidite-5'-O-(4,4-dimethoxytrityl-[D-erythro-ribofuranosyl]-5-methylselenylthymidine (7, X=$CH_3$Se).** Diisopropyl-ethylamine (20 mL, 0.12 mmol) was added to a solution of **6** (0.3 g, 0.48 mmol), 5-benzylthiotetrazole (45.4 mg, 0.24 mmol) and 2-cyanoethyl-*N,N,N,N*-tetraisopropyl phosphane (288 mg, 0.96 mmol) in dry $CH_2Cl_2$ (15 mL) at 0 $^oC$. The mixture was stirred for 2 h at room temperature then slowly poured into pentane (100 mL). The produced white precipitate was filtered off, dissolved in $CH_2Cl_2$ (1 mL), and precipitated in pentane. The collected fine powdered white solid was dissolved in $CH_2Cl_2$ and dried under reduced pressure to give 324 mg, 82% of **7** as a mixture of two diastereomers and was directly used for solid phase synthesis. An analytically pure sample was purified by a preparative TLC (eluate: 30% EtOAc in $CH_2Cl_2$) to give a mixture of two diastereomers: $^1$H-NMR ($CD_3CN$, two sets of signals for a mixture of two diastereomers): 9.16 (1H, s, NH, exchanged with $D_2O$), 7.76 and 7.17 (1H each, s, H-6), 7.52-7.22 (9H, m, DMTr), 6.95-6.82 (4H, m, DMTr), 6.19 (1H, dd, H-1'), 4.57 (1H, m, H-3'), 4.10 and 4.05 (1H, m, H-4'), 3.75 (6H, 2 s, OMe), 3.65 and 3.55 (m, CH-ipr), 3.30 (2H, m, H5'a and H5'b), 2.63-2.51 (2H, dd, $CH_2$), 2.47-2.31 (1H, H-2'a and H-2'b), 1.96 (3H, s, Se$CH_3$), 1.17-1.03 (2x 24H, m, $CH_3$-iPr); $^{13}$C-NMR ($CD_3CN$, two sets of signals for a mixture of two diastereomers) 162.63 (C4), 159.82 (Ar), 151.21 (C2), 146.01 (Ar), 141.65 and 141.57 (C-6), 136.84 (Ar), 136.79 (Ar), 136.73 (Ar), 132.30 (Ar), 131.20 (Ar), 131.17 (Ar), 131.15 (Ar), 129.76 (Ar), 129.12 (Ar), 127.98 (Ar), 114.20 (Ar), 104.44 and 104.32 (C-5), 118.80 and 118.38 (CN), 87.52 (Ar), 86.38 and 86.34 (C4'), 86.14 and 86.08 (C-1'), 74.54 and 74.37 (C-3'), 64.41 and 64.23 (C-5'), 59.64 and 59.45 (OMe), 44.15 and 44.03 (C2'), 40.47,40.43, 40.37, 40.32 (CH-iPr), 24.99, 24.93 and 24.86 ($CH_3$-iPr), 21.13, 21.06, 20.99 ($CH_2$), 7.09 and 7.05 (Se$CH_3$); $^{31}$P-NMR ($CD_3CN$) 148; HRMS (ESI-TOF): Molecular formula, $C_{40}H_{49}N_4O_8PSe$; $[M-H]^-$: 823.2374 (calc. 823.2453).

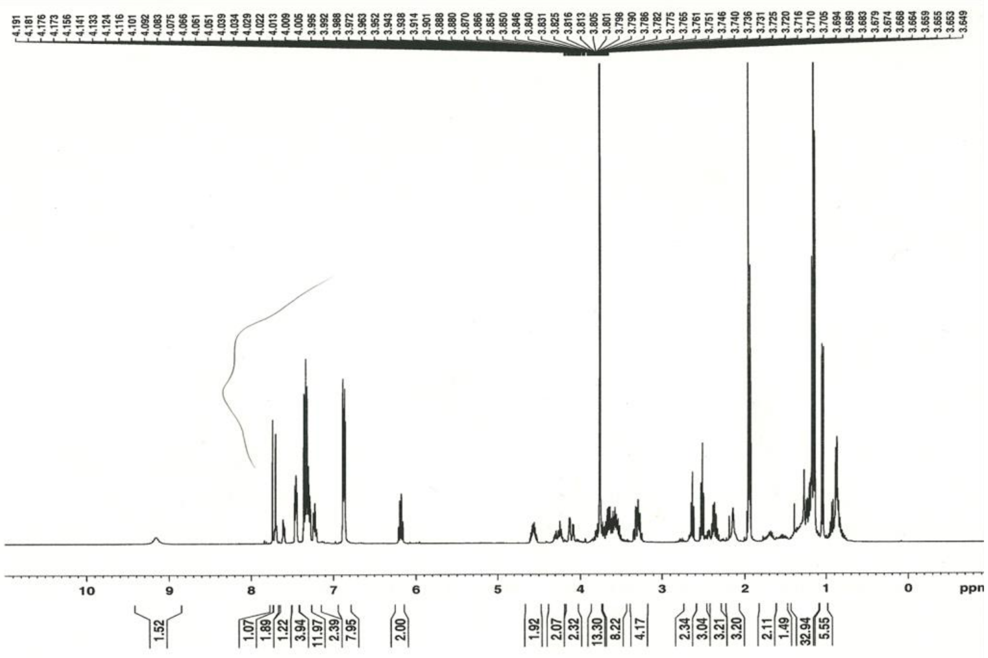

Figure S7. $^1$H NMR of compound **7** (X=$CH_3$Se).

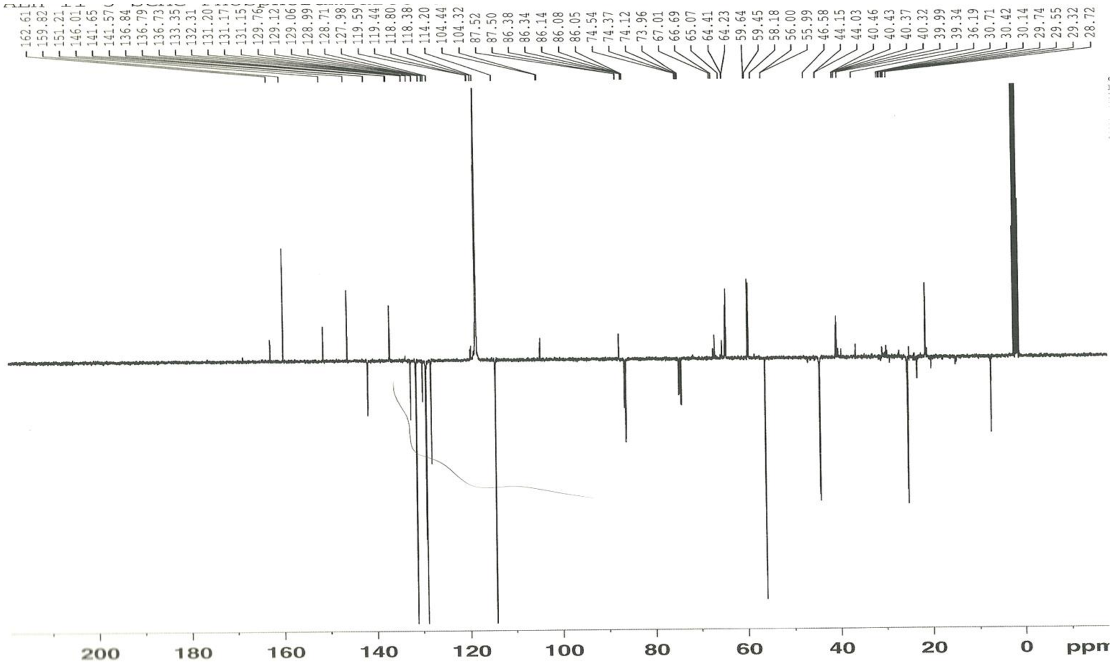


Figure S8. $^{13}C$ NMR of compound **7** (X=$CH_3$Se).

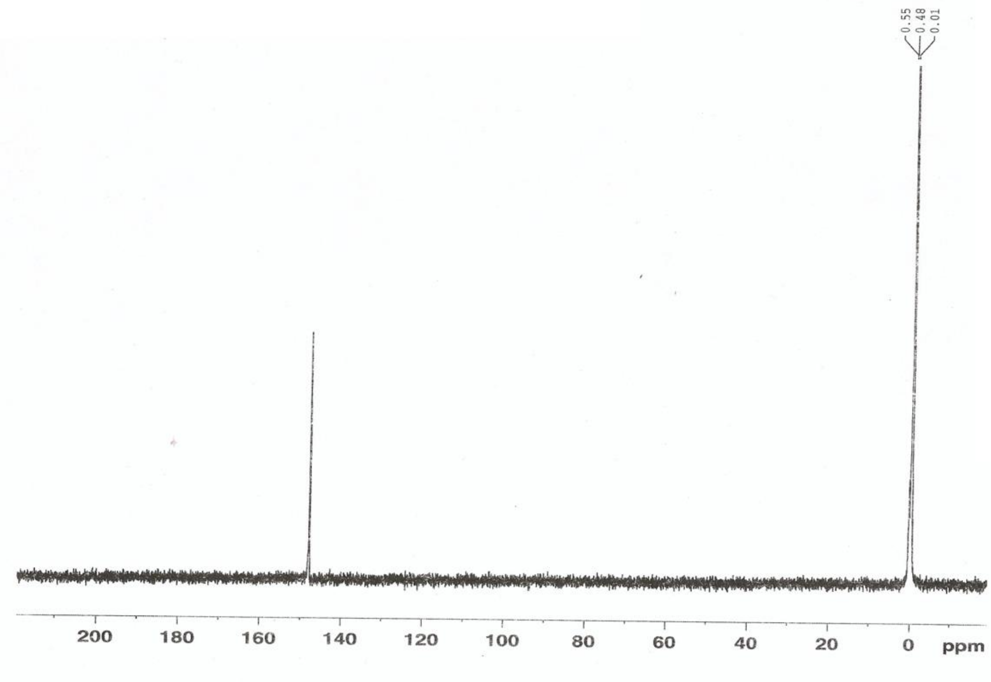


Figure S9. $^{31}P$ NMR of compound **7** (X=$CH_3$Se).

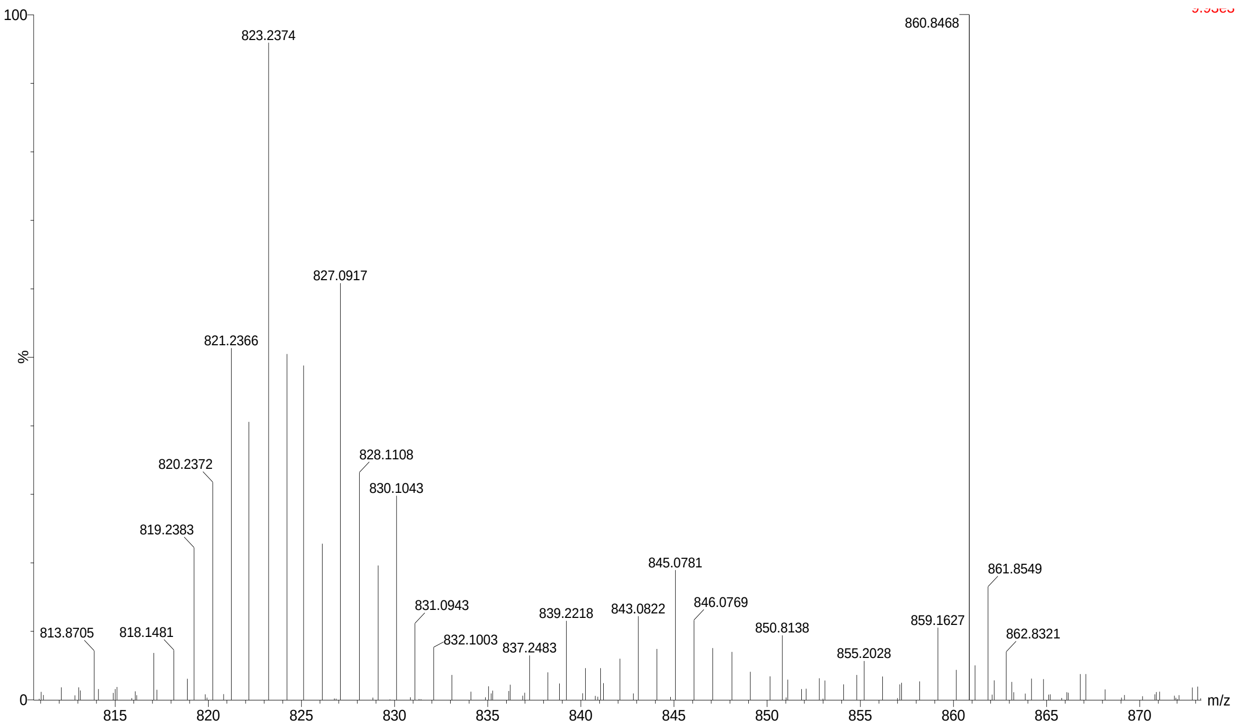


Figure S10. MS spectrum of compound **7** (X=$CH_3Se$).

**3'-O-*tert*-Butyldimethylsilyl-5'-O-(4,4-dimethoxytrityl-2'-deoxy-5-methylthio-thymidine (5, X=$CH_3S$).** Operation was the same as the synthesis of 5-$CH_3Se$-derivatized nucleoside, and yield was 88%. $^1$H-NMR ($CDCl_3$): 8.93 (1H, br s, NH, exchanged with D2O), 7.99 (1H, s, H-6), 7.47-7.22 (9 H, m, Ar), 6.87-6.84 (4H, m, Ar), 6.31 (1H, dd, H-1', J = 5.9, J = 7.6 Hz), 4.46 (1H, m, H-3'), 4.02 (1H, m, H-4'), 3.81 (6H, s, CH3O), 3.41 (1H, dd, H-5'a, J = 3.2, J = 10.7 Hz), 3.31 (1H, dd, H-5'b, J = 3.4, J = 10.7 Hz), 2.39 (1H, ddd, H-2'a, J = 2.6, J = 5.8, J = 13.2 Hz), 2.19 (1H, m, H-2'b), 2.05 (9 H, s, *tert*-Butyl), 0.05 (3H, s, $CH_3$), 0.01 (3H, s, $CH_3$); $^{13}$C-NMR ($CDCl_3$): 161.62 (C4), 158.67 (Ar), 150.09 (C2), 144.42 (Ar), 141.57 (C-6), 135.60 (Ar), 135.54 (Ar), 130.11 (Ar), 128.12 (Ar), 127.95 (Ar), 127.00 (Ar), 113.28 (Ar), 103.56 (C-5), 87.13 (Ar), 86.86 (C4'), 85.55 (C-1'), 72.54 (C-3'), 63.13 (C-5'), 55.25 (OMe), 41.72 (C2'), 25.74 ($CMe_3$),17.95 ($CMe_3$), 7.30 ($SeCH_3$), -4.69 ($SiMe_2$), -4.86 ($SiMe_2$).

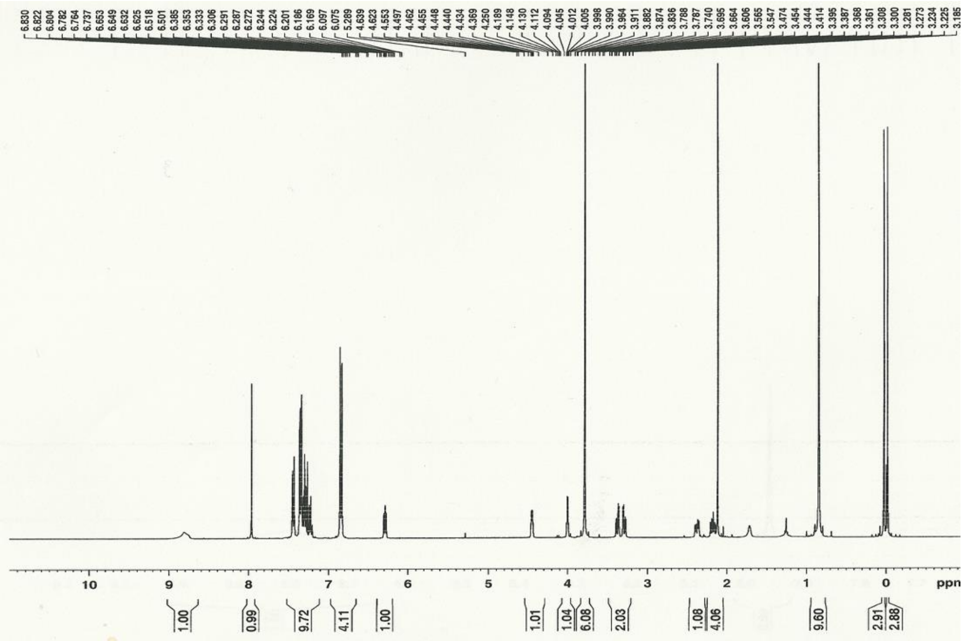

Figure S11. $^{1}$H NMR of compound **5** (X=$CH_3S$).

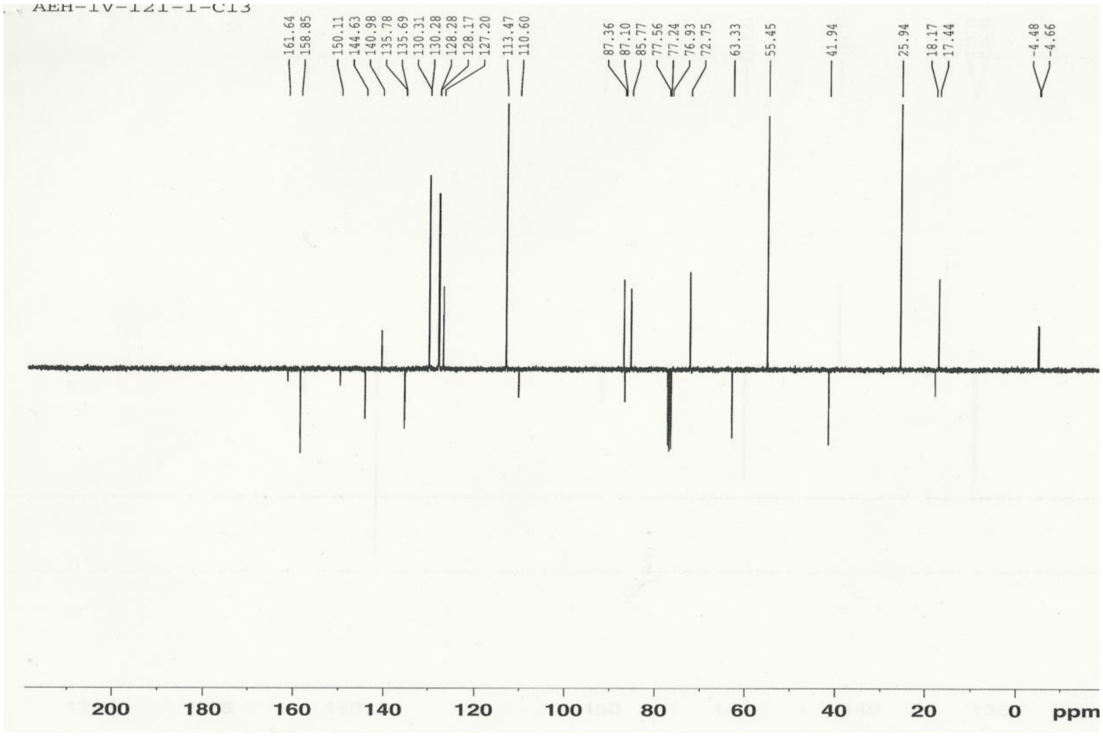

Figure S12. $^{13}$C NMR of compound **5** (X=$CH_3S$).

**5'-O-(4,4-dimethoxytrityl-2'-deoxy-5-methylthio-thymidine (6, X=$CH_3$S).** Operation was the same as the synthesis of 5-$CH_3$Se-derivatized nucleoside, and yield was 91%. $^1$H-NMR ($CD_2Cl_2$) 9.39 (1H, s, NH, exchanged with $D_2O$), 7.90 (1H, s, H-6), 7.46-7.21 (9H, m, Ar), 6.87-6.84(4H, m, Ar), 6.29 (1H, dd, H-1', J = 6.4, J = 7.6 Hz), 4.43 (1H, m, H-3'), 4.05 (1H, m, H-4'), 3.68 (6H, 2 s, OMe), 3.30 (1H, dd, H5'a, J = 3.8, J = 10.5 Hz), 3.24 (1H, dd, H5'b, J = 3.6, J = 10.5 Hz), 2.58 (1H, d, 3'-OH), 2.42 (1H, ddd, H-2'a, J = 3.8, J = 7.7, J = 10.8 Hz), 2.36 (1H, m, H-2'b), 1.90 (3H, s, SeCH$_3$); $^{13}$C-NMR ($CD_2Cl_2$) 162.48 (C4), 159.29 (Ar), 150.94 (C2), 145.27 (Ar), 141.81(C-6), 136.21 (Ar), 136.06 (Ar), 130.64 (Ar), 130.62 (Ar), 128.57 (Ar), 128.50 (Ar), 127.47 (Ar), 113.76 (Ar), 104.21 (C-5), 87.33 (Ar), 86.82 (C4'), 85.89 (C-1'), 72.82 (C-3'), 64.24 (C-5'), 41.55 (C2'), 7.44 (SeCH$_3$); HRMS (ESI-TOF): Molecular formula $C_{31}H_{32}N_2O_7S$ [M-H]$^+$: 575.2209 (calc. 575.1930).

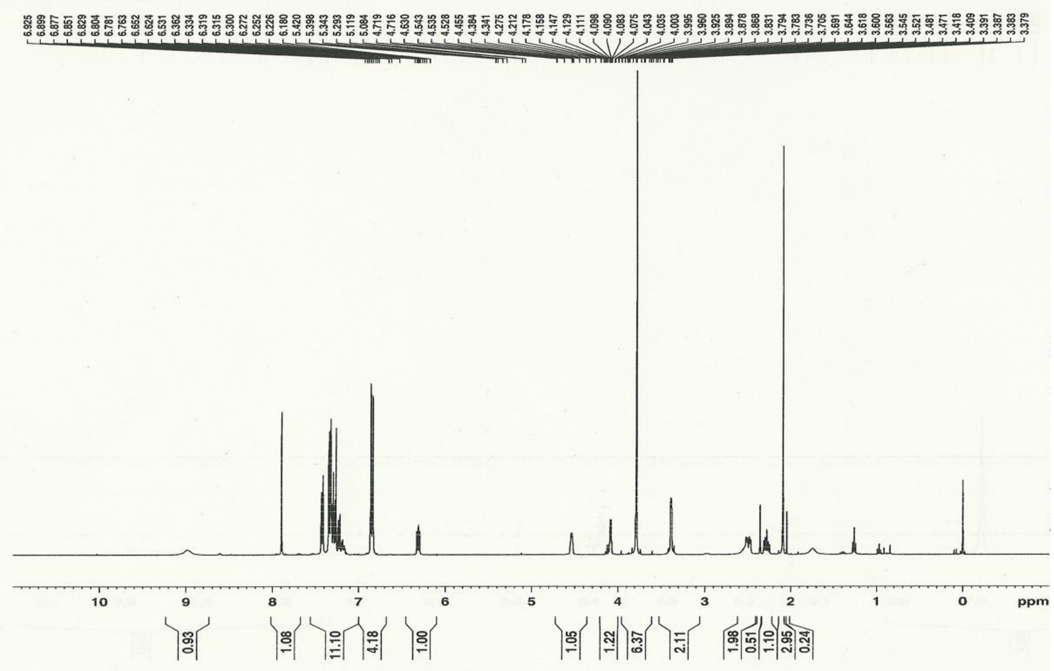


Figure S13. $^1$H NMR of compound **6** (X=$CH_3$S).

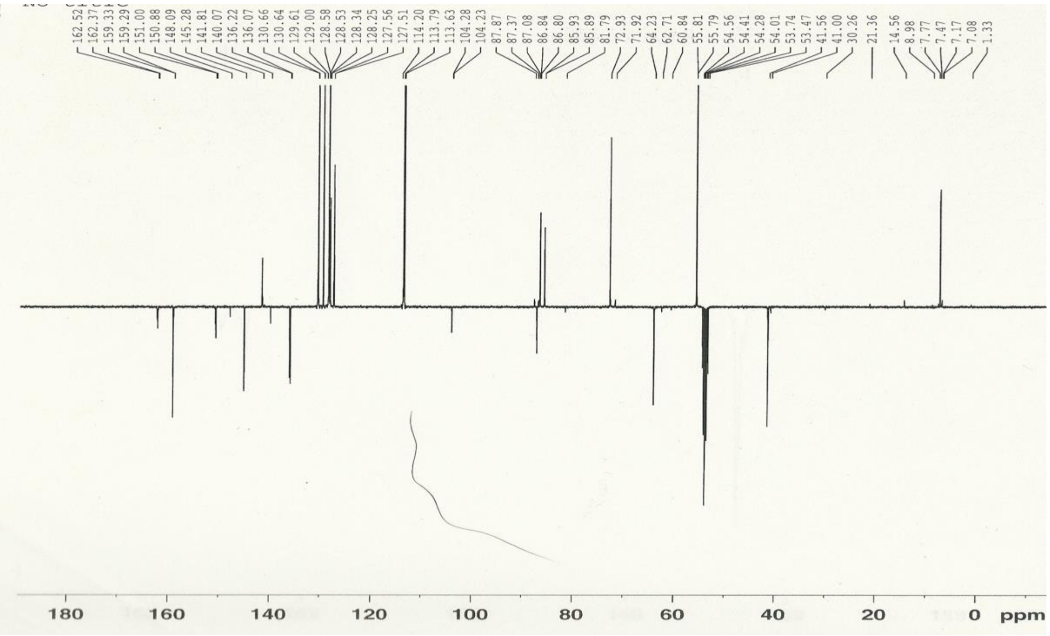

Figure S14. $^{13}C$ NMR of compound **6** ($X=CH_3S$).

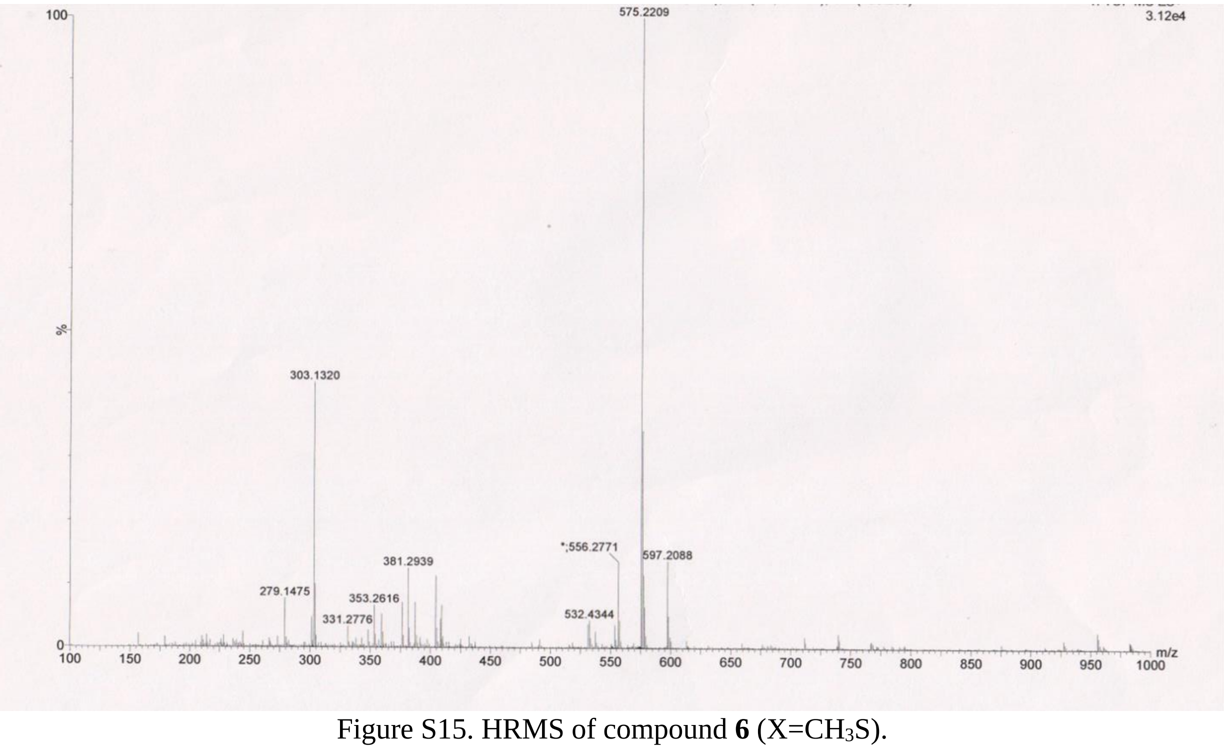

Figure S15. HRMS of compound **6** ($X=CH_3S$).

**1-[2'-deoxy-3'-O-(2-cyanoethyl-*N*,*N*-diisopropylamino)-phosphoramidite-5'-O-(4,4-dimethoxytrityl-[D-erythro-ribofuranosyl]-5-methylthio-thymidine (7, $X=CH_3S$).** Disopropyl-ethylamine (20 mL, 0.12 mmol) was added to a solution of 6 (0.3 g, 0.48 mmol), 5-benzylthiotetrazole (45.4 mg, 0.24 mmol) and 2-cyanoethyl-*N*,*N*,*N*,*N*-tetraisopropyl phosphane

(288 mg, 0.96 mmol) in dry $CH_2Cl_2$ (15 mL) at 0 °C. The mixture was stirred for 2 h at room temperature then slowly poured into pentane (100 mL). The produced white precipitate was filtered off, dried under high vacuum and directly applied in solid phase synthesis without further purification.

**3'-O-*tert*-Butyldimethylsilyl-5’-O-(4,4-dimethoxytrityl-2'-deoxy-5-phenylselenyl-thymidine (5, X=PhSe).** Operation was the same as the synthesis of 5-$CH_3$Se-derivatized nucleoside, and yield was 81%. $^1$H-NMR ($CDCl_3$) 8.93 (1H, br s, NH, exchanged with D2O), 7.99 (1H, s, H-6), 7.47-7.22 (9 H, m, Ar), 6.87-6.84 (4H, m, Ar), 6.31 (1H, dd, H-1', J = 5.9, J = 7.6 Hz), 4.46 (1H, m, H-3'), 4.02 (1H, m, H-4'), 3.81 (6H, s, $CH_3$O), 3.41 (1H, dd, H-5'a, J = 3.2, J = 10.7 Hz), 3.31 (1H, dd, H-5'b, J = 3.4, J = 10.7 Hz), 2.39 (1H, ddd, H-2'a, J = 2.6, J = 5.8, J = 13.2 Hz), 2.19 (1H, m, H-2'b), 2.05 (9 H, s, *tert*-Butyl), 0.05 (3H, s, $CH_3$), 0.01 (3H, s, $CH_3$); HRMS (ESI-TOF): Molecular formula $C_{42}H_{48}N_2O_7SeSi$ $[M+Na]^+$: 823.2403 (calc.823.2396).

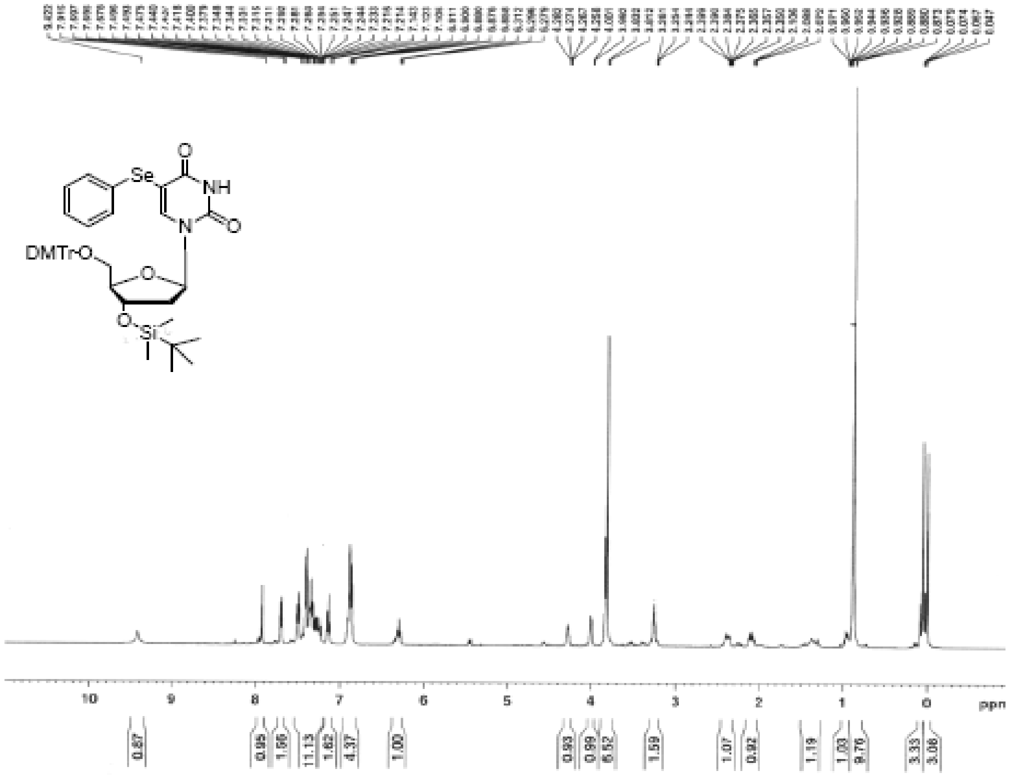


Figure S16. $^1$H NMR of compound **5** (X=PhSe).

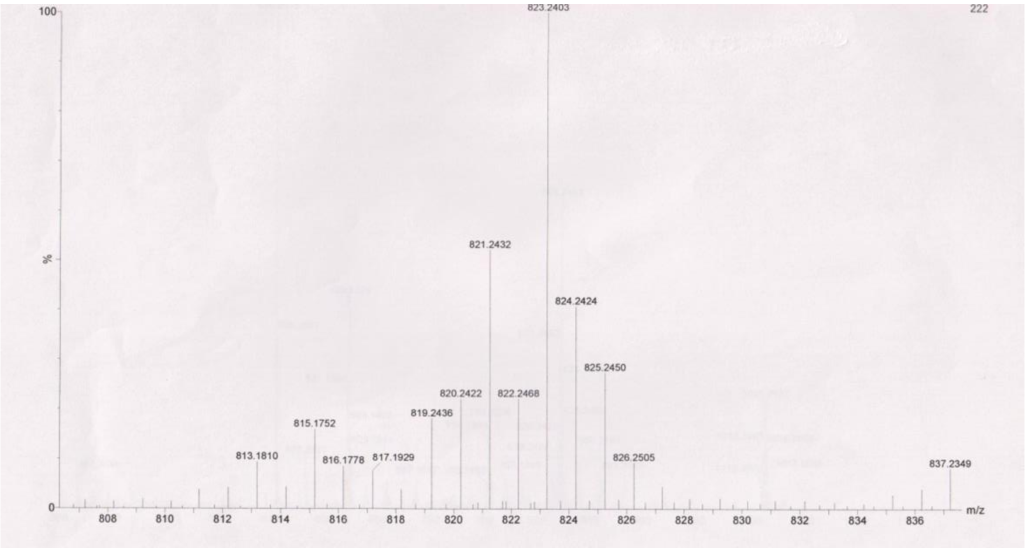

Figure S17. HRMS of compound **5** (X=PhSe).

**5'-O-(4,4-dimethoxytrityl-2'-deoxy-5-phenylselenylthymidine (6, X=PhSe).** Operation was the same as the synthesis of 5-$CH_3$Se-derivatized nucleoside, and yield was 90%. $^1$H-NMR ($CD_2Cl_2$) 9.39 (1H, s, NH, exchanged with $D_2O$), 7.90 (1H, s, H-6), 7.46-7.21 (9H, m, Ar), 6.87-6.84(4H, m, Ar), 6.29 (1H, dd, H-1', J = 6.4, J = 7.6 Hz), 4.43 (1H, m, H-3'), 4.05 (1H, m, H-4'), 3.68 (6H, 2 s, OMe), 3.30 (1H, dd, H5'a, J = 3.8, J = 10.5 Hz), 3.24 (1H, dd, H5'b, J = 3.6, J = 10.5 Hz), 2.58 (1H, d, 3'-OH), 2.42 (1H, ddd, H-2'a, J = 3.8, J = 7.7, J = 10.8 Hz), 2.36 (1H, m, H-2'b), 1.90 (3H, s, Se$CH_3$); $^{13}$C-NMR ($CD_2Cl_2$) 162.48 (C4), 159.29 (Ar), 150.94 (C2), 145.27 (Ar), 141.81(C-6), 136.21 (Ar), 136.06 (Ar), 130.64 (Ar), 130.62 (Ar), 128.57 (Ar), 128.50 (Ar), 127.47 (Ar), 113.76 (Ar), 104.21 (C-5), 87.33 (Ar), 86.82 (C4'), 85.89 (C-1'), 72.82 (C-3'), 64.24 (C-5'), 41.55 (C2'), 7.44 (Se$CH_3$); HRMS (ESI-TOF): Molecular formula $C_{36}H_{34}N_2O_7Se$ $[M]^+$: 684.1609 (calc. 684.1531).

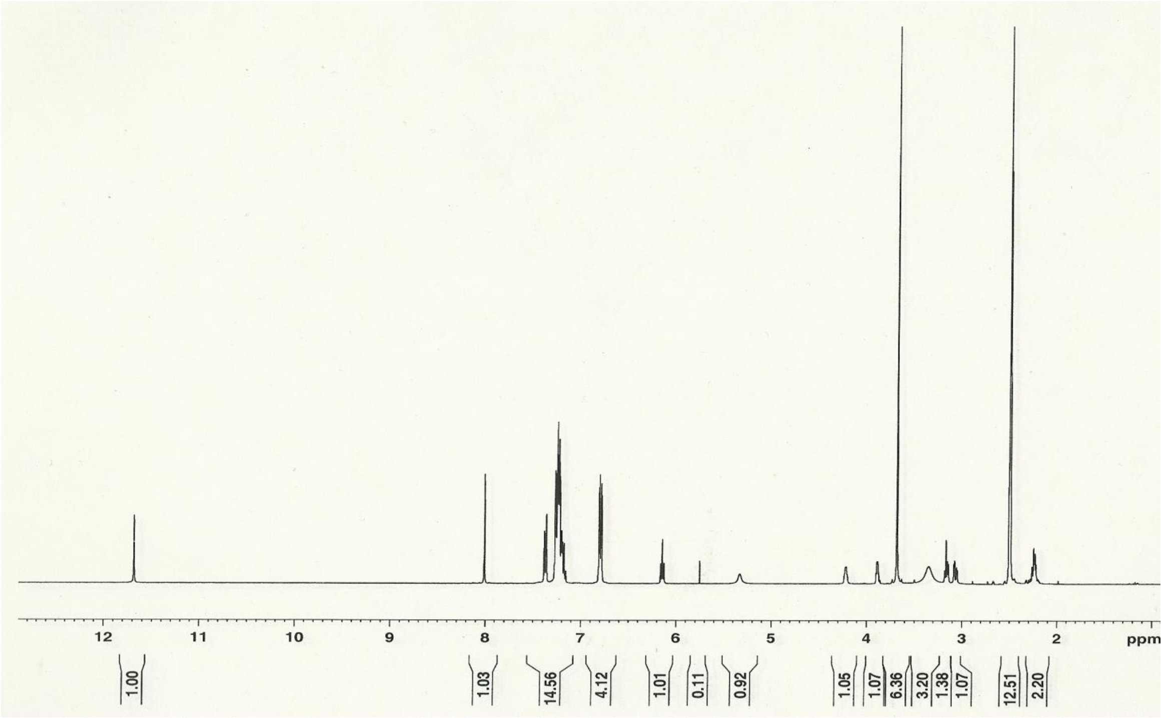

Figure S18. $^{1}$H NMR of compound **6** (X=PhSe).

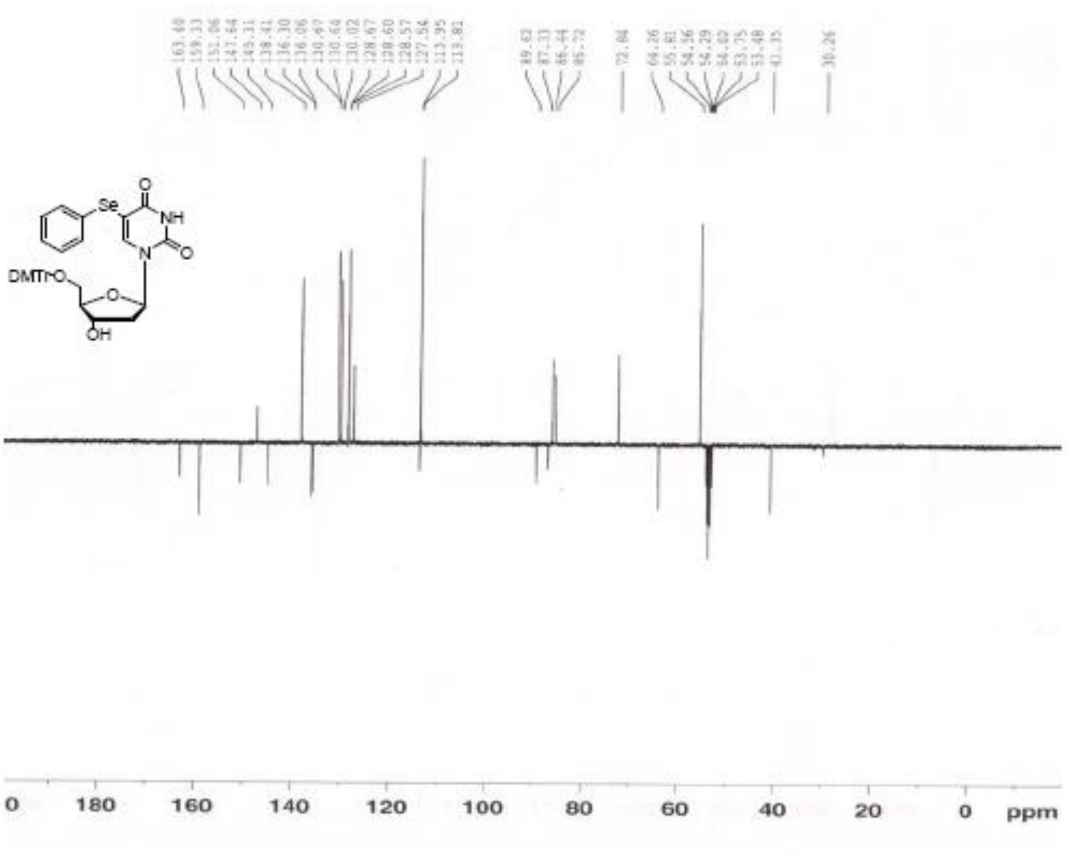

Figure S19. $^{13}$C NMR of compound **6** (X=PhSe).

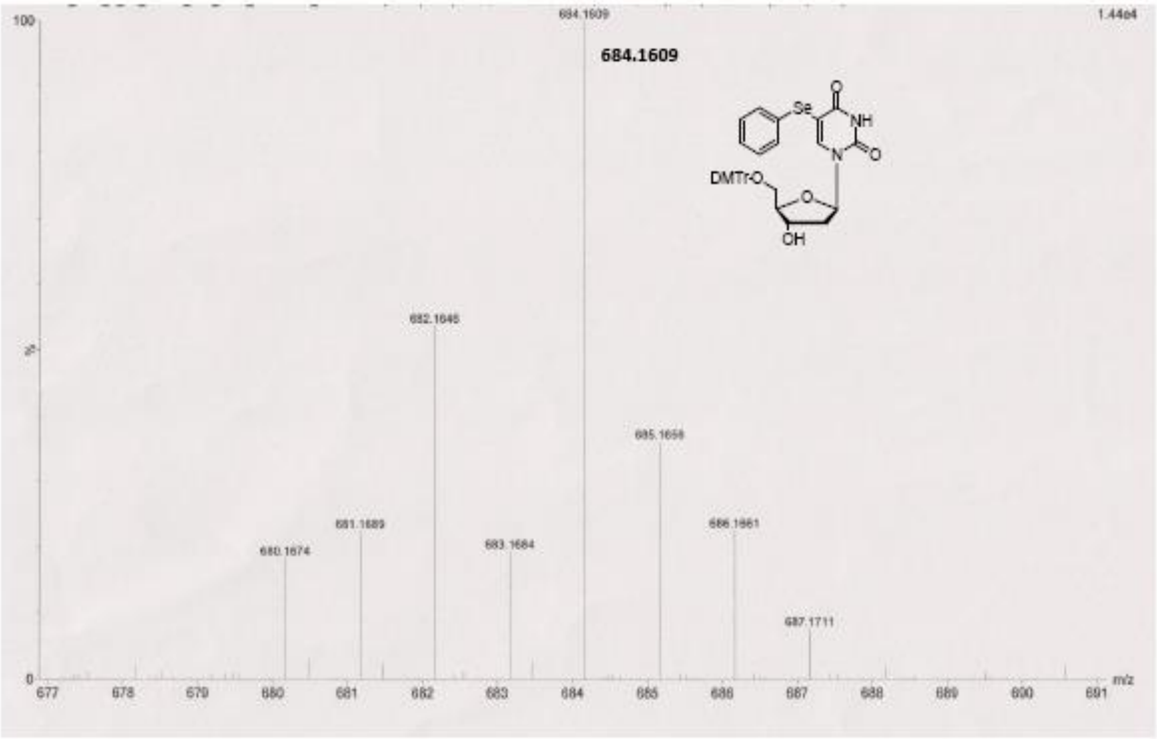


Figure S20. HRMS of compound **6** (X=PhSe).

**1-[2'-deoxy-3'-O-(2-cyanoethyl-*N,N*-diisopropylamino)-phosphoramidite-5'-O-(4,4-dimethoxytrityl-[D-erythro-ribofuranosyl]-5-phenylselenylthymidine (7, X=PhSe).** Disopropyl-ethylamine (20 mL, 0.12 mmol) was added to a solution of 6 (0.3 g, 0.48 mmol), 5-benzylthiotetrazole (45.4 mg, 0.24 mmol) and 2-cyanoethyl-*N,N,N,N*-tetraisopropyl phosphane (288 mg, 0.96 mmol) in dry $CH_2Cl_2$ (15 mL) at 0 $^oC$. The mixture was stirred for 2 h at room temperature then slowly poured into pentane (100 mL). The produced white precipitate was filtered off, dried under high vacuum and directly applied in solid phase synthesis without further purification.

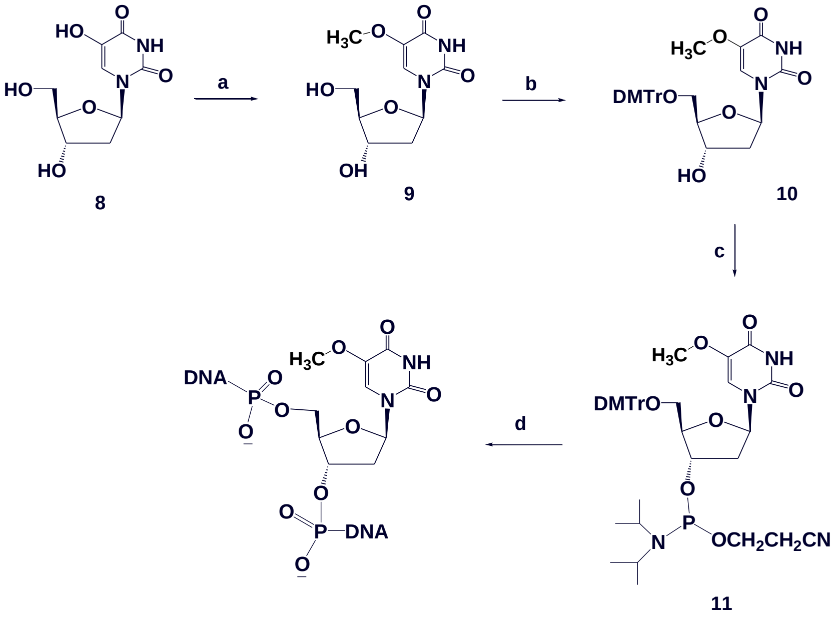


Scheme S2: Synthesis of 5-Methoxy-T DNAs. Reagents and conditions: a) $CH_3I$, NaOH, MeOH, $H_2O$, rt; b) DMTr-Cl, Pyridine, DMAP, rt, 85%; c) i-$Pr_2NP(Cl)CH_2CH_2CN$, DIEA, $CH_2Cl_2$, 1 h, rt, 82%; d) solid phase synthesis.

**2'-deoxy-5-methoxy-thymidine (9)** 5-hydroxy-2'-deoxyuridine (1.288 g, 5 mmol) was dissolved in a solution of $H_2O$ (20 mL) and methonal (10 mL). NaOH solution 5 mL (1 M, 5 mmol) and $CH_3I$ (1.24 mL, 20 mmol) were injected, before the mixture was stirred at room temperature for 4 days. The solvent was evaporated and the residue was purified by silica gel column chromatography (eluate 5%-10% MeOH in $CHCl_3$) to give (0.58 mg, 92%) of **9**: $^1$H-NMR ($CD_2Cl_2$) 9.39 (1H, s, NH, exchanged with $D_2O$), 7.90 (1H, s, H-6), 7.46-7.21 (9H, m, Ar), 6.87-6.84(4H, m, Ar), 6.29 (1H, dd, H-1', J = 6.4, J = 7.6 Hz), 4.43 (1H, m, H-3'), 4.05 (1H, m, H-4'), 3.68 (6H, 2 s, OMe), 3.30 (1H, dd, H5'a, J = 3.8, J = 10.5 Hz), 3.24 (1H, dd, H5'b, J = 3.6, J = 10.5 Hz), 2.58 (1H, d, 3'-OH), 2.42 (1H, ddd, H-2'a, J = 3.8, J = 7.7, J = 10.8 Hz), 2.36 (1H, m, H-2'b), 1.90 (3H, s, $SeCH_3$); 13C-NMR ($CD_2Cl_2$) 162.48 (C4), 159.29 (Ar), 150.94 (C2), 145.27 (Ar), 141.81(C-6), 136.21 (Ar), 136.06 (Ar), 130.64 (Ar), 130.62 (Ar), 128.57 (Ar), 128.50 (Ar), 127.47 (Ar), 113.76 (Ar), 104.21 (C-5), 87.33 (Ar), 86.82 (C4'), 85.89 (C-1'), 72.82 (C-3'), 64.24 (C-5'), 41.55 (C2'), 7.44 ($SeCH_3$); HRMS (ESI-TOF): Molecular formula $C_{10}H_{14}N_2O_6$ $[M+H]^+$: 260.2 (calc. 259.1).

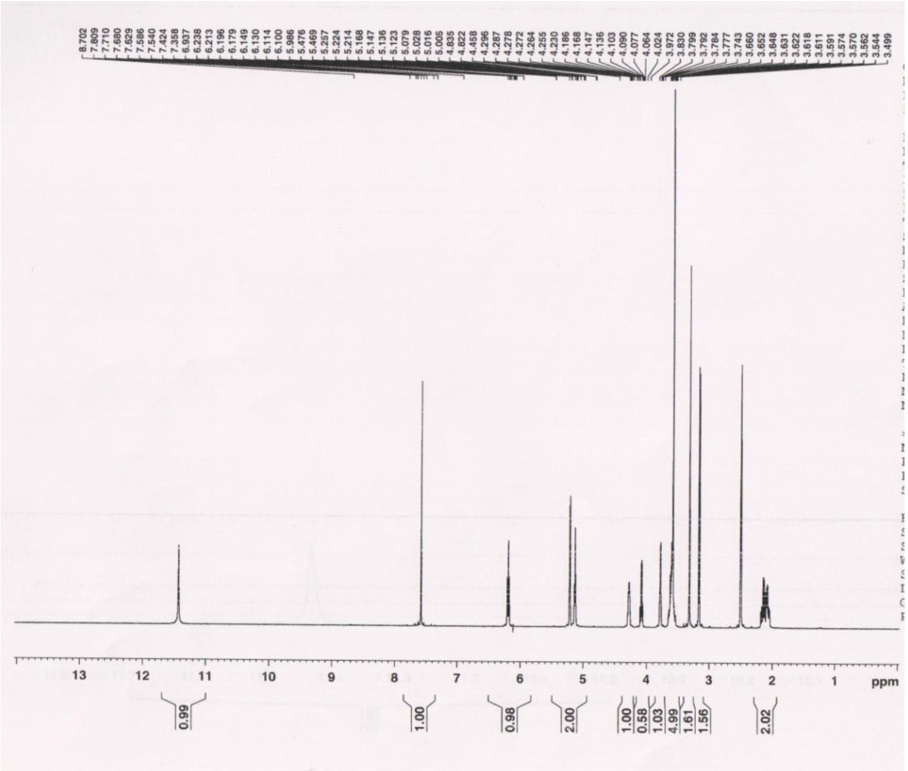

Figure S21. $^{1}$H NMR of compound **9**.

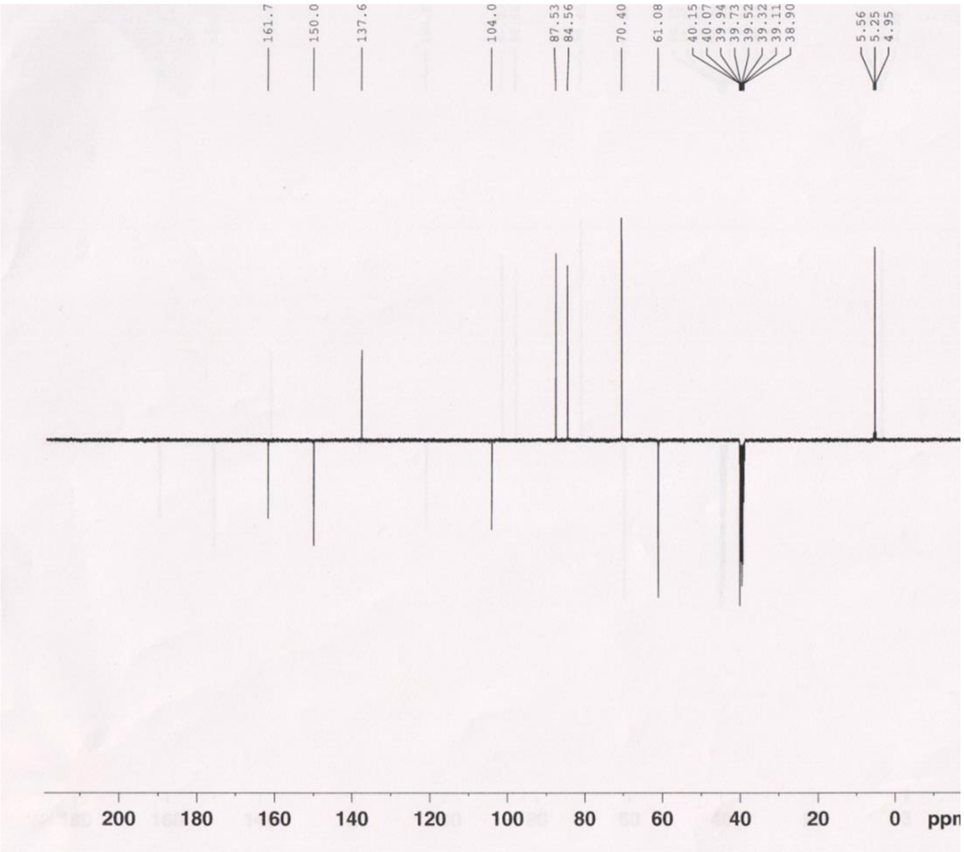

Figure S22. $^{13}$C NMR of compound **9**.

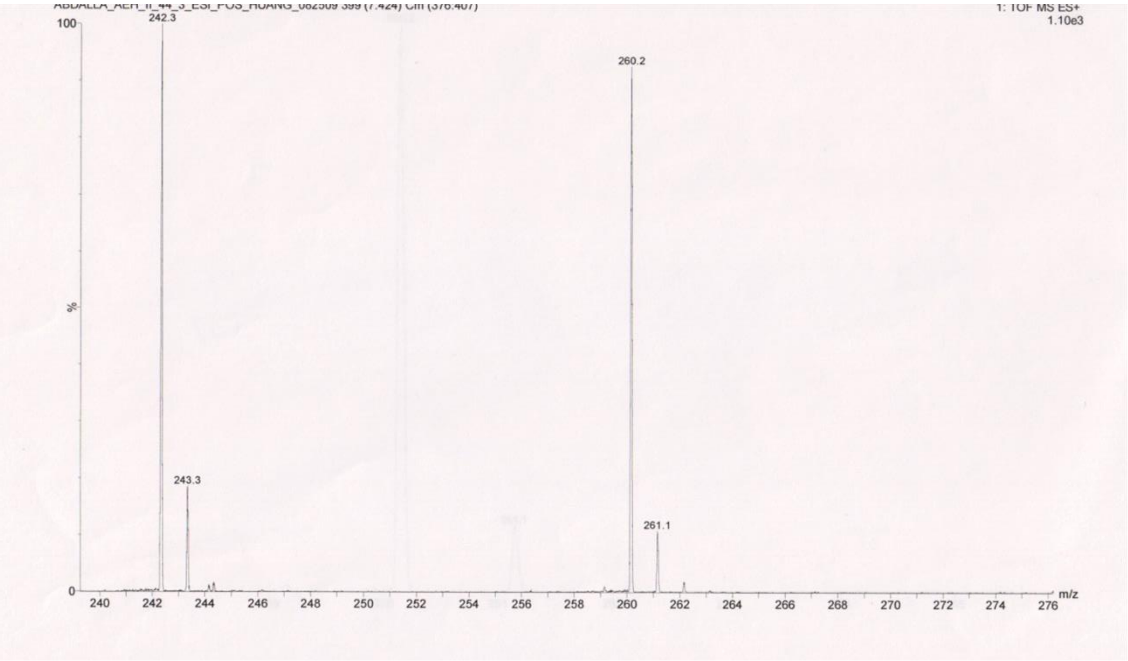


Figure S23. MS of compound **9**.

**5'-(*O*-4,4'-dimethoxytrityl)-2'-deoxy-5-methoxy-thymidine (10)** 4,4'-Dimethoxytrityl chloride (0.95 g, 2.79 mmol) was added to a solution of 2-thiothymidine (0.6 g, 2.32 mmol) and DMAP (10 mg) in dry pyridine (6 mL) at O °C. The mixture was stirred for 6 h at rt, and then MeOH (0.5 mL) was added to the mixture. The solvents were evaporated under reduced pressure and the residue was partitioned between EtOAc and H2O. The organic phase was dried (MgSO4) and evaporated. The residue was purified by flash column chromatography ($SiO_2$, pre-equalized with 1% pyridine in $CH_2Cl_2$: 10% EtOAc in $CH_2Cl_2$) gave (1.1 g, 85%) of **10** as a colorless foam. $^1$H-NMR ($CD_2Cl_2$) 9.39 (1H, s, NH, exchanged with $D_2O$), 7.90 (1H, s, H-6), 7.46-7.21 (9H, m, Ar), 6.87-6.84(4H, m, Ar), 6.29 (1H, dd, H-1', J = 6.4, J = 7.6 Hz), 4.43 (1H, m, H-3'), 4.05 (1H, m, H-4'), 3.68 (6H, 2 s, OMe), 3.30 (1H, dd, H5'a, J = 3.8, J = 10.5 Hz), 3.24 (1H, dd, H5'b, J = 3.6, J = 10.5 Hz), 2.58 (1H, d, 3'-OH), 2.42 (1H, ddd, H-2'a, J = 3.8, J = 7.7, J = 10.8 Hz), 2.36 (1H, m, H-2'b), 1.90 (3H, s, $SeCH_3$); 13C-NMR ($CD_2Cl_2$) 162.48 (C4), 159.29 (Ar), 150.94 (C2), 145.27 (Ar), 141.81(C-6), 136.21 (Ar), 136.06 (Ar), 130.64 (Ar), 130.62 (Ar), 128.57 (Ar), 128.50 (Ar), 127.47 (Ar), 113.76 (Ar), 104.21 (C-5), 87.33 (Ar), 86.82 (C4'), 85.89 (C-1'), 72.82 (C-3'), 64.24 (C-5'), 41.55 (C2'), 7.44 ($SeCH_3$); HRMS (ESI-TOF): Molecular formula $C_{31}H_{32}N_2O_8$ $[M+Na]^+$: 583.1883 (calc. 583.2159).

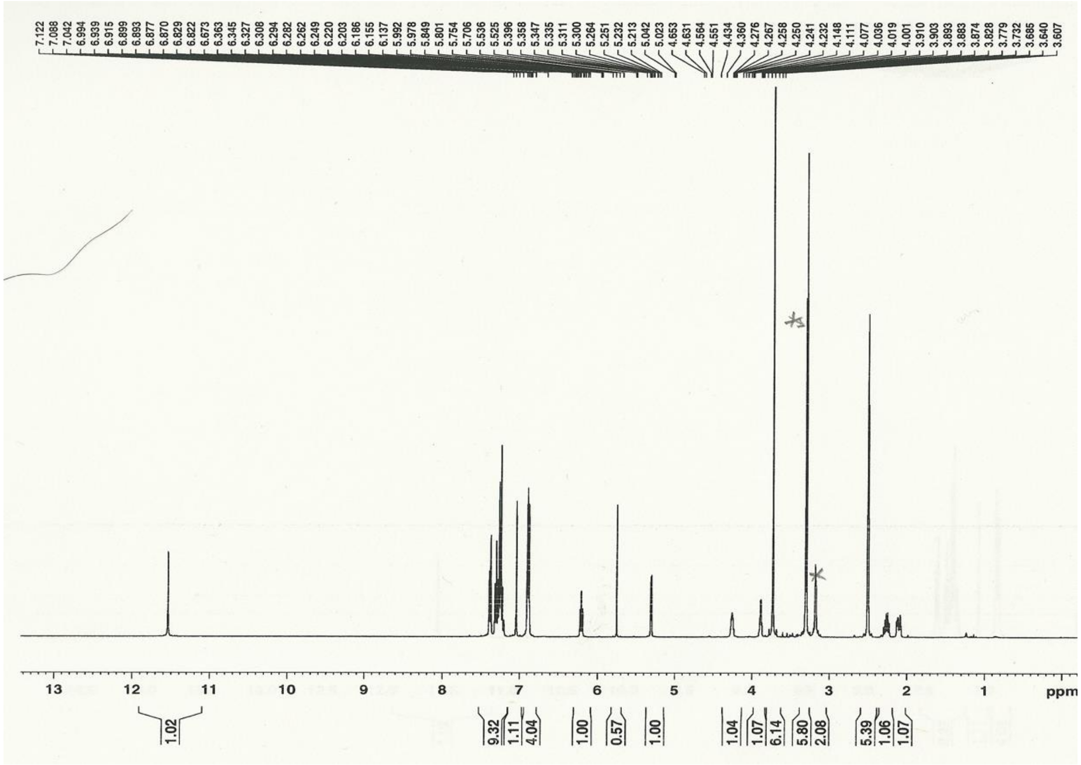

Figure S24. $^{1}$H NMR of compound **10.**

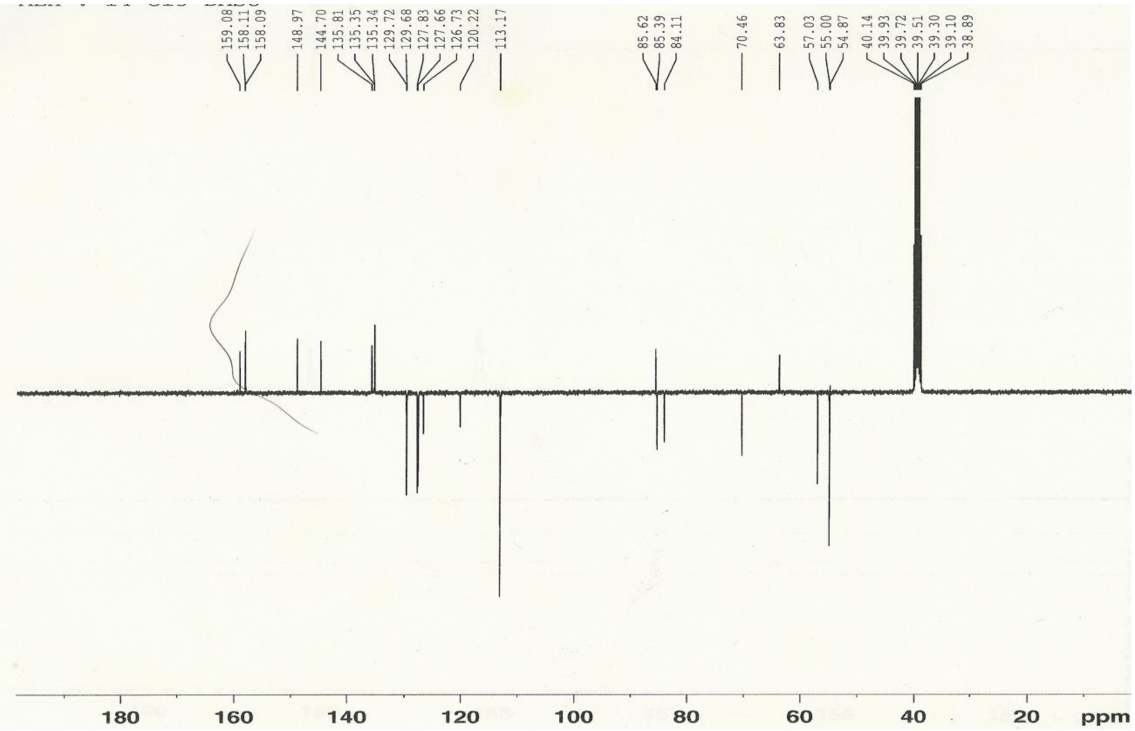

Figure S25. $^{13}$C NMR of compound **10.**

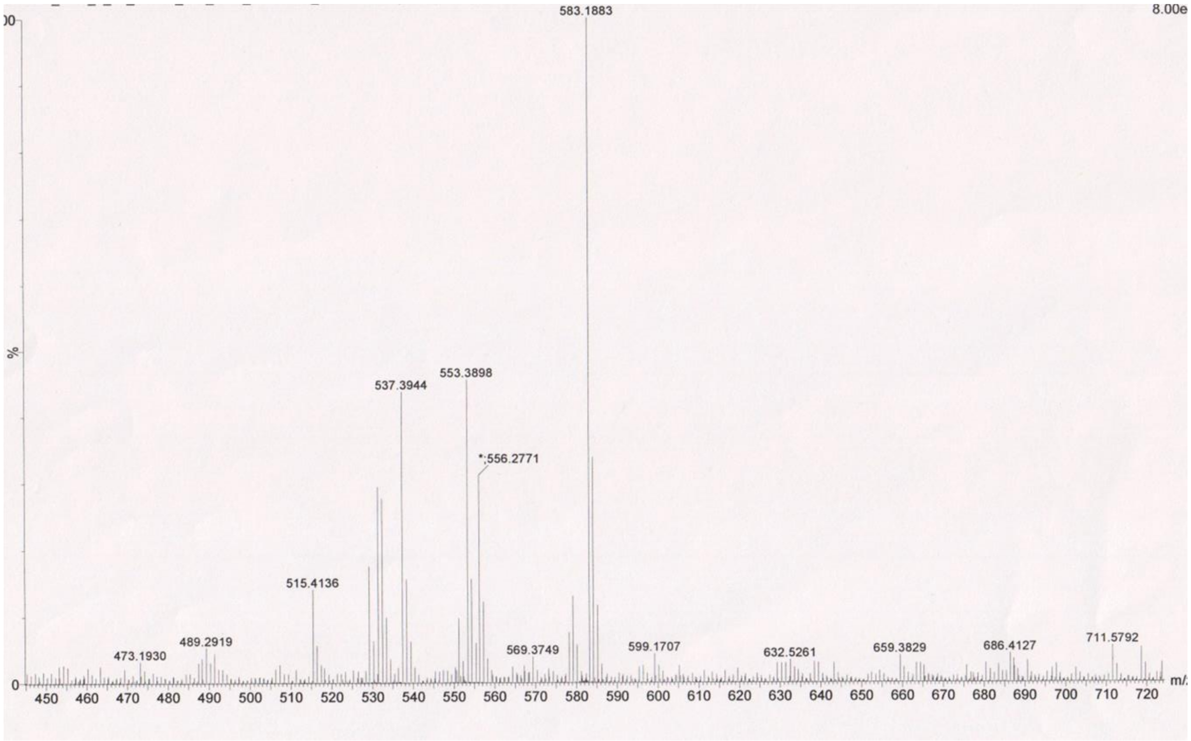


Figure S26. HRMS of compound **10.**

**1-[2'-deoxy-3'-*O*-(2-cyanoethyl-*N,N*-diisopropylamino)-phosphoramidite-5'-*O*-(4,4-dimethoxytrityl-[D-erythro-ribofuranosyl]-5-methoxythymidine (11)** Disopropyl-ethylamine (20 mL, 0.12 mmol) was added to a solution of **10** (0.3 g, 0.48 mmol), 5-benzylthiotetrazole (45.4 mg, 0.24 mmol) and 2-cyanoethyl-*N,N,N,N*-tetraisopropyl phosphane (288 mg, 0.96 mmol) in dry $CH_2Cl_2$ (15 mL) at 0 $^oC$. The mixture was stirred for 2 h at room temperature then slowly poured into pentane (100 mL). The produced white precipitate was filtered off, dried under high vacuum and directly applied in solid phase synthesis without further purification.

#### 3. HPLC and MALDI-TOF MS spectra of synthetic DNA oligonucleotides.

##### Synthesis of the 5-chalcogen-T functionalized DNA oligonucleotides

All the DNA oligonucleotides were chemically synthesized in a 1.0 μmol scale using an ABI392 or ABI3400 DNA/RNA Synthesizer. The regular nucleoside phosphoramidite reagents were used in this work (Glen Research). The concentration of the 5-chalcogen-T phosphoramidite was identical to that of the conventional ones (0.1 M in acetonitrile). Coupling was carried out using a 5-(benzylmercapto)-1H-tetrazole (5-BMT) solution (0.25 M) in acetonitrile. The coupling time was 25 seconds for both native and modified samples. 3% trichloroacetic acid in methylene chloride was used for the 5'-detritylation. Synthesis were performed on control pore glass (CPG-500) immobilized with the appropriate nucleoside through a succinate linker. All the oligonucleotides were prepared with DMTr-on form. After synthesis, the DNA oligonucleotides were cleaved from the solid support and fully deprotected by the treatment of ammonium (conc.) for 8 hr at 55 $^oC$. The 5'-DMTr deprotection was performed in a 3% trichloroacetic acid solution for 2 min, followed by neutralization to pH 7.0 with a freshly made aqueous solution of triethylamine (1.1 M) and extraction by petroleum ether to remove DMTr-OH.

Table S1. MALDI-TOF MS data of the 5-chalcogen-T DNAs.

| entry | Chalcogen-oligonucleotide | measured (calcd.) m/z |
|---|---|---|
| 1 | 5'-GdU$_{2'\text{-Se}}$G-$^{5CH3Se}$T-ACAC-3'<br>$C_{78}H_{99}N_{30}O_{46}P_7Se_2$ | [M+H]$^+$: 2569.0 (2568.6) |
| 2 | 5'-ATGG-$^{5CH3Se}$T-GCTC-3'<br>$C_{88}H_{112}N_{32}O_{54}P_8Se$ | [M]$^+$: 2808.7 (2808.4) |
| 3 | 5'-CTCCCA-$^{5CH3Se}$T-CC-3'<br>$C_{84}H_{111}N_{27}O_{53}P_8Se$ | [M+H]$^+$: 2674.5 (2674.6) |
| 4 | 5'-CTTCT-$^{5CH3Se}$T-GTCCG-3'<br>$C_{106}H_{138}N_{32}O_{69}P_{10}Se$ | [M]$^+$: 3352.4 (3353.3) |
| 5 | 5'-GdU$_{2'\text{-Se}}$G-$^{5CH3S}$T-ACAC-3'<br>$C_{78}H_{99}N_{30}O_{46}P_7SSe$ | [M]$^+$: 2521.1 (2520.6) |
| 6 | 5'-ATGG-$^{5CH3S}$T-GCTC-3'<br>$C_{88}H_{112}N_{32}O_{54}P_8S$ | [M]$^+$: 2760.8 (2761.7) |
| 7 | 5'-CTCCCA-$^{5CH3S}$T-CC-3'<br>$C_{84}H_{111}N_{27}O_{53}P_8S$ | [M]$^+$: 2626.7 (2626.6) |
| 8 | 5'-CTTCT-$^{5CH3S}$T-GTCCG-3'<br>$C_{106}H_{138}N_{32}O_{69}P_{10}S$ | [M]$^+$: 3305.8 (3306.3) |
| 9 | 5'-GdU$_{2'\text{-Se}}$G-$^{5CH3O}$T-ACAC-3'<br>$C_{78}H_{99}N_{30}O_{47}P_7Se$ | [M+H]$^+$: 2505.8 (2507.0) |
| 10 | 5'-ATGG-$^{5CH3O}$T-GCTC-3'<br>$C_{88}H_{112}N_{32}O_{55}P_8$ | [M+Na]$^+$: 2871.9 (2869.7) |
| 11 | 5'-CTCCCA-$^{5CH3O}$T-CC-3'<br>$C_{84}H_{111}N_{27}O_{54}P_8$ | [M+H]$^+$: 2611.0 (2611.6) |
| 12 | 5'-CTTCT-$^{5CH3O}$T-GTCCG-3'<br>$C_{106}H_{138}N_{32}O_{70}P_{10}$ | [M+H]$^+$: 3290.9 (3291.3) |
| 13 | 5'-GdU$_{2'\text{-Se}}$G-$^{5PhSe}$T-ACAC-3'<br>$C_{83}H_{101}N_{30}O_{46}P_7Se_2$ | [M]+: 2627.9 (2629.6) |
| 14 | 5'-ATGG-$^{5PhSe}$T-GCTC-3'<br>$C_{93}H_{114}N_{32}O_{54}P_8Se$ | [M+H]$^+$: 2873.6 (2872.7) |
| 15 | 5'-CTCCCA-$^{5PhSe}$T-CC-3'<br>$C_{89}H_{113}N_{27}O_{53}P_8Se$ | [M+H]$^+$: 2738.7 (2736.6) |
| 16 | 5'-CTTCT-$^{5PhSe}$T-GTCCG-3'<br>$C_{111}H_{140}N_{32}O_{69}P_{10}Se$ | [M+H]$^+$: 3417.9 (3416.3) |

#### HPLC analysis, purification and characterization

The DNA oligonucleotides were analyzed and purified by reversed-phase high performance liquid chromatography (RP-HPLC) both DMTr-on and DMTr-off. Purification was carried out using a 21.2 x 250 mm Zorbax, RX-C8 column at a flow rate of 6 mL/min. Buffer A consisted of 20 mM triethylammonium acetate (TEAAc, pH 7.1), while buffer B contained 50% acetonitrile in 20 mM triethylammonium acetate (TEAAc, pH 7.1). The DMTr-on oligonucleotides were eluded and purified in a linear gradient reaching 100% buffer B in 20 min, while the DMTr-off oligonucleotides were eluded and purified in a linear gradient reaching 70% buffer B in 20 min. The collected fractions were lyophilized and the purified compounds were re-dissolved in water. The pH was adjusted to 7.0 after the final purification of the Se-oligonucleotides without the DMTr group. Similarly, analysis was performed on a Zorbax SB-C18 column (4.6 x 250 mm) at a flow of 1.0 mL/min, in a linear gradient reaching 70% buffer B in 20 min. MALDI-TOF MS is used to characterize all Se-DNA samples.

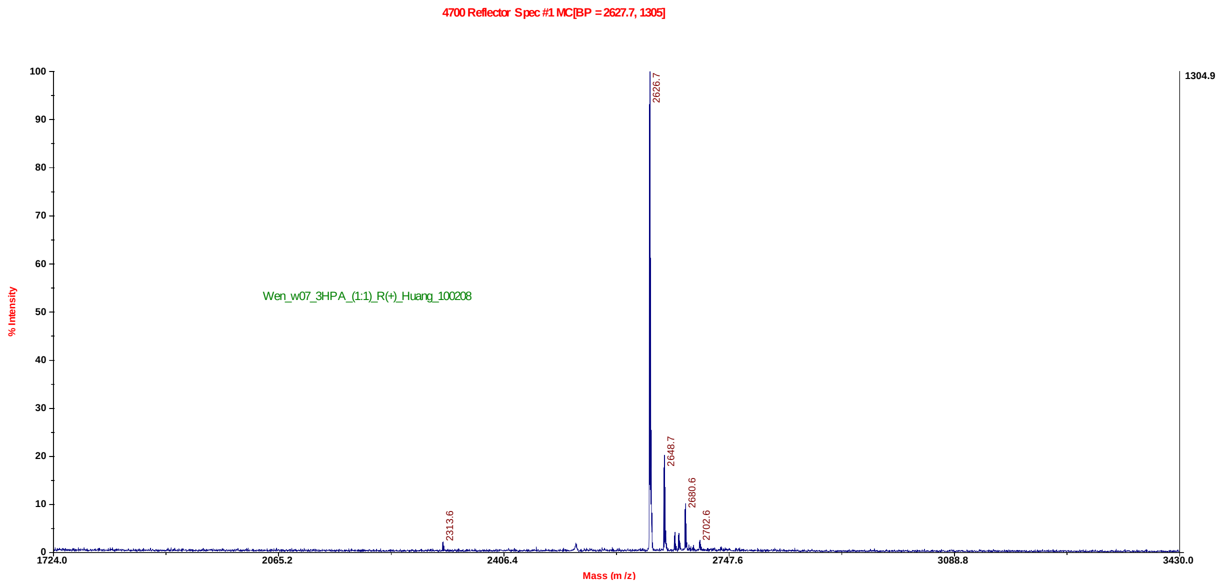


**Figure S27.** MALDI-TOF MS spectrum of 5'-CTCCCA-$^{5\text{-}CH3S}$T-CC-3', Molecular formula: $C_{84}H_{111}N_{27}O_{53}P_8S$, $[M]^+$: 2626.7 (calc. 2626.6).

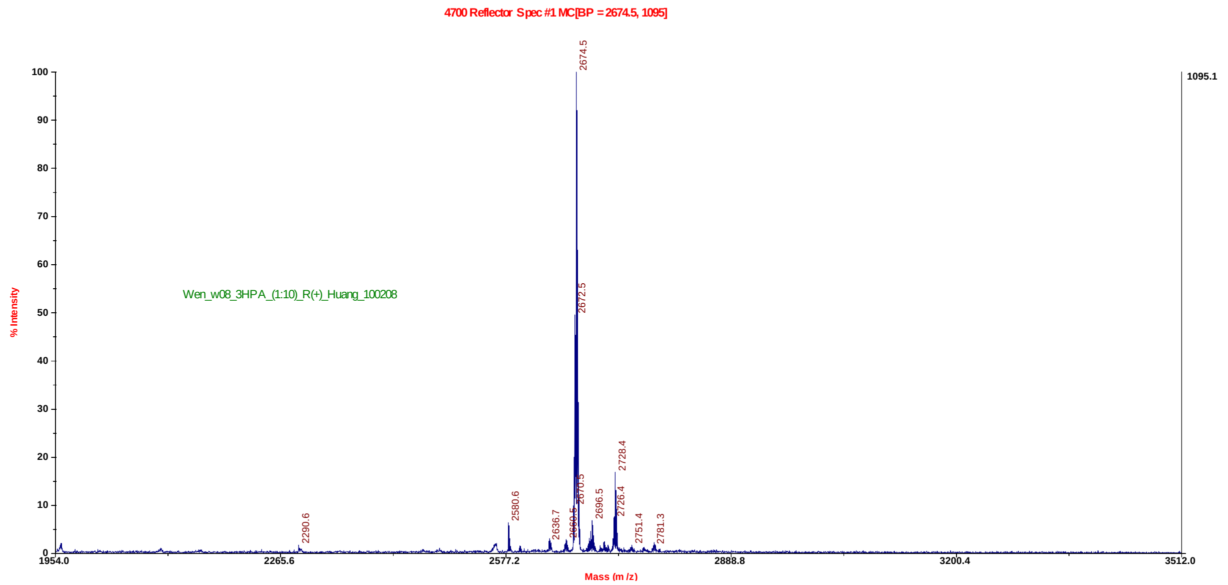


**Figure S28**. MALDI-TOF MS spectrum of 5'-CTCCCA-$^{5\text{-}CH3Se}$T-CC-3', Molecular formula: $C_{84}H_{111}N_{27}O_{53}P_{8}Se$, $[M+H]^{+}$: 2674.5 (calc. 2674.6).

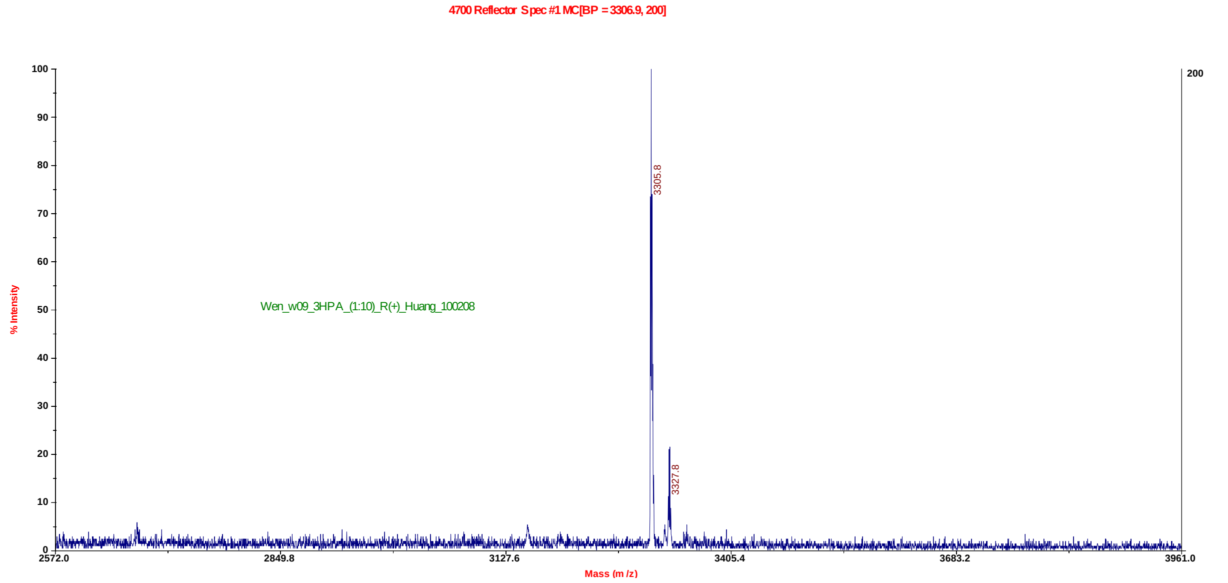


**Figure S29**. MALDI-TOF MS spectrum of 5'-CTTCT-$^{5\text{-}CH3S}$T-GTCCG-3', Molecular formula: $C_{106}H_{138}N_{32}O_{69}P_{10}S$ $[M]^{+}$: 3305.8 (calc. 3306.3).

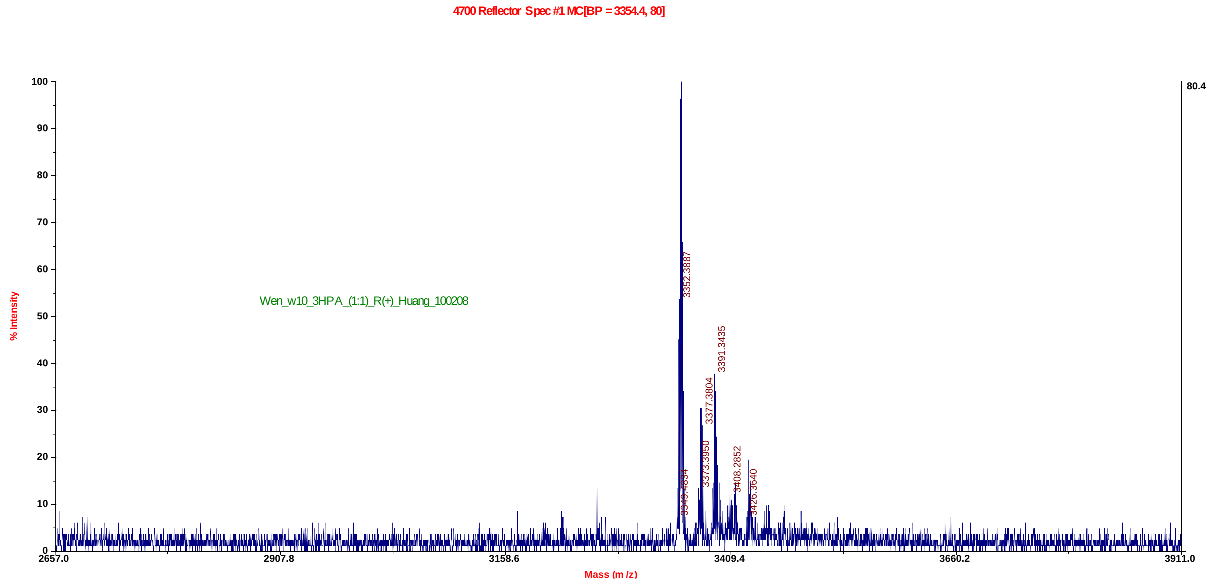


**Figure S30**. MALDI-TOF MS spectrum of 5'-CTTCT-$^{5\text{-}CH3Se}$T-GTCCG-3', Molecular formula: $C_{106}H_{138}N_{32}O_{69}P_{10}Se$ $[M]^+$: 3352.4 (calc. 3353.3).

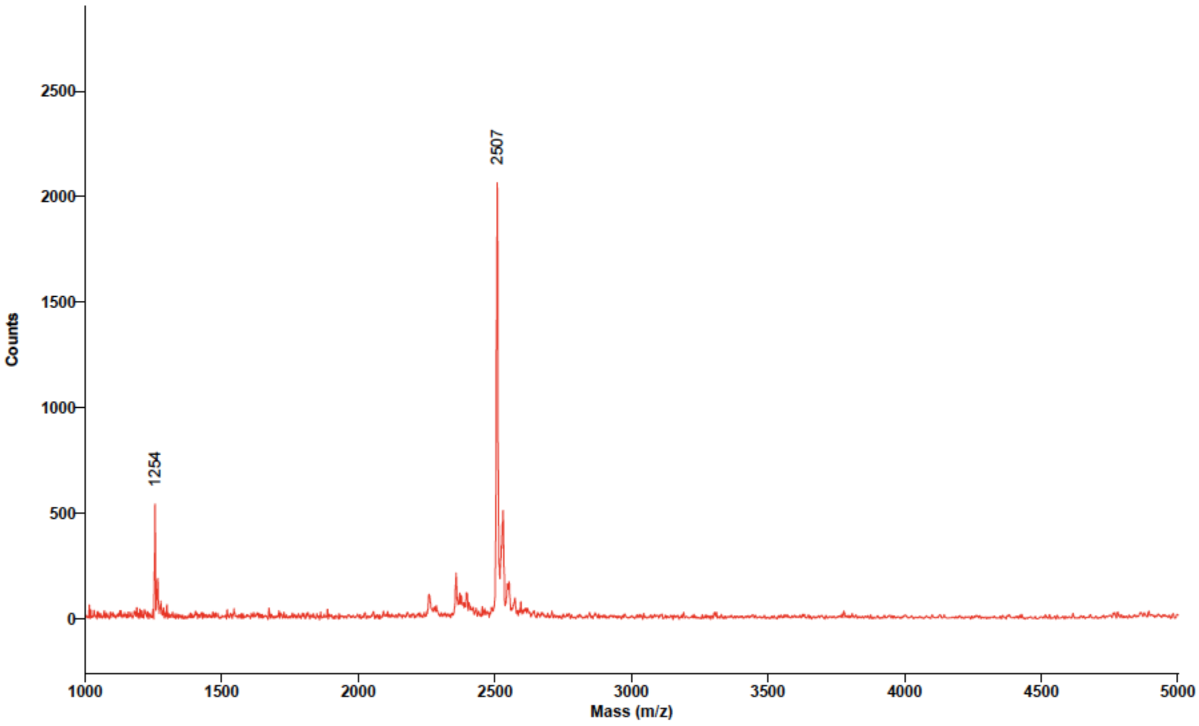


**Figure S31**. MALDI-TOF MS spectrum of 5'-GdU$_{2'\text{-}Se}$G-$^{5\text{-}CH3O}$T-ACAC-3', Molecular formula: $C_{78}H_{99}N_{30}O_{47}P_{7}Se$ $[M+H]^+$: 2507 (calc. 2505.8).

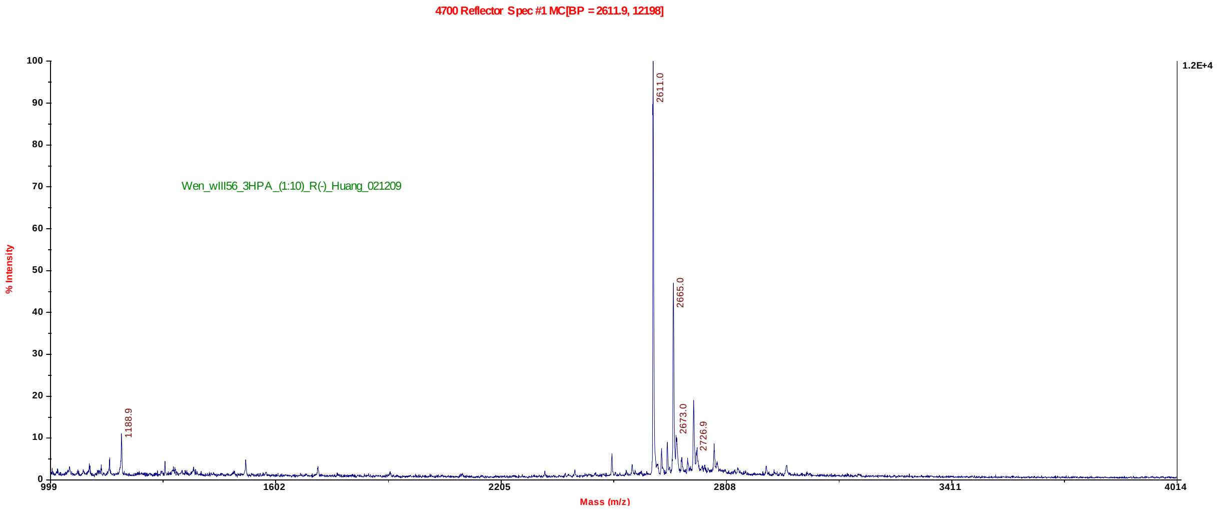


**Figure S32**. MALDI-TOF MS spectrum of 5'-CTCCCA-$^{5\text{-}CH3O}$T-CC-3', Molecular formula: $C_{84}H_{111}N_{27}O_{54}P_{8}$, [M+H]$^+$: 2611.0 (calc. 2611.6).

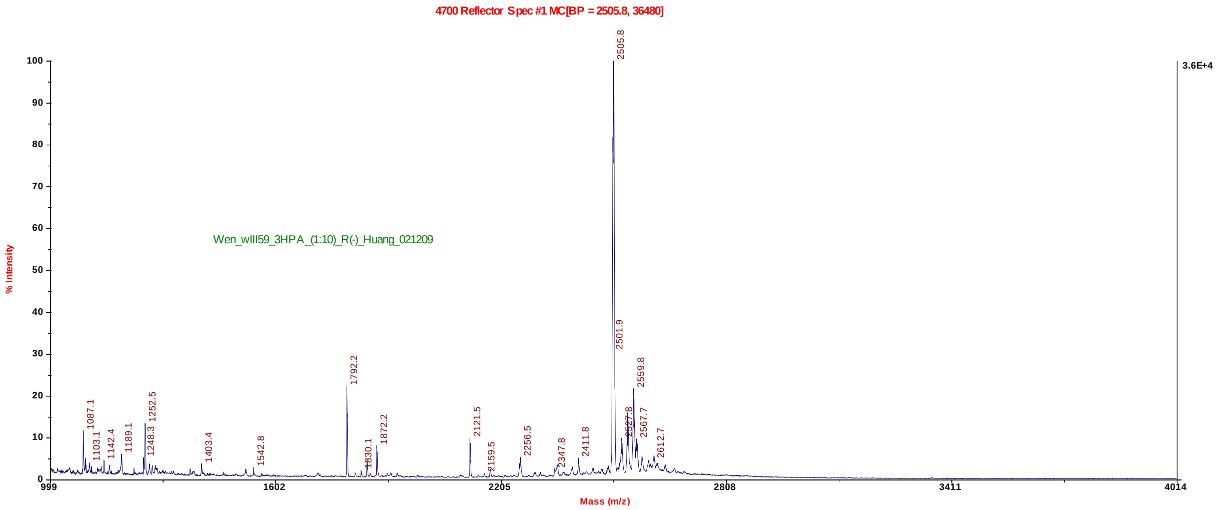


**Figure S33**. MALDI-TOF MS spectrum of 5'-GdU$_{2'\text{-}Se}$G-$^{5\text{-}CH3O}$T-ACAC-3', Molecular formula: $C_{78}H_{99}N_{30}O_{47}P_{7}Se$, [M+H]$^+$: 2505.8 (calc. 2507.0).

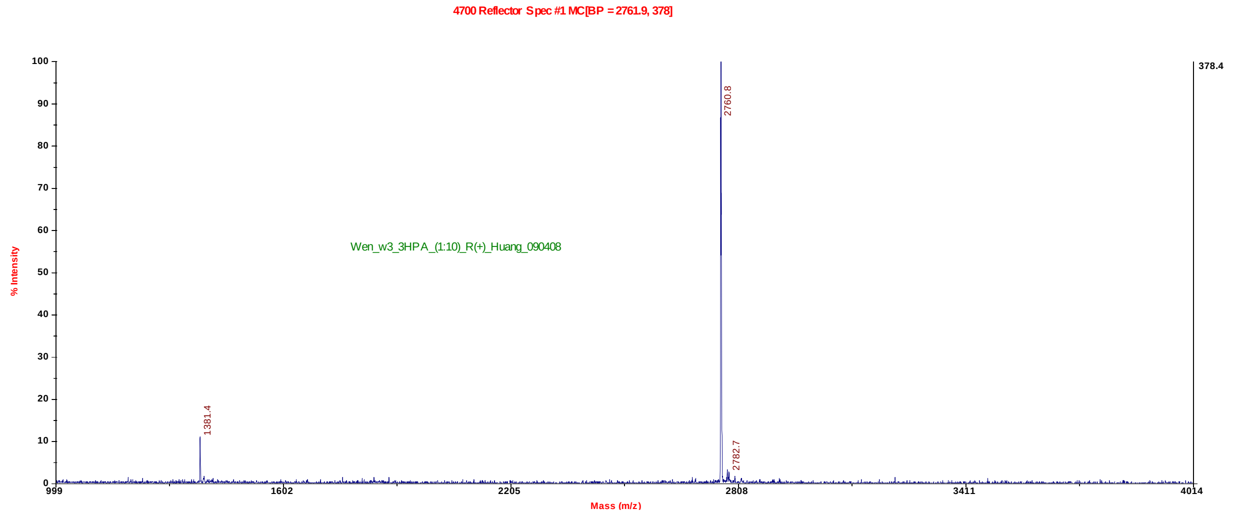


**Figure S34**. MALDI-TOF MS spectrum of 5'-ATGG-$^{5\text{-}CH3S}$T-GCTC-3', Molecular formula: $C_{88}H_{112}N_{32}O_{54}P_8S$, $[M]^+$: 2760.8 (calc. 2761.7).

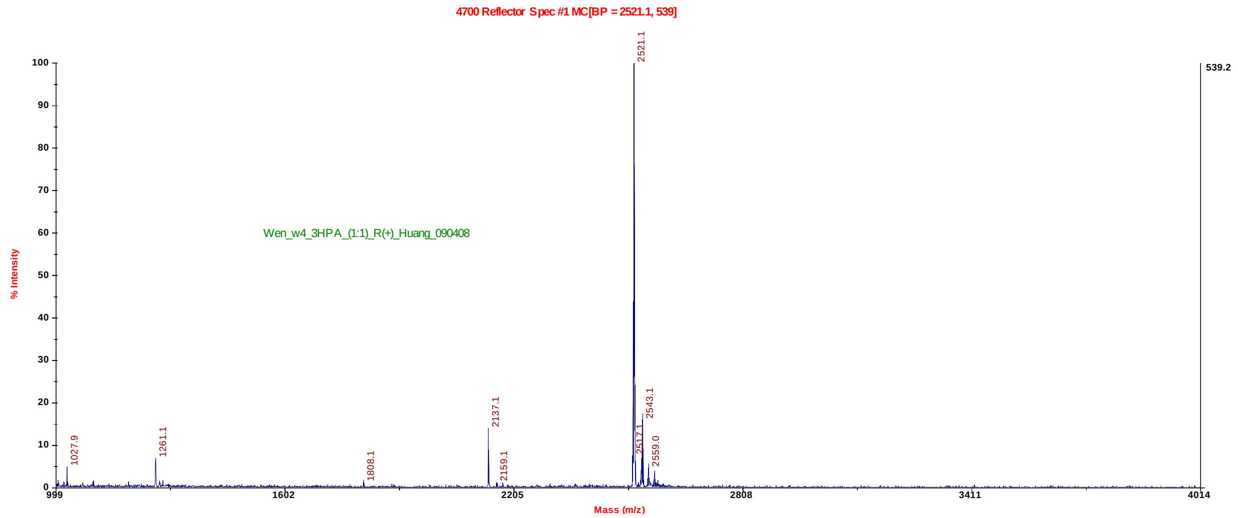


**Figure S35.** MALDI-TOF MS spectrum of 5'-GdU$_{2'\text{-}Se}$G-$^{5\text{-}CH3S}$T-ACAC-3', Molecular formula: $C_{78}H_{99}N_{30}O_{46}P_7SSe$, $[M]^+$: 2521.1 (calc. 2520.6).

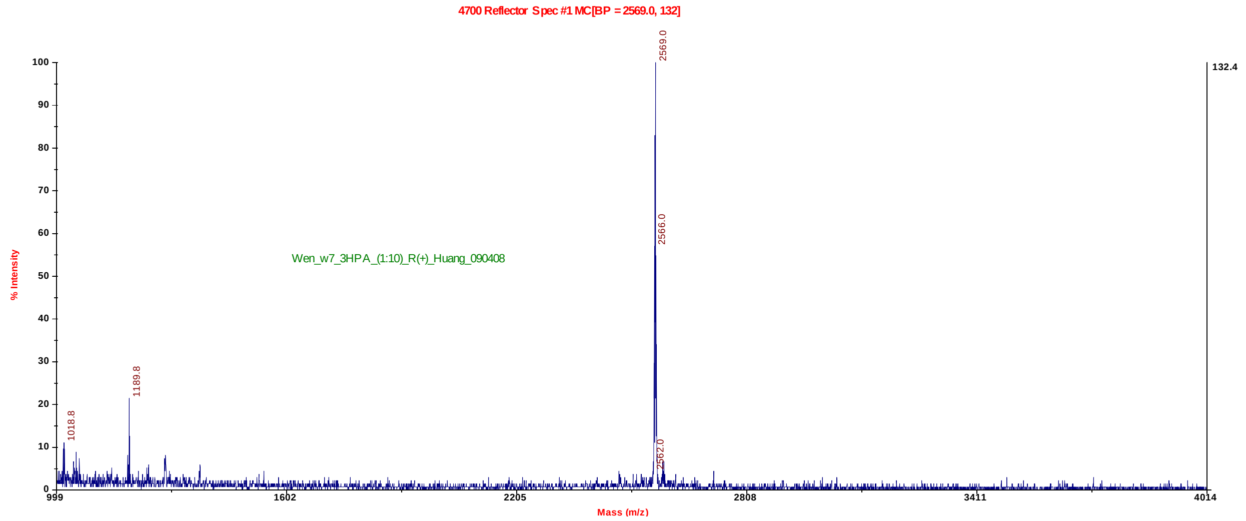


**Figure S36**. MALDI-TOF MS spectrum of 5'-GdU$_{2'\text{-}Se}$G-$^{5\text{-}CH3Se}$T-ACAC-3', Molecular formula: $C_{78}H_{99}N_{30}O_{46}P_7Se_2$, [M+H]$^+$: 2569.0 (calc. 2568.6).

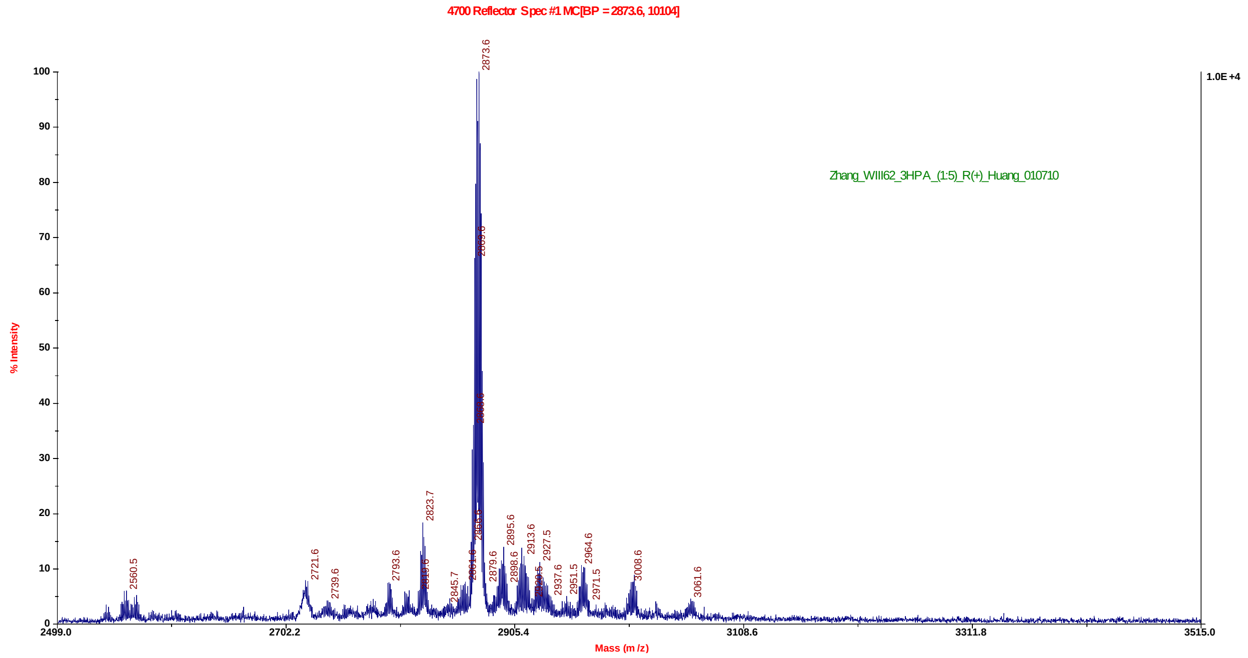


**Figure S37**. MALDI-TOF MS spectrum of 5'-ATGG-$^{5\text{-}PhSe}$T-GCTC-3', Molecular formula: $C_{93}H_{114}N_{32}O_{54}P_8Se$, [M+H]$^+$: 2873.6 (calc. 2872.7).

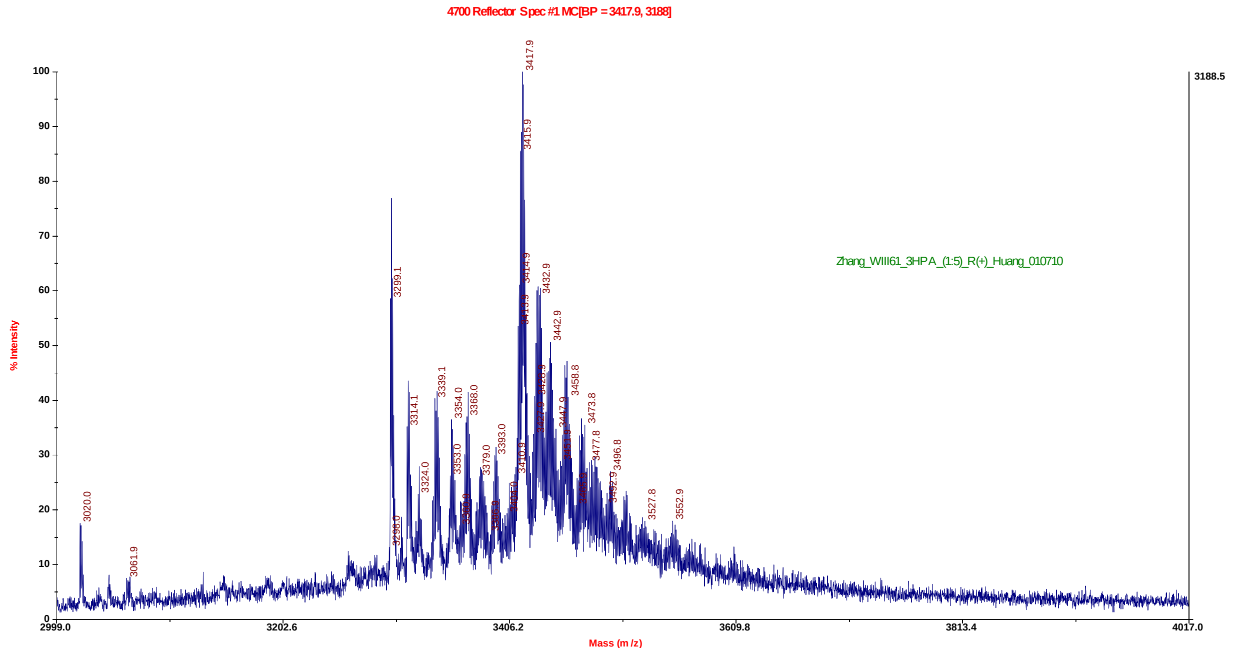


**Figure S38**. MALDI-TOF MS spectrum of 5'-CTTCT-$^{5\text{-PhSe}}$T-GTCCG-3', Molecular formula: $C_{111}H_{140}N_{32}O_{69}P_{10}Se$ $[M+H]^+$: 3417.9.6 (calc. 3416.3).

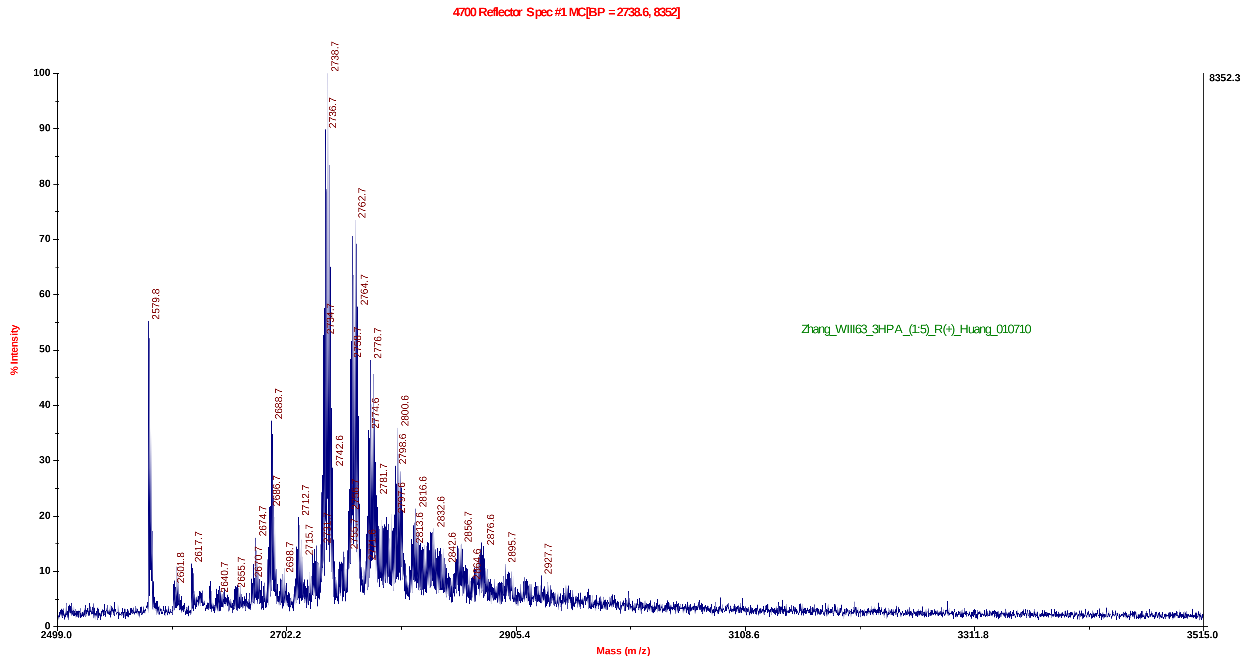


**Figure S39**. MALDI-TOF MS spectrum of 5'-CTCCCA-$^{5\text{-PhSe}}$T-CC-3', Molecular formula: $C_{89}H_{113}N_{27}O_{53}P_8Se$, $[M+H]^+$: 2738.7 (calc. 2736.6).

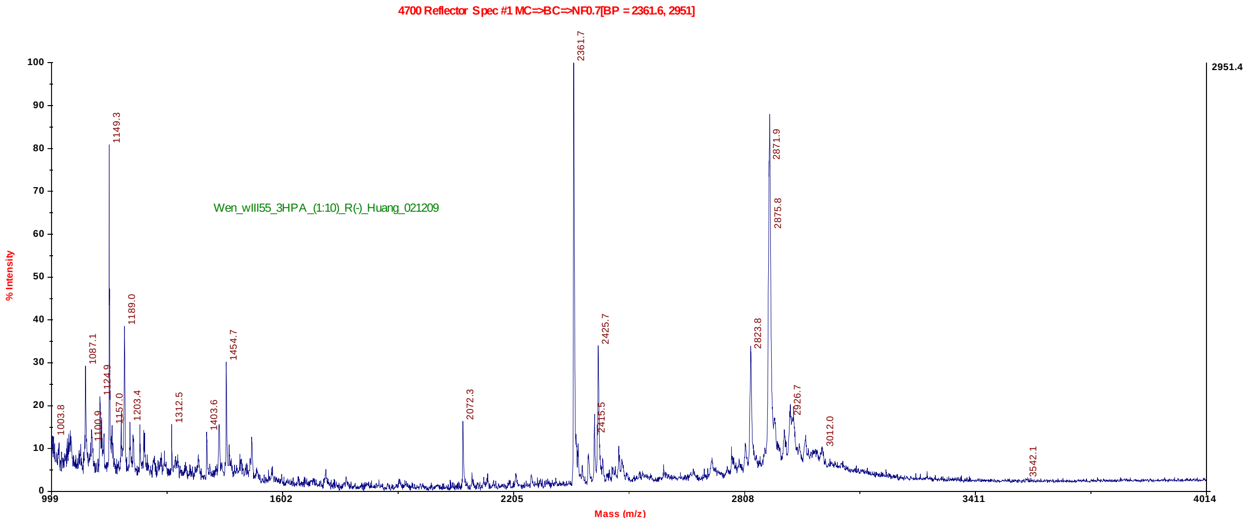


**Figure S40.** MALDI-TOF MS spectrum of 5'-ATGG-$^{5\text{-}CH3O}$T-GCTC-3', Molecular formula: $C_{88}H_{112}N_{32}O_{55}P_8$, $[M+Na]^+$: 2871.9 (calc. 2869.7).

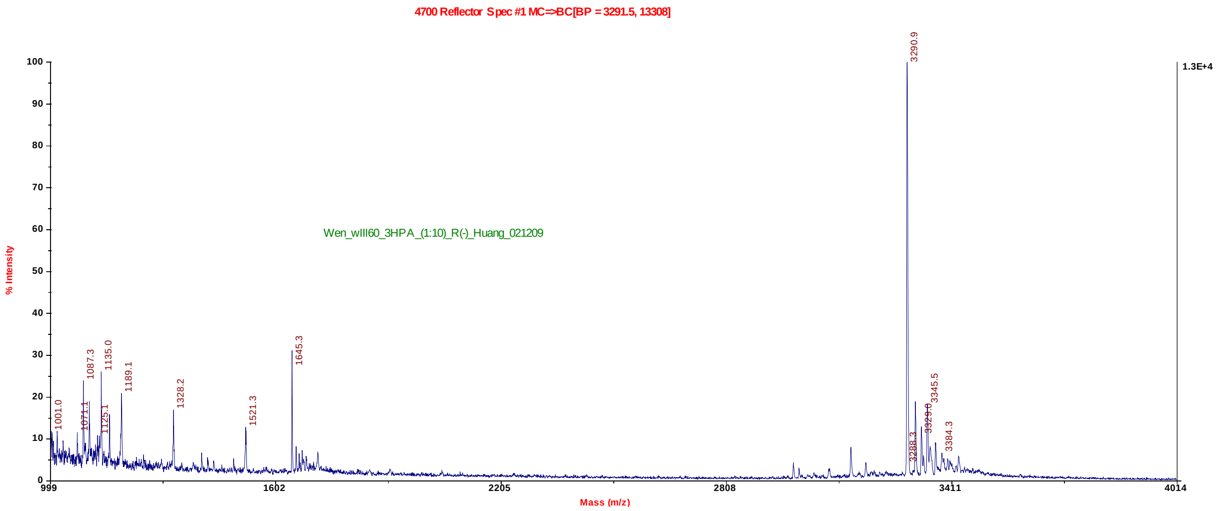


**Figure S41**. MALDI-TOF MS spectrum of 5'-CTTCT-$^{5\text{-}CH3O}$T-GTCCG-3', Molecular formula: $C_{106}H_{138}N_{32}O_{70}P_{10}$, $[M+H]^+$: 3290.9(calc. 3291.3).

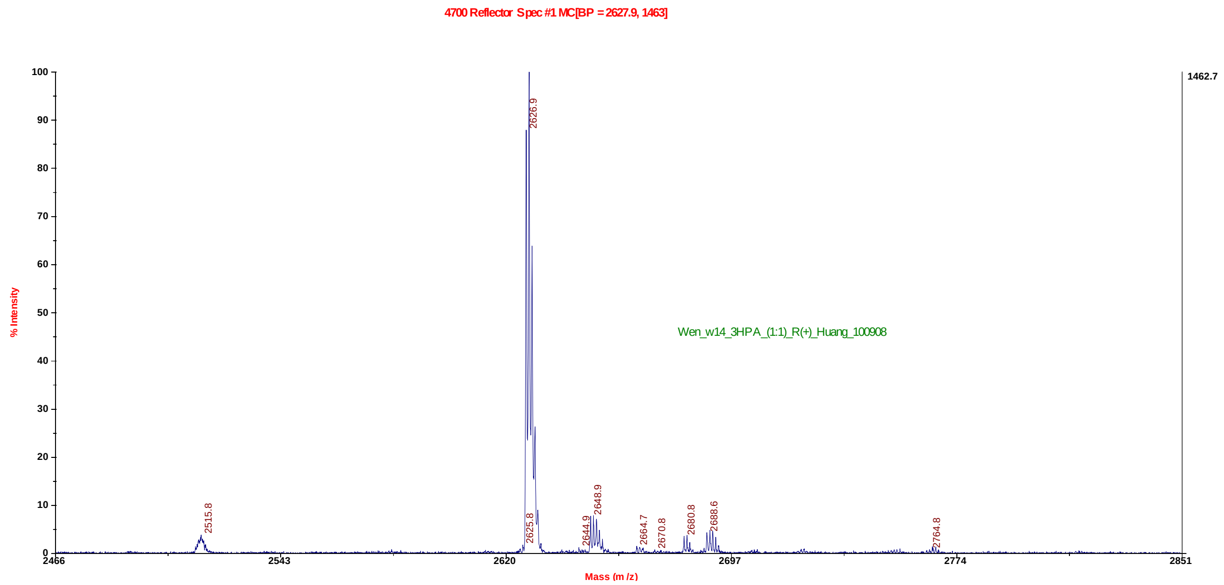


**Figure S42**. MALDI-TOF MS spectrum of 5'-GdU$_{2'\text{-Se}}$G-$^{5\text{-PhSe}}$T-ACAC-3', Molecular formula: $C_{83}H_{101}N_{30}O_{46}P_7Se_2$, $[M]^+$: 2627.9(calc. 2629.6).

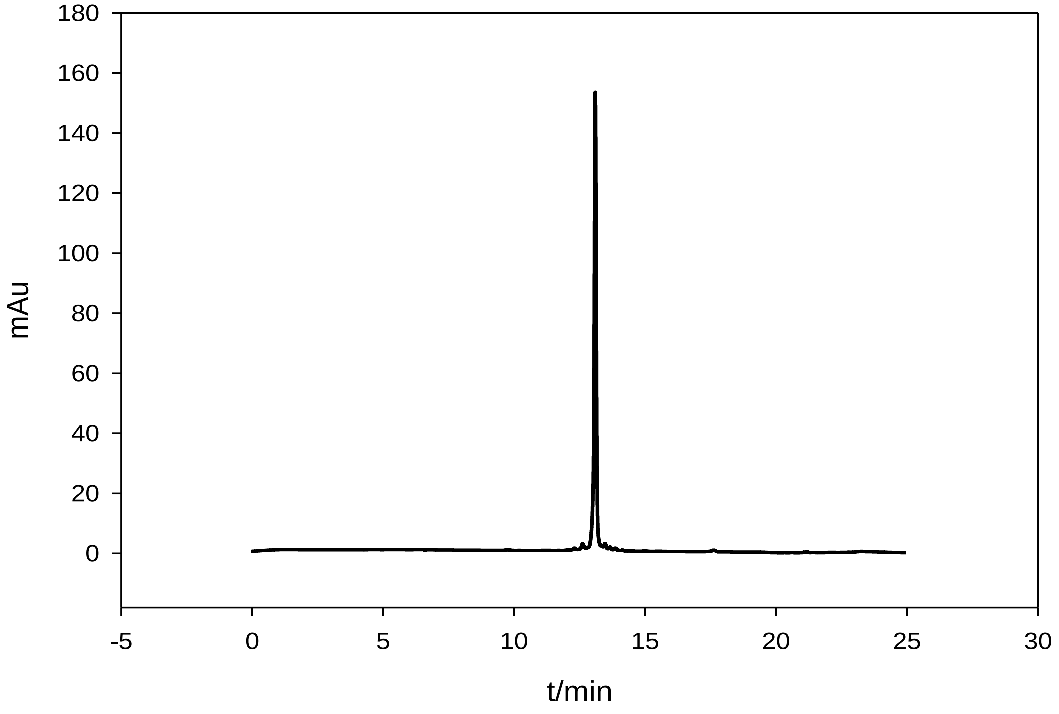


**Figure S43**. HPLC analysis of DNA (5'-CTTCT$^{5\text{-CH3O}}$TGTCCG-3', retention time= 13.6 min)

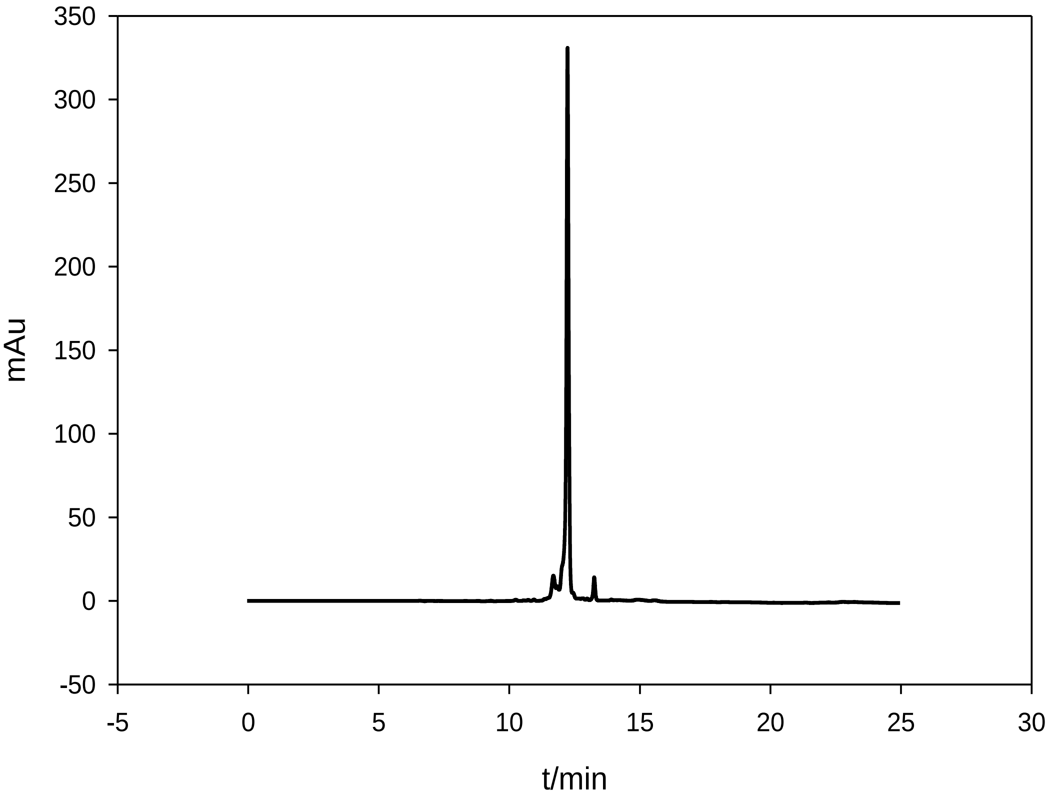


**Figure S44**. HPLC analysis of DNA (5'-CTCCCA-$^{5\text{-}CH3O}$$\underline{T}$-CC-3', retention time= 12.4 min)

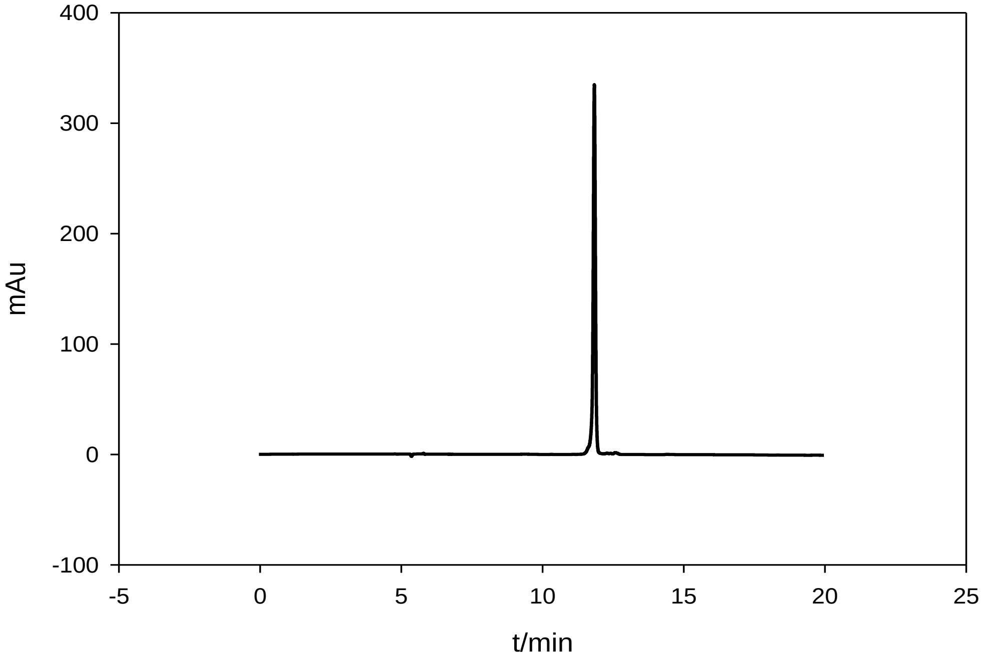


**Figure S45.** HPLC analysis of DNA (5'-CTCCCA-$^{5\text{-}CH3S}$$\underline{T}$-CC-3', retention time= 12.1 min)

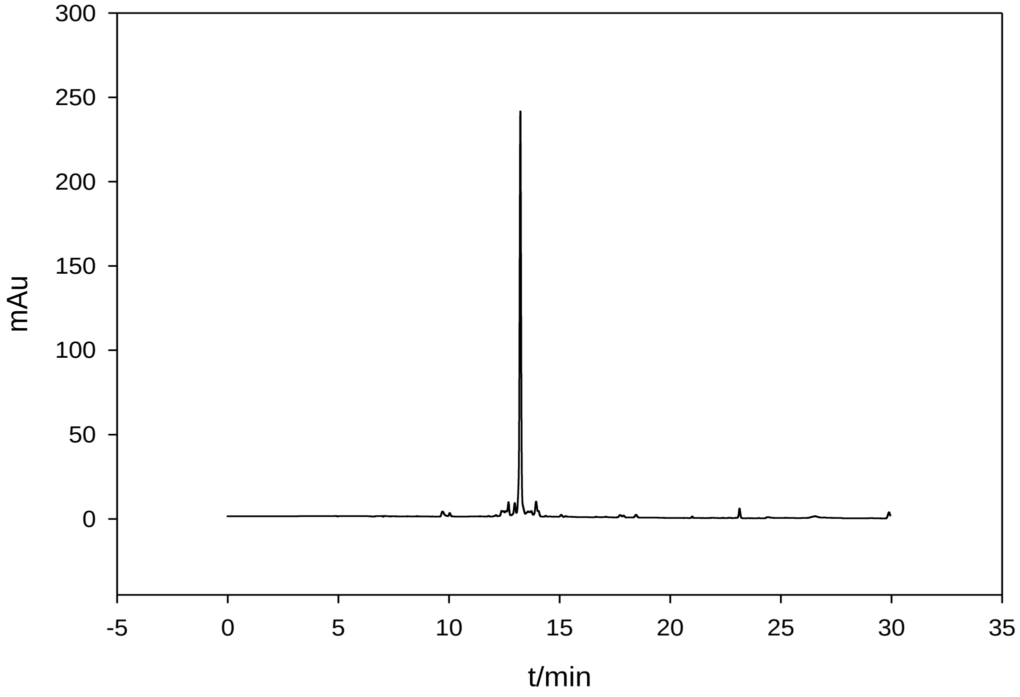


**Figure S46**. HPLC analysis of DNA (5'-CTTCT$^{5\text{-}CH3S}$T̲GTCCG-3', retention time= 13.9 min)

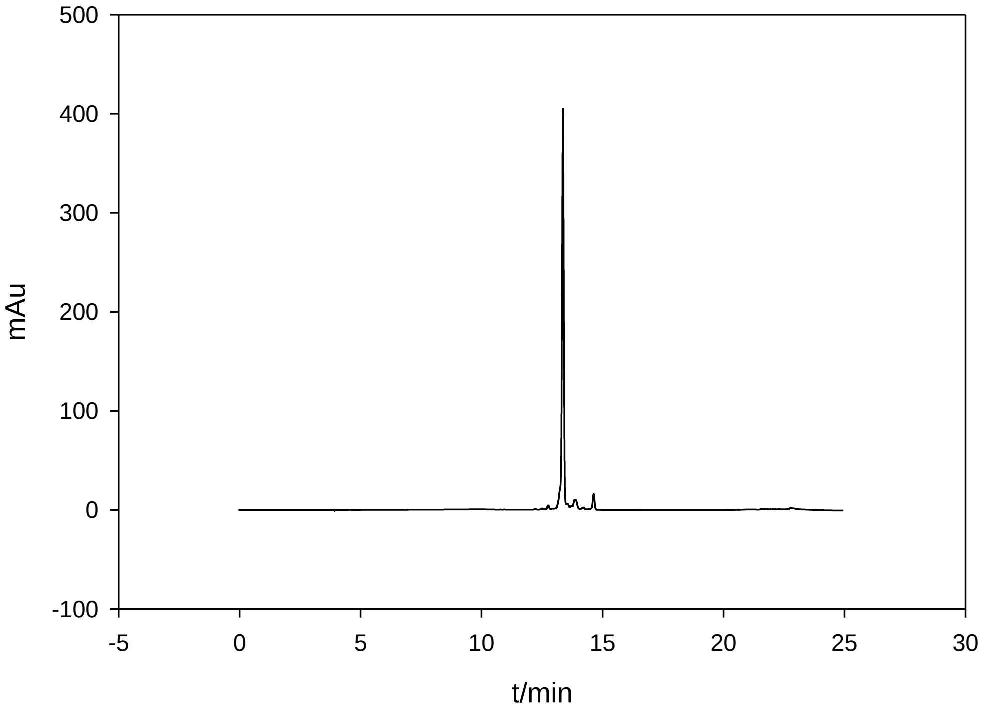


**Figure S47**. HPLC analysis of DNA (5'-CTTCT$^{5\text{-}PhSe}$T̲GTCCG-3', retention time= 13.9 min)

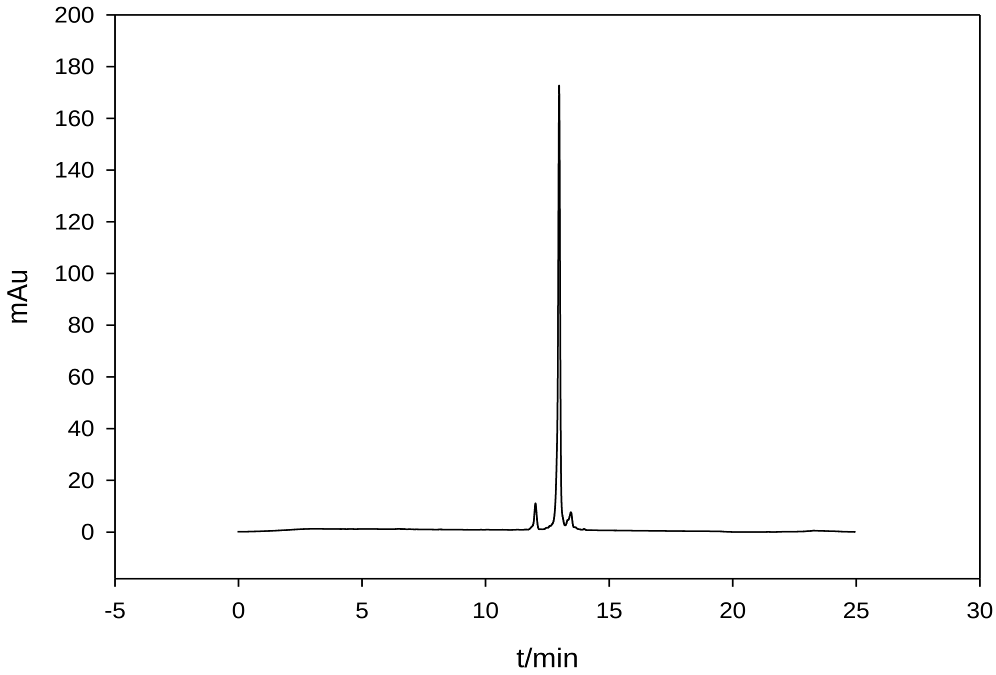

**Figure S48**. HPLC analysis of DNA (5'-CTTCT$^{5\text{-}CH3Se}$<u>T</u>GTCCG-3', retention time= 13.4 min)

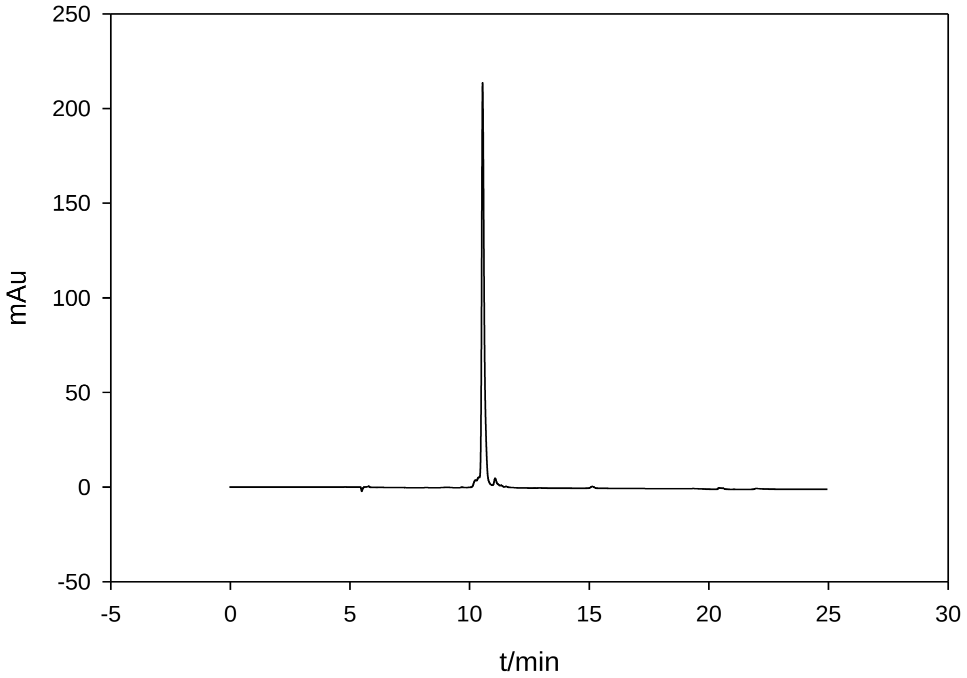

**Figure S49.** HPLC analysis of DNA (5'-ATGG-$^{5\text{-}CH3S}$<u>T</u>-GCTC-3', retention time= 10.8 min)

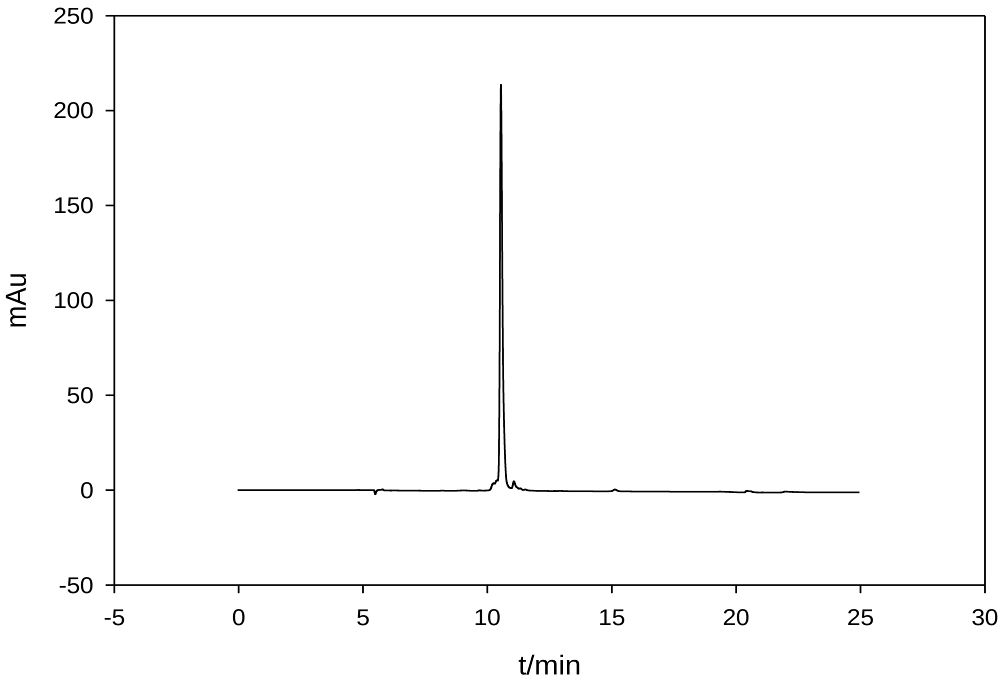


**Figure S50**. HPLC analysis of DNA (5'-ATGG-$^{5\text{-}CH3O}$T-GCTC-3', retention time= 10.3 min)

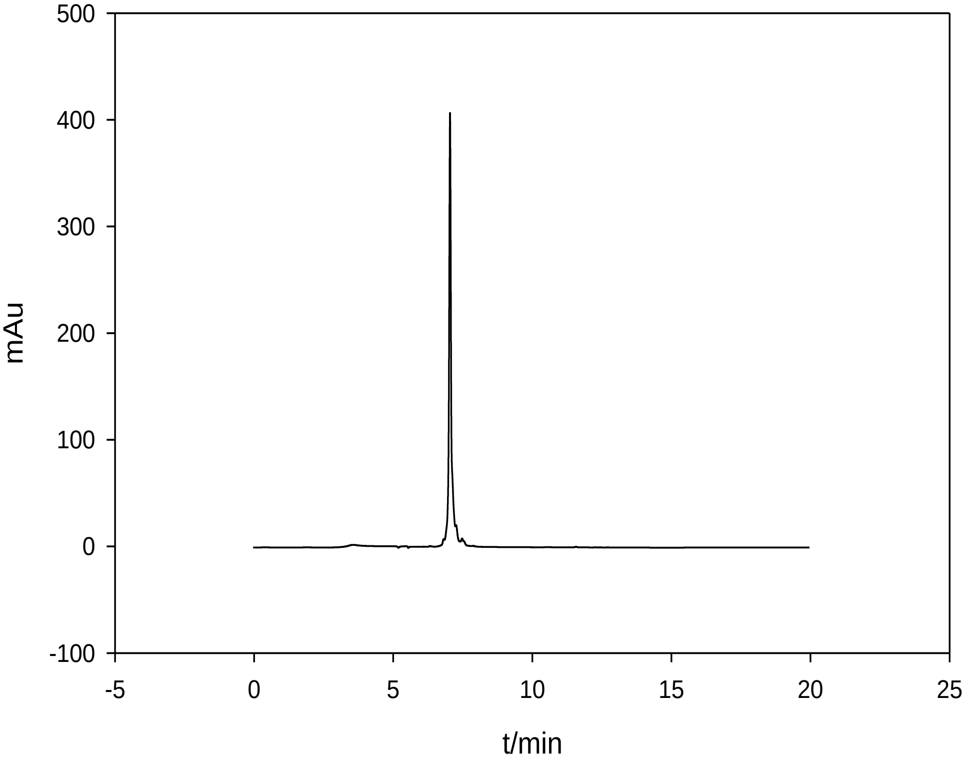


**Figure S51**. HPLC analysis of DNA (5'-GdU$_{2'\text{-}Se}$G-$^{5\text{-}CH3Se}$T-ACAC-3', retention time= 7.2 min)

#### 4. UV-melting temperature experiments

The experiments were performed using the samples (2 µM DNA duplexes) dissolved in the buffer of 50 mM NaCl, 10 mM $NaH_2PO_4$-$Na_2HPO_4$ (pH 6.3), 0.1 mM EDTA, and 10 mM $MgCl_2$. These DNA samples were heated to 85 ºC and allowed to cool down to 5 ºC slowly. These experiments were carried out by Cary 300 UV-Visible Spectrophotometer with a temperature controller at a heating rate of 0.5 ºC/min. The thermal dynamic parameters were calculated through curve fitting by Meltwin 3.5 (http://www.meltwin.com). Typical UV-denaturing curves are shown here.

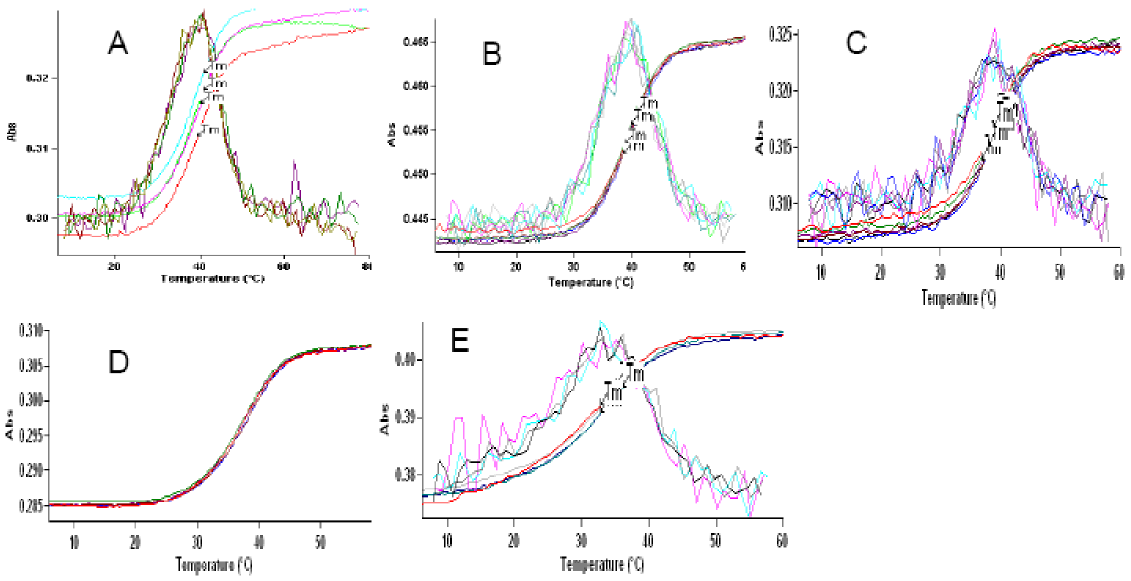


**Figure S52**. UV melting curve of DNA 5'-ATGGXGCTC-3'/5'-GAGCACCAT-3' duplex. (A: X=T; B: X=$^{5CH3Se}$T; C: X=$^{5CH3S}$T; D: X=$^{5CH3O}$T; E: X=$^{5PhSe}$T)

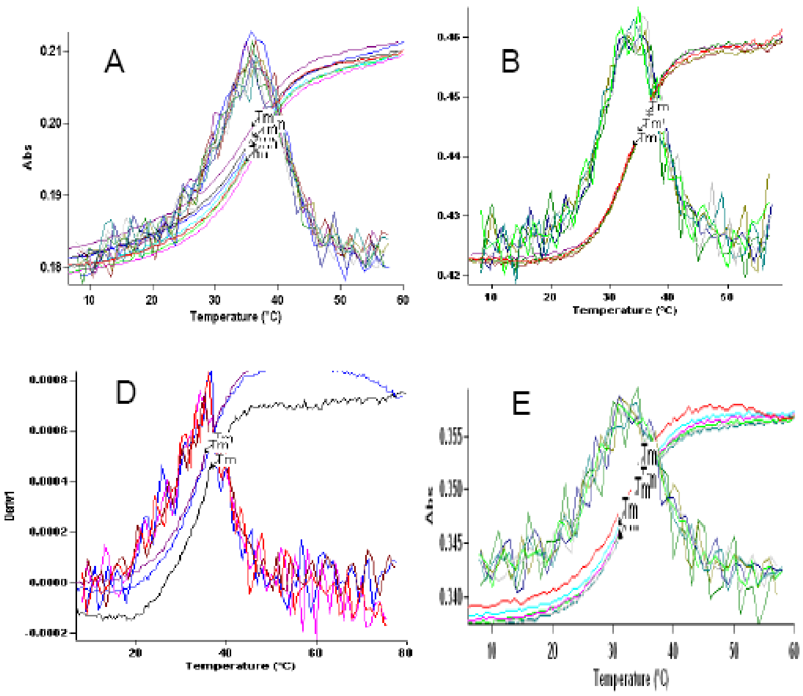


**Figure S53**. UV melting curve of DNA 5'-CTCCCAXCC-3'/5'-GGATGGGAG-3' duplex. (A: X=T; B: X=$^{5CH3Se}$T; C: X=$^{5CH3S}$T; D: X=$^{5CH3O}$T; E: X=$^{5PhSe}$T)

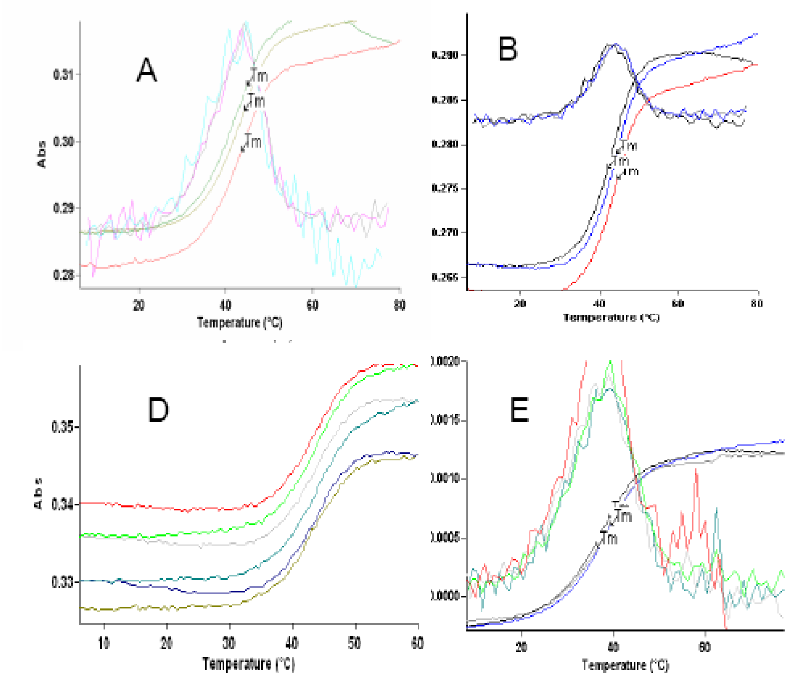


**Figure S54**. UV melting curve of DNA 5'-CTTCTXGTCCG-3'/5'-CGGACAAGAAG-3'. (A: X=T; B: X=$^{5CH3Se}$T; C: X=$^{5CH3S}$T; D: X=$^{5CH3O}$T; E: X=$^{5PhSe}$T)

#### 4. Nuclease stability experiment

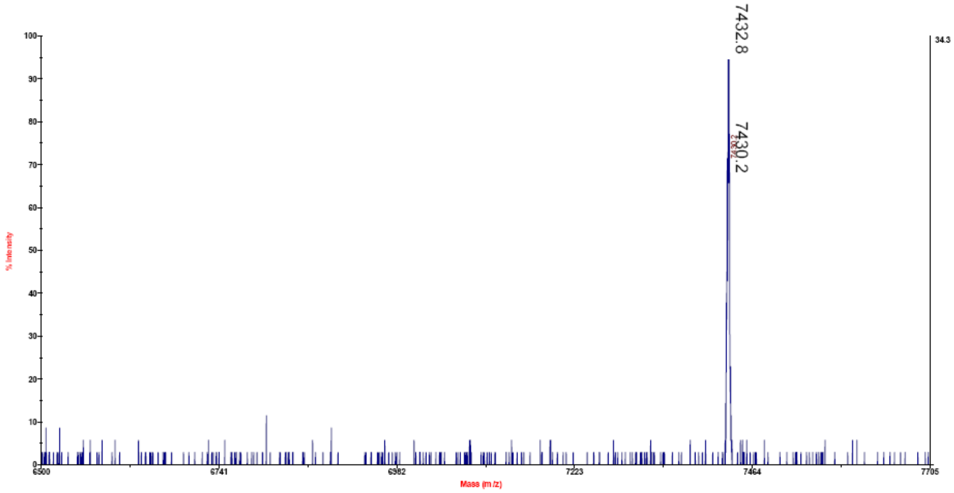


**Figure S55**. MALDI-TOF MS spectrum of DNA2 5'-GTGCACTGATCAA$^{5\text{-MeSe}}$TTAATGTCGAC-3'. Molecular formula: $C_{235}H_{296}N_{89}O_{142}P_{23}Se$; $[M]^+$: 7430.2 (calc. 7430.9), $[M+2H]^+$: 7432.9 (calc. 7432.8).

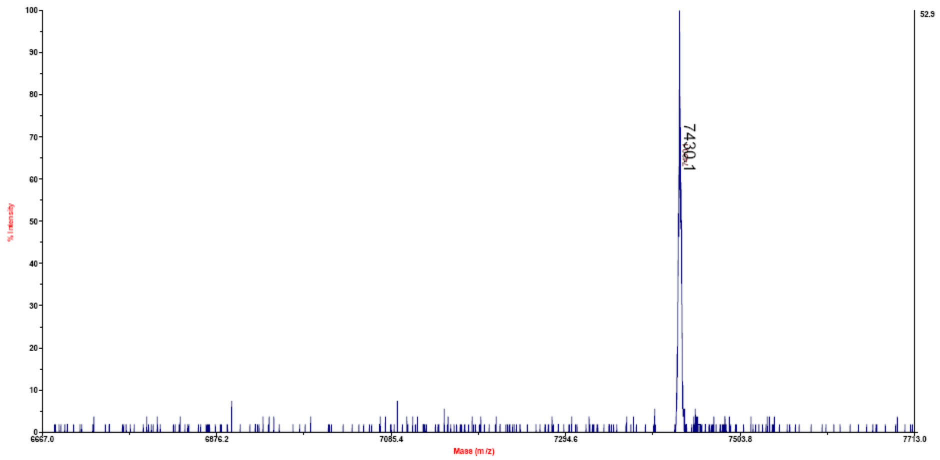


**Figure S56**. MALDI-MS spectrum of DNA3 5'-GTGCACTGATCAAT$^{5\text{-MeSe}}$TAATGTCGAC-3'. Molecular formula: $C_{235}H_{296}N_{89}O_{142}P_{23}Se$; $[M]^+$: 7430.1 (calc. 7430.9).

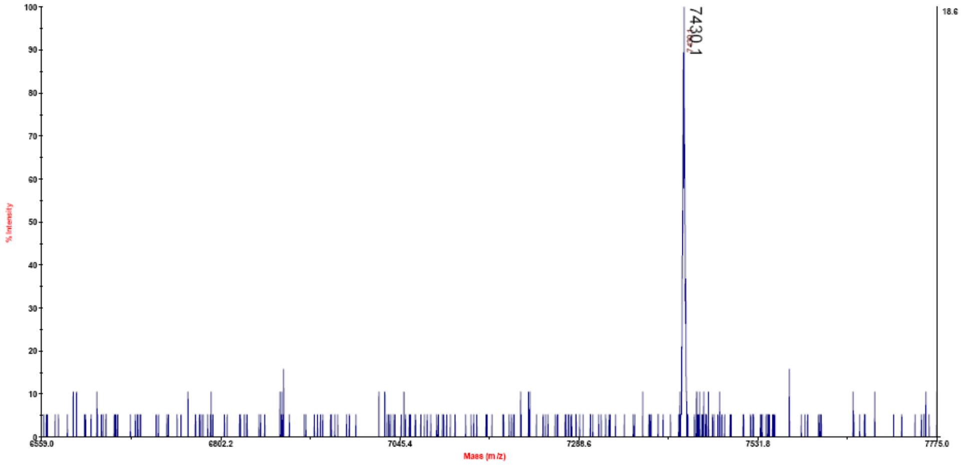


**Figure S57**. MALDI-MS spectrum of DNA4 5'-GTGCACTGATCAATTAA$^{5\text{-MeSe}}$<u>T</u>GTCGAC-3'. Molecular formula: $C_{235}H_{296}N_{89}O_{142}P_{23}Se$; $[M]^+$: 7430.1 (calc. 7430.9).

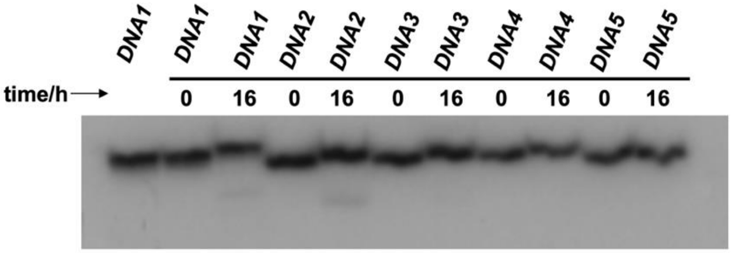


**Figure S58:** Enzymatic stabilities of $^{5\text{-CH3Se}}$T DNAs to endonuclease SalI.

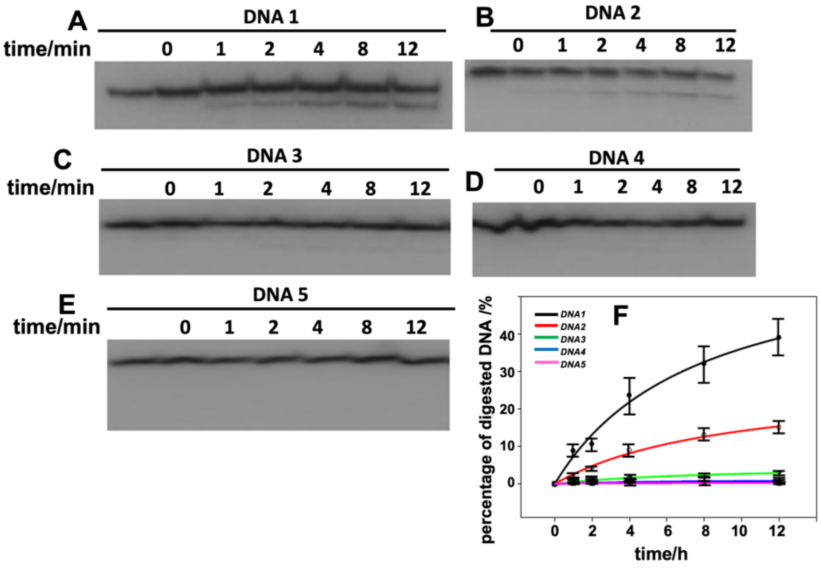


**Figure S59**: Endonuclease SalI digestion of DNAs containing $^{5\text{-CH3Se}}$T. (A) native DNA 1; (B) DNA 2; (C) DNA 3; (D) DNA 4; (E) DNA5; (F) Plot of the ratio of DNA digested over time.

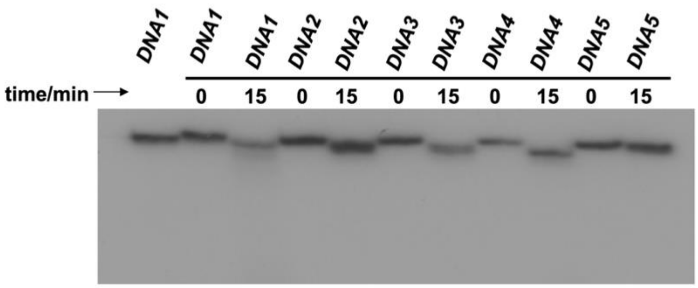


**Figure S60:** Enzymatic stabilities of 5-CH3SeT DNAs to exonuclease III.

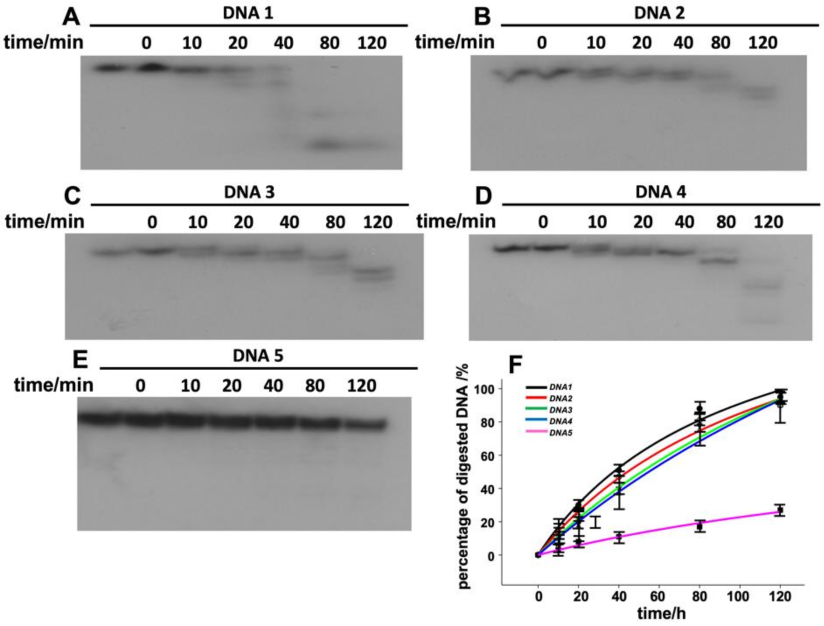


**Figure S61**: Exonuclease III digestion of DNAs containing 5-CH3SeT. (A) native DNA 1; (B) DNA 2; (C) DNA 3; (D) DNA 4; (E) DNA5; (F) Plot of the ratio of DNA digested over time.

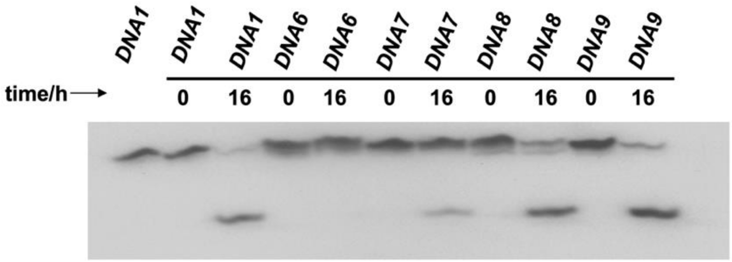


**Figure S62:** Enzymatic stabilities of 5-CH3ST DNAs to endonuclease AseI.

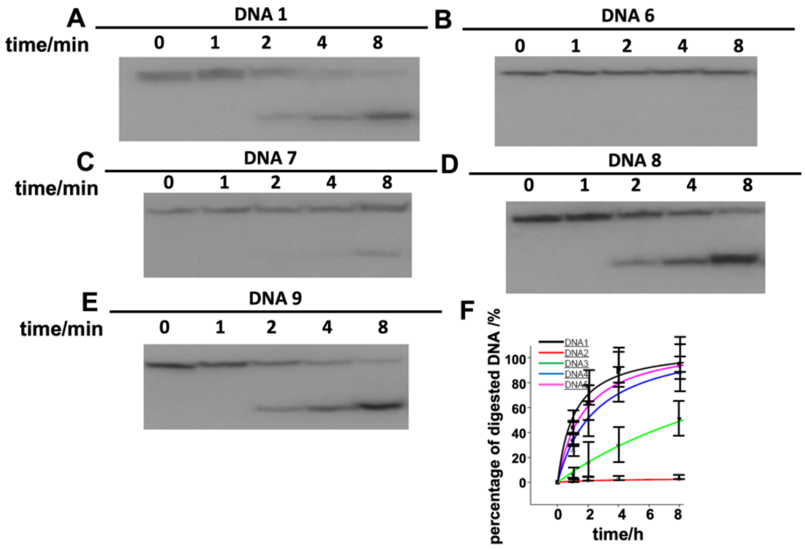


**Figure S63**: Endonuclease AseI digestion of DNAs containing $^{5\text{-CH3S}}$T. (A) native DNA 1; (B) DNA 6; (C) DNA 7; (D) DNA 8; (E) DNA9; (F) Plot of the ratio of DNA digested over time.

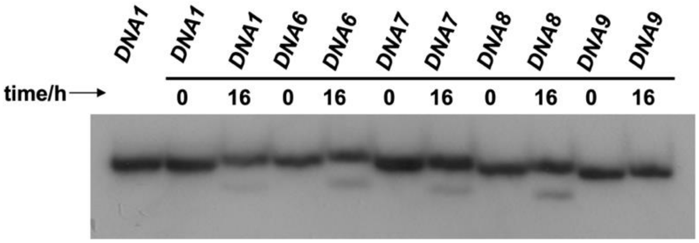


**Figure S64:** Enzymatic stabilities of $^{5\text{-CH3S}}$T DNAs to endonuclease SalI.

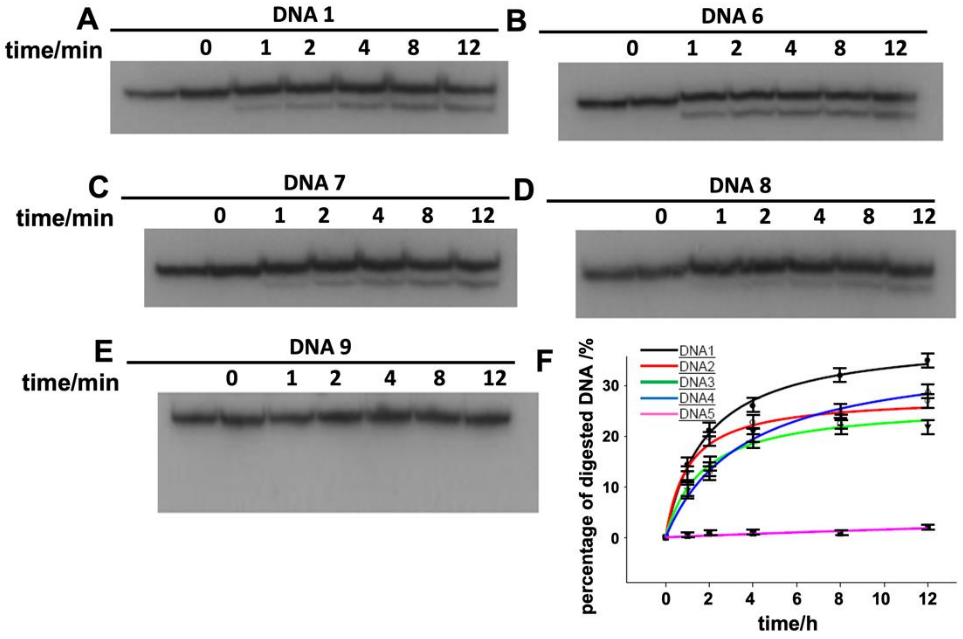


**Figure S65**: Endonuclease SalI digestion of DNAs containing $^{5\text{-}CH3S}$T. (A) native DNA 1; (B) DNA 6; (C) DNA 7; (D) DNA 8; (E) DNA9; (F) Plot of the ratio of DNA digested over time.

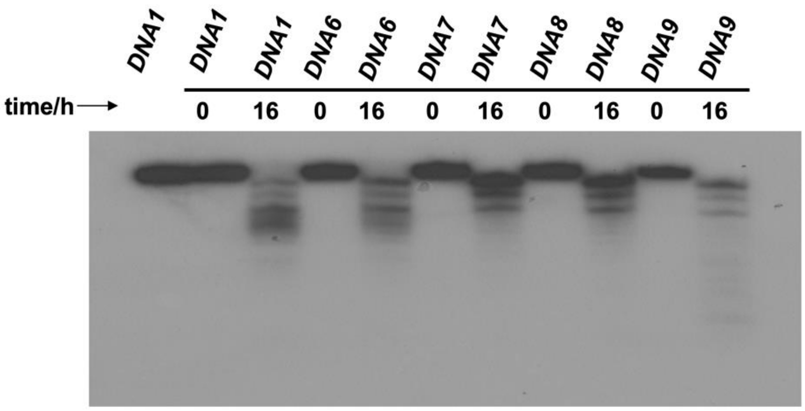


**Figure S66:** Enzymatic stabilities of $^{5\text{-}CH3S}$T DNAs to exonuclease III.

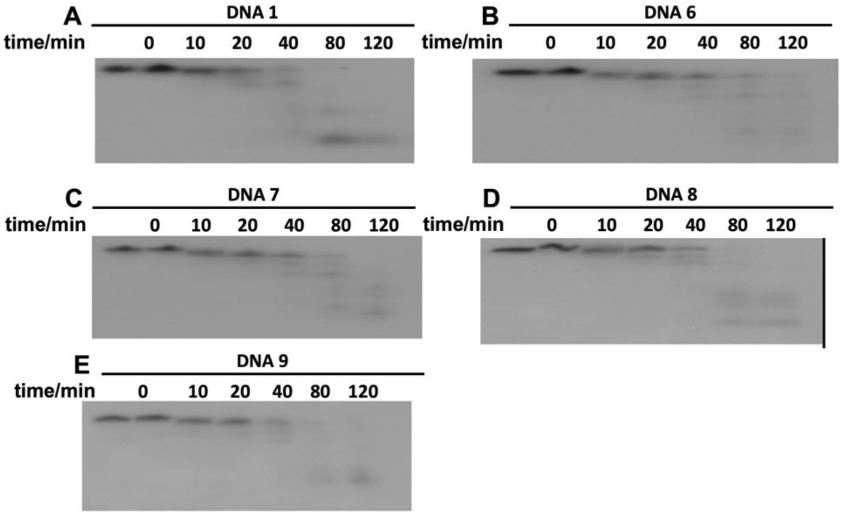


**Figure S67**: Exonuclease III digestion of DNAs containing $^{5\text{-}CH3S}$T. (A) native DNA 1; (B) DNA 6; (C) DNA 7; (D) DNA 8; (E) DNA9; (F) Plot of the ratio of DNA digested over time.

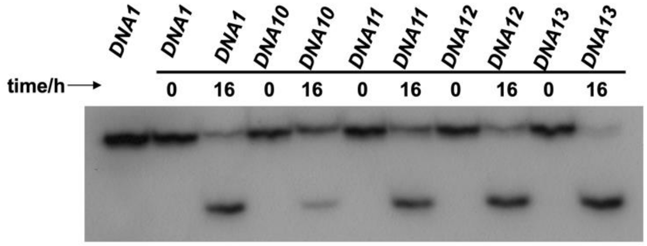


**Figure S68:** Enzymatic stabilities of $^{5\text{-}CH3O}$T DNAs to endonuclease AseI.

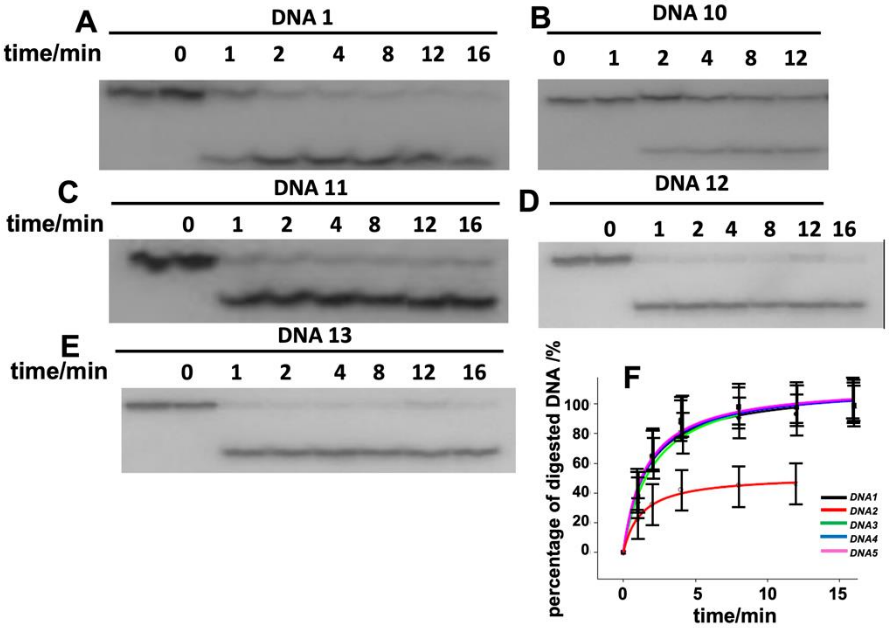


**Figure S69**: Endonuclease AseI digestion of DNAs containing $^{5\text{-}CH3O}$T. (A) native DNA 1; (B) DNA 10; (C) DNA 11; (D) DNA 12; (E) DNA13; (F) Plot of the ratio of DNA digested over time.

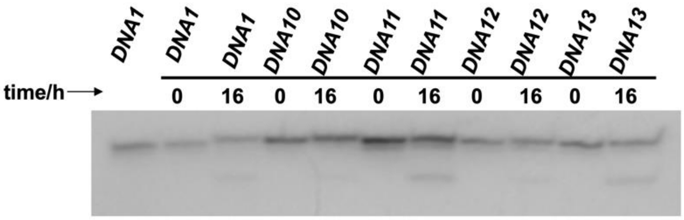


**Figure S70:** Enzymatic stabilities of $^{5\text{-}CH3O}$T DNAs to endonuclease SalI.

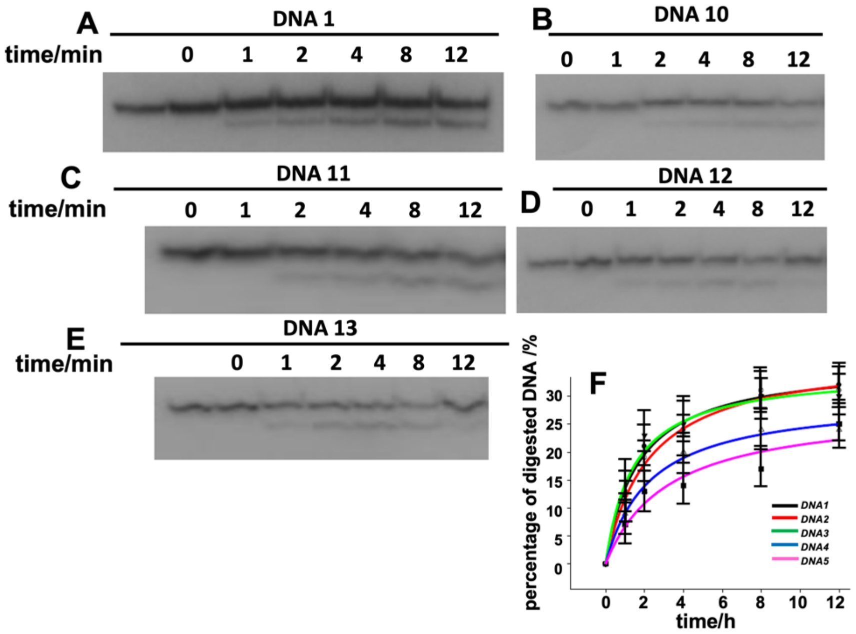


**Figure S71**: Endonuclease SalI digestion of DNAs containing $^{5\text{-}CH3O}$T. (A) native DNA 1; (B) DNA 10; (C) DNA 11; (D) DNA 12; (E) DNA13; (F) Plot of the ratio of DNA digested over time.

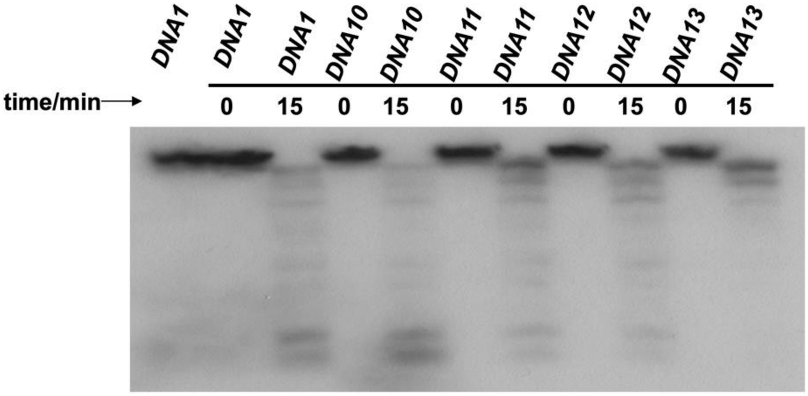


**Figure S72:** Enzymatic stabilities of $^{5\text{-}CH3O}$T DNAs to exonuclease III.

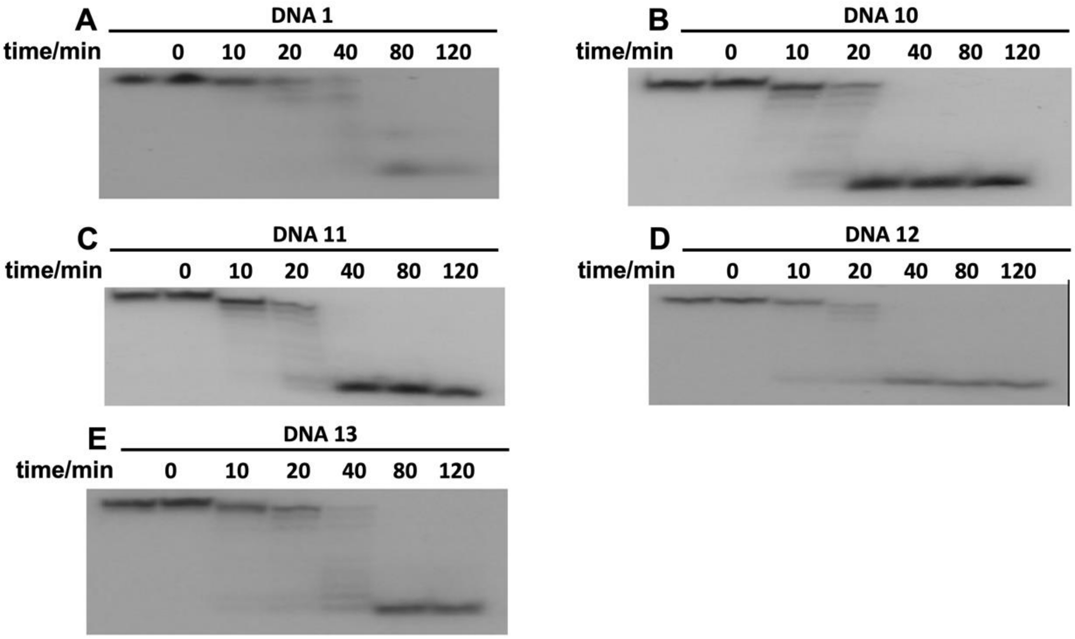


**Figure S73**: Exonuclease III digestion of DNAs containing $^{5\text{-}CH3O}$T. (A) native DNA 1; (B) DNA 10; (C) DNA 11; (D) DNA 12; (E) DNA13; (F) Plot of the ratio of DNA digested over time.

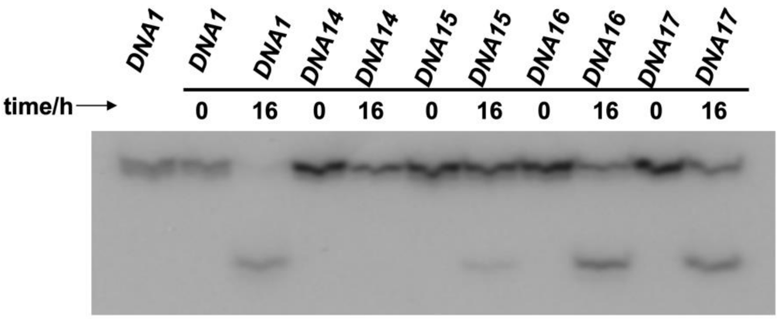


**Figure S74:** Enzymatic stabilities of $^{5\text{-}PhSe}$T DNAs to endonuclease AseI.

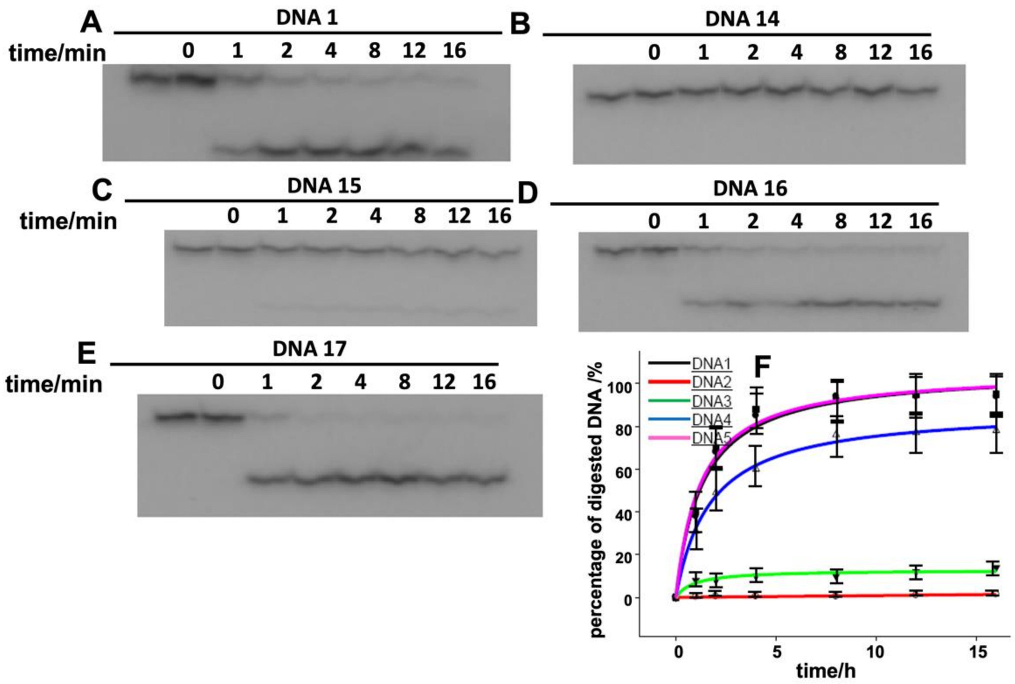


**Figure S75**: Endonuclease AseI digestion of DNAs containing $^{5\text{-PhSe}}$T. (A) native DNA 1; (B) DNA 14; (C) DNA 15; (D) DNA 16; (E) DNA17; (F) Plot of the ratio of DNA digested over time.

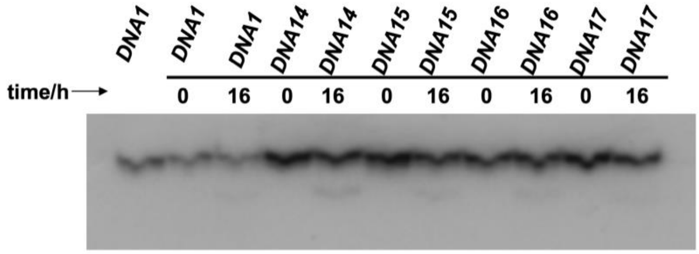


**Figure S76:** Enzymatic stabilities of $^{5\text{-PhSe}}$T DNAs to endonuclease SalI.

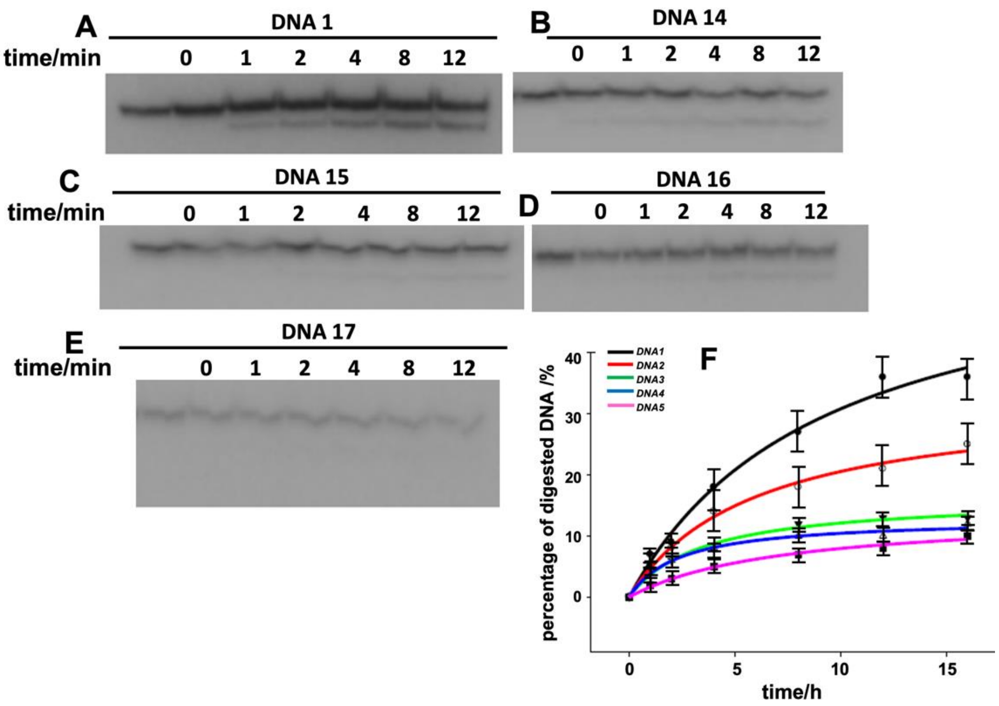


**Figure S77**: Endonuclease SalI digestion of DNAs containing $^{5\text{-PhSe}}$T. (A) native DNA 1; (B) DNA 14; (C) DNA 15; (D) DNA 16; (E) DNA17; (F) Plot of the ratio of DNA digested over time.

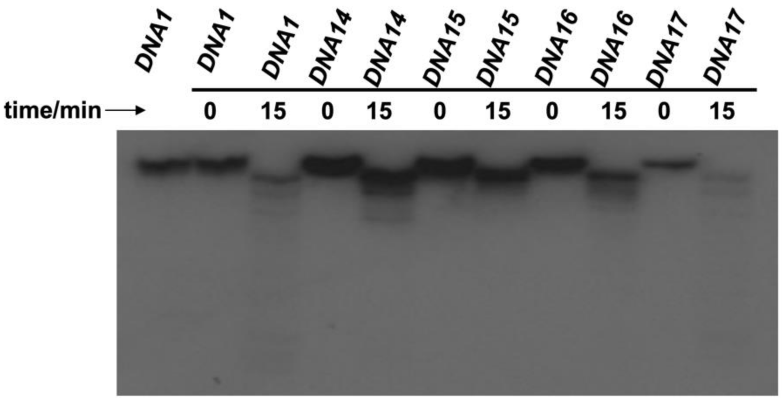


**Figure S78:** Enzymatic stabilities of $^{5\text{-PhSe}}$T DNAs to exonuclease III.

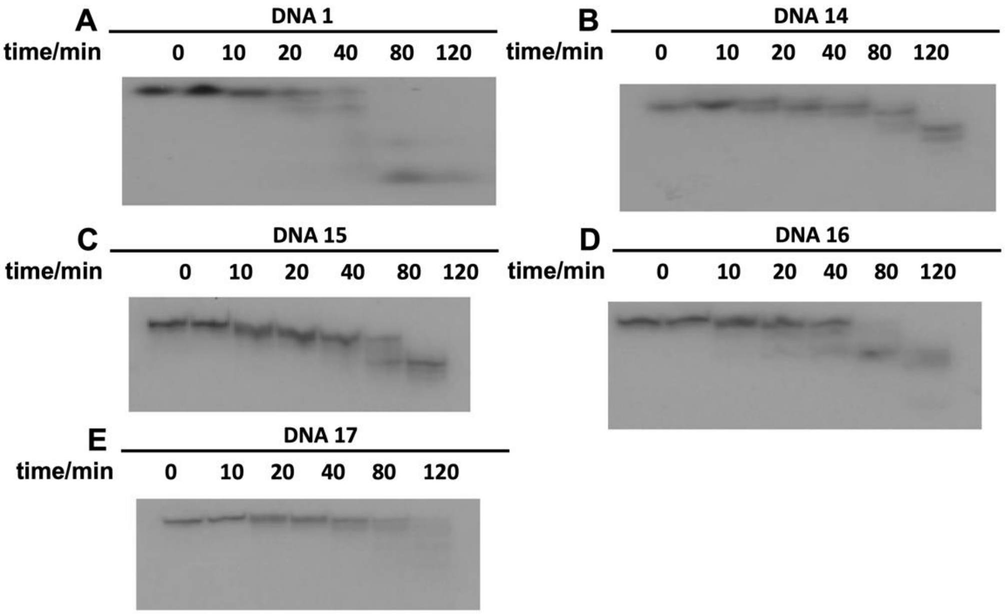


**Figure S79**: Exonuclease III digestion of DNAs containing $^{5\text{-PhSe}}$T. (A) native DNA 1; (B) DNA 14; (C) DNA 15; (D) DNA 16; (E) DNA17; (F) Plot of the ratio of DNA digested over time.

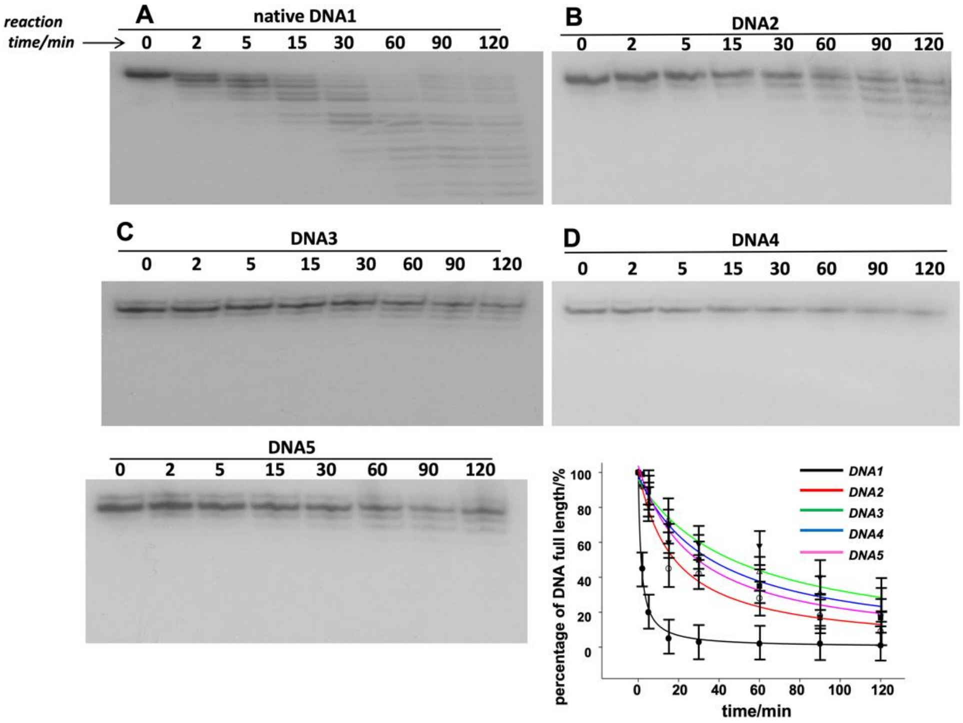


**Figure S80**: Snake venom phosphodiesterase digestion of DNAs containing $^{5\text{-CH3Se}}$T. (A) native DNA 1; (B) DNA 2; (C) DNA 3; (D) DNA 4; (E) DNA5; (F) Plot of the ratio of DNA digested over time.

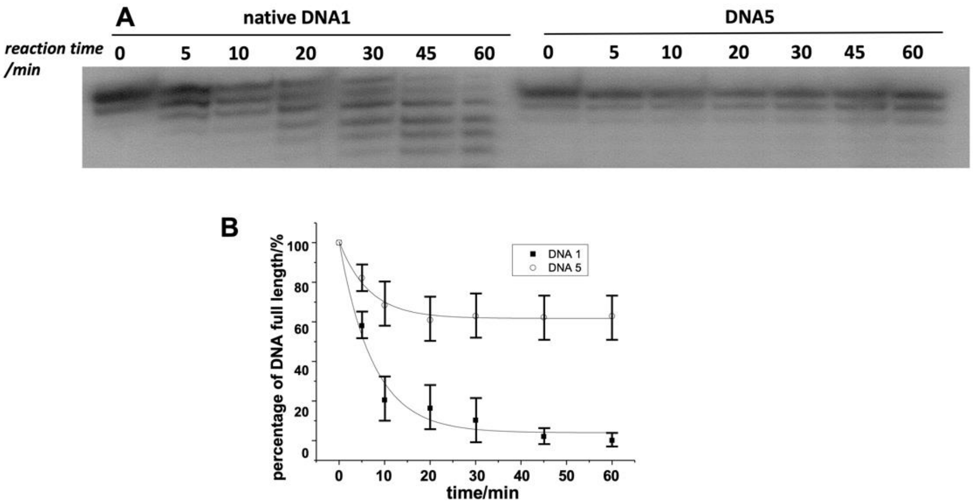


**Figure S81**: Serum digestion of native DNA and DNA5 containing $^{5\text{-}CH3Se}$T. Bottom: Plot of the ratio of DNA digested over time.

#### 5. MS spectrum of Se-antisense to suppress EGFP gene expression in HeLa cells

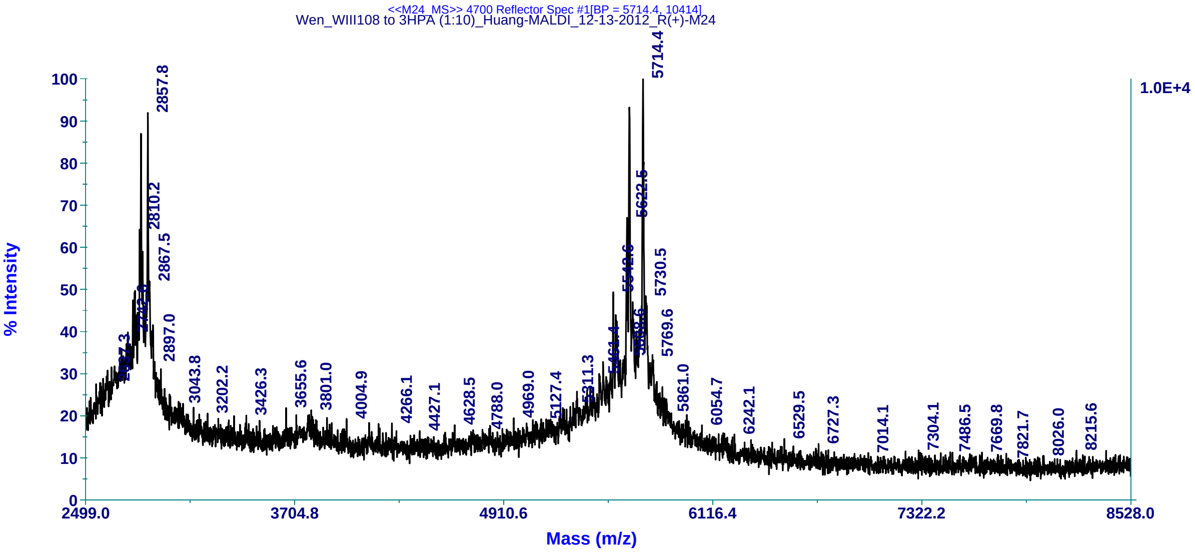


**Figure S82**. MALDI-MS spectrum of DNAe 5'-GAGC$^{5\text{-}CH3Se}$<u>T</u>GCACGC$^{5\text{-}CH3Se}$<u>T</u>GCCG$^{5\text{-}CH3Se}$<u>T</u>C-3'. Molecular formula: $C_{235}H_{296}N_{89}O_{142}P_{23}Se_3$; $[M+H]^+$: 5714.4 (calc. 5714.6).

#### 6. X-ray Crystallography statistics

**Crystal preparation.** QIAGEN Classics Suite Kit were used for screening crystallization conditions by the hanging drop vapor diffusion method. Solutions containing the DNA/RNA duplex sample (0.5 mM each) were heated to 90 °C for 1 min and cooled slowly to room temperature, then mixed with RNase H (6 mg/mL) at 1:1 ratio. All of the crystals grew at 20 °C, and the mother liquor containing 25% glycerol was used as a cryoprotectant during crystal mounting. All data collection was taken under a stream of nitrogen at 99 K. The data sets were collected at the SIBYLS beamline 501 and 822 at the Advanced Light Source, Lawrence Berkeley National Laboratory. The distances between the detector and the crystal were set to 300 mm and the collecting wavelength was set to 1.00 Å. The crystals were exposed for 1 second per image with one degree oscillations, and 180 images were taken for each data set. The optimized crystallization conditions for the RNase complex is #96 of the crystallization screen [Buffer: 0.1 M MES, pH 6.5; precipitant: 12% (w/v), PEG 20000].

**Data collection and structure refinement.** The data were processed using HKL2000 and CCP4. The structures were solved by molecular replacement using PDB 2G8U as model. The resulted model was refined using Refmac5 within CCP4i. The modified DNA was modeled into the structure using Coot. After several cycles of refinement, a number of highly ordered water molecules and metal ions were added either automatically or manually using Coot. Data collection, phasing, and refinement statistics of the determined structures are listed in Table S2.

**Table S2.** Data Collection Statistics.

| **Structure** | RNase H /RNA/DNA | RNase H /RNA/Se-DNA | RNase H /RNA/Se-DNA | RNase H /RNA/Se-DNA |
|---|---|---|---|---|
| DNA sequence | 5'-ATGTCG-3' | 5'-A$^{se}\underline{T}$GTCG-3' | 5'-ATG$^{se}\underline{T}$CG-3' | 5'-A$^{se}\underline{T}$G$^{se}\underline{T}$CG-3' |
| Space group | C121 | C121 | C121 | C121 |
| Unit cell parameters (Å, $^{o}$) | 80.3, 37.6, 62.1<br>90, 95.9, 90 | 81.2, 37.7, 62.2<br>90, 96.4, 90 | 80.9, 37.6, 62.3<br>90, 96.2, 90 | 80.9, 37.9, 62.1<br>90, 96.2, 90 |
| Resolution range, Å (last shell) | 61.79-1.70 (1.76-1.70) | 50.00-1.80 | 50.00-1.73 | 50.00-1.70 |
| Unique reflections | 20483 | 17472 | 190601 | 20662 |
| Completeness, % | 92.1 (53.7) | 100.0 | 99.8 | 100.0 |
| $R_{merge}$, % | 3.3 (53.7) | 2.5 | 3.9 | 2.3 |
| <I/σ(I)> | 21.1 (3.5) | 29.0 | 53.7 | 31.5 |
| Redundancy | 5.8 (3.3) | 7.3 | 6 | 7.3 |

| **Structure** | 5-MeO-DNA | 5-MeS-DNA | 5-MeSe-DNA |
|---|---|---|---|
| DNA sequence | 5'-G$_{2'\text{-}Se}$TG$^{5\text{-}MeO}$ $\underline{T}$ACAC-3' | 5'-G$_{2'\text{-}Se}$TG$^{5\text{-}MeS}\underline{T}$ACAC-3' | 5'-G$_{2'\text{-}Se}$TG$^{5\text{-}MeSe}\underline{T}$ACAC-3' |
| Space group | P43212 | P43212 | P43212 |
| Unit cell parameters (Å, $^{o}$) | 43.2, 43.2, 23.7<br>90, 90, 90 | 42.9, 42.9, 23.7<br>90, 90, 90 | 43.1, 43.1, 23.8<br>90, 90, 90 |
| Resolution range, Å (last shell) | 50.00-1.30 (1.32-1.30) | 50.00-1.38 (1.43-1.38) | 50.00-1.40 (1.45-1.40) |
| Unique reflections | 5561(264) | 4616 | 4661 (446) |
| Completeness, % | 94.5 (50.8) | 94.0 (67.8) | 97.8 (99.3) |
| $R_{merge}$, % | 3.7 (68.2) | 6.3 (24) | 8.7 (13.6) |
| <I/σ(I)> | 38.4 (1.0) | 23.0 | 18.9 (14.5) |
| Redundancy | 12.7 (2.0) | 8.2 (5.2) | 23.0 (11.1) |

**Table S3.** Structure Refinement Statistics.

| **Structure** | RNase H /RNA/DNA | RNase H /RNA/Se-DNA | RNase H /RNA/Se-DNA | RNase H /RNA/Se-DNA |
|---|---|---|---|---|
| PDB code | 5WJR | 5USA | 5USE | 5USG |
| Molecules per asymmetric unit | 1 | 1 | 1 | 1 |
| Resolution range, Å | 61.79-1.70 | 35.66-1.80 | 30.95-1.73 | 35.51-1.70 |
| $R_{work}$, % | 18.9 | 19.0 | 19.2 | 17.8 |
| $R_{free}$, % | 23.4 | 21.6 | 23.4 | 22.1 |
| Number of reflections | 17002 | 16605 | 18567 | 19632 |
| Completeness for Rrange, % | 87.3 | 99.8 | 99.7 | 99.9 |
| Bond length R.M.S., Å | 0.019 | 0.018 | 0.021 | 0.021 |
| Bond angle R.M.S. | 1.849 | 1.905 | 2.039 | 2.128 |
| Average B-factors, Å$^2$ | 15.65 | 24.30 | 27.80 | 22.11 |

| **Structure** | 5-MeO-DNA | 5-MeS-DNA | 5-MeSe-DNA |
|---|---|---|---|
| PDB code | 3IJK | 3HG8 | 3LTU |
| duplex per asymmetric unit | 0.5 | 0.5 | 0.5 |
| Resolution range, Å | 30.56-1.30 | 20.73-1.38 | 30.46-1.40 |
| $R_{work}$, % | 19.7 | 18.7 | 17.5 |
| $R_{free}$, % | 21.9 | 19.2 | 20.0 |
| Number of reflections | 5301 | 4373 | 4421 |
| Completeness for Rrange, % | 95.1 | 94.1 | 97.8 |
| Bond length R.M.S., Å | 0.006 | 0.007 | 0.004 |
| Bond angle R.M.S. | 1.600 | 3.116 | 1.108 |
| Average B-factors, Å$^2$ | 12.79 | 12.43 | 8.96 |
